# Parallels in the Regulatory Landscape of Dimorphic Female and Male Genital Structures in *Drosophila melanogaster*

**DOI:** 10.1101/2025.05.12.653573

**Authors:** Eden McQueen, Gavin Rice, Shanker Pillai, Omid Saleh Ziabari, Ben J Vincent, Mark Rebeiz

## Abstract

Understanding how morphological structures evolve via changes to their development is an ongoing pursuit in biology. Comparative approaches examine changes in the expression or function of key developmental molecules (e.g. transcription factors, signaling molecules or cellular effectors) within homologous structures, and correlate these changes with structural divergence across species, populations, the sexes, or even between different body parts within individuals. The female and male genitalia of *Drosophila* offer an excellent opportunity to investigate homology and trait evolution, as fruit fly genital structures are developmentally tractable and evolve rapidly. While previous work has characterized gene regulatory networks operating in the development and evolution of male genital structures in *Drosophila*, female pupal genitalia are comparatively understudied. Here, we traced the development of female pupal genitalia to determine when and how individual structures form. We then measured the expression patterns of 29 transcription factors in both female and male genital structures at high resolution using hybridization chain reaction and confocal microscopy. We found that these transcription factors are highly patterned in both sexes, and some serve as marker genes for distinct genital structures in females. Our results suggest that the same transcription factors may control developmental processes in female and male genitalia, and this data enables future studies that interrogate how developmental gene regulatory networks specialize and evolve in both sexes.

## Introduction

Demystifying the developmental processes that produce organismal structures and applying that knowledge to understand how those structures evolve is an ongoing pursuit in biology. Comparative approaches are a cornerstone of this field–changes to key molecules (e.g. transcription factors, signaling molecules or cellular effectors) within homologous structures can be correlated with structural divergence across species, populations, the sexes, or even between different body parts within individuals (Smith et al., 2018). This approach works well for many intuitively homologous features such as mammalian digits (Cooper et al., 2014; Lopez-Rios et al., 2014), or serially homologous segments such as vertebrae (Burke et al., 1995) and crustacean limbs (Averof & Patel, 1997). However, homology relationships are not always straightforward to identify. Sexually dimorphic traits, deep homology of long-diverged structures across species, or drastically diversified serial homologs can be challenging to recognize because evolutionary changes can mask underlying similarities (DiFrisco & Jaeger, 2021). Furthermore, the emergence of evolutionary novelties often involves network co-option: the redeployment of a pre-existing gene regulatory network in a new developmental context (E. McQueen & Rebeiz, 2020). This mechanism can produce similar regulatory networks that control the development of nonhomologous structures (DiFrisco & Jaeger, 2021; E. McQueen & Rebeiz, 2020).

Whenever a homology relationship exists between structures that must develop from, and be inherited within, the context of the same genome, (i.e. sexually dimorphic traits, serial homology, network co-option), pleiotropy may influence how these structures evolve (Angelini et al., 2012). Thus, identifying even highly cryptic cases of homology is critical for understanding the evolution of forms. For these types of features, trait-specific molecular markers have been used to assign homology and facilitate evolutionary analysis. In particular, structures and cell types can be defined using expression patterns for genes that serve key functions in development, and shifts in the presence, timing, or spatial patterns of these molecular markers can be compared across groups (Fisher et al., 2020; Hu et al., 2019; Moczek & Rose, 2009; Prud’homme et al., 2011). However, the use of molecular methods to identify cryptic homology relationships and demarcate organismal features is still emerging in the field of evolutionary biology (Rice et al., 2023).

The female and male genitalia of *Drosophila* offer an excellent opportunity to gain new insights into the study of homology and trait evolution using comparative developmental approaches. Due to their rapid diversification, animal genitalia have long been used as models to address important questions in both evolutionary and developmental biology (Eberhard, 1985; Hosken & Stockley, 2004; Simmons, 2014; Sloan & Simmons, 2019). In *Drosophila*, the genitalia of both females and males are structurally diverse and rapidly evolving (Green et al., 2019; Kamimura, 2016; Muto et al., 2018; Onuma et al., 2022; Rice et al., 2023; Urum et al., 2024; Yassin & Orgogozo, 2013). In several cases, the rapid evolution of *Drosophila* genital structures has been found to involve developmental network co-option (Glassford et al., 2015; G. Rice et al., 2024). The preponderance of historical research in Drosophilids and other species has focused on the male genitalia (Ah-King et al., 2014; Méndez & Córdoba-Aguilar, 2004; Orbach, 2022). However, because female and male genitalia develop from serially homologous segments (Keisman et al., 2001), closing this sex-bias through further investigation of both sexes opens exciting opportunities to study multiple interconnected levels of homology relationships (Ah-King et al., 2014; Brennan, 2016; Brennan & Prum, 2015; Yassin & Orgogozo, 2013).

In *Drosophila melanogaster,* previous studies have carefully examined adult genital morphology of males (Rice et al., 2019; Urum et al., 2024) and females (McQueen et al., 2022). Still, challenges arise for comparative approaches involving female genital traits in *Drosophila*, as boundaries between some key genital features are less distinct compared to the male, and not all of the features that vary from species to species are rigid or external (McQueen et al., 2022; Yassin & Orgogozo, 2013). Nascent adult structures for females are especially difficult to distinguish at earlier developmental stages (Green et al., 2019). In addition, early studies of *Drosophila* genital development were performed on imaginal discs in third instar larvae, but determining how these tissues are transformed into the adult genital structures in both sexes remains challenging (Chatterjee et al., 2011). Nevertheless, researchers studying female reproductive traits in this system have overcome some of these obstacles with new methods for visualizing internal structures (Mattei et al., 2015). These achievements underscore the determination and innovative mindset of those dedicated to addressing the sex bias present in this literature.

In *Drosophila*, modern molecular methods can overcome the obstacles inherent in investigating the evolution of structures with underlying homology relationships. The application of *in situ* hybridization, immunohistochemistry, and other genetic techniques can aid in structural demarcation, even in early stages of genital development when boundaries are less distinct (Green et al., 2019; Smith et al., 2020). In previous work, we characterized the transcription factor landscape that patterns developing male genitalia in *Drosophila melanogaster (Vincent et al., 2019)*. We identified several transcription factors that serve as markers for genital structures with clear boundaries, and also found evidence that some structures may contain multiple cell types with unknown functions. This work demonstrated that the developing male terminalia is patterned by a complex network of transcription factors, and some of these genes are restricted to a few structures or cell types. However, colorimetric *in situ* hybridization does not allow for precise visualization of expression patterns for internal structures, nor the capture of relative spatial arrangement of multiple transcription factors within the same sample. Addressing these limitations will advance our ability to examine homology and structural evolution in this system using molecular methods.

In this study, we used hybridization chain reaction (HCR) and confocal microscopy to visualize the gene expression patterns of transcription factors in the genitalia of both sexes, with a particular focus on the female (Choi et al., 2018; Dirks & Pierce, 2004). Compared to colorimetric *in situ* hybridization, HCR enables greater sensitivity and resolution in three dimensions. In addition, HCR can identify putative regulatory relationships between genes because it allows for the simultaneous detection of multiple gene expression patterns in the same sample (Duckhorn et al., 2022; Schwarzkopf et al., 2021). Using this technique, we found that many transcription factors are expressed in both female and male genital tissues, and that multiple structures in the female can be subdivided into domains with distinct genetic profiles. Many of the expression patterns we observe here corroborate tissue domain delineations that were previously proposed based on a taxonomically-informed analysis of the adult morphology (McQueen et. al., 2022). Our experiments illuminate the cryptic complexity of the developing female genitalia and identify molecular markers that may inform future studies on the function and evolution of genital structures in both sexes.

## Results

### Developmental time course of female genitalia

The female genitalia of *D. melanogaster* develop primarily from the eighth abdominal segment (Epper & Nöthiger, 1982). The ninth abdominal segment is reduced in females and is the progenitor of only a few internal genital structures (Keisman et al., 2001; Sánchez et al., 2001). Notably, the opposite is true in the case of the male genitalia–the eighth segment is reduced in males, and most genital structures form from the ninth segment (Epper & Nöthiger, 1982; Schüpbach et al., 1978). The analia forms from the tenth abdominal segment in both sexes (Nöthiger et al., 1977). Here, we examined the morphology of female genitalia during pupal development, which includes timepoints during which species-specific differences have been previously traced (Green et al. 2019). At the beginning of this period, 44h after puparium formation (APF), we already observe the three main tissue groups described in McQueen et al. 2022: the analia, epigynium/hypogynium, and oviprovector/vagina (Figure 1A).

**Figure 1:**
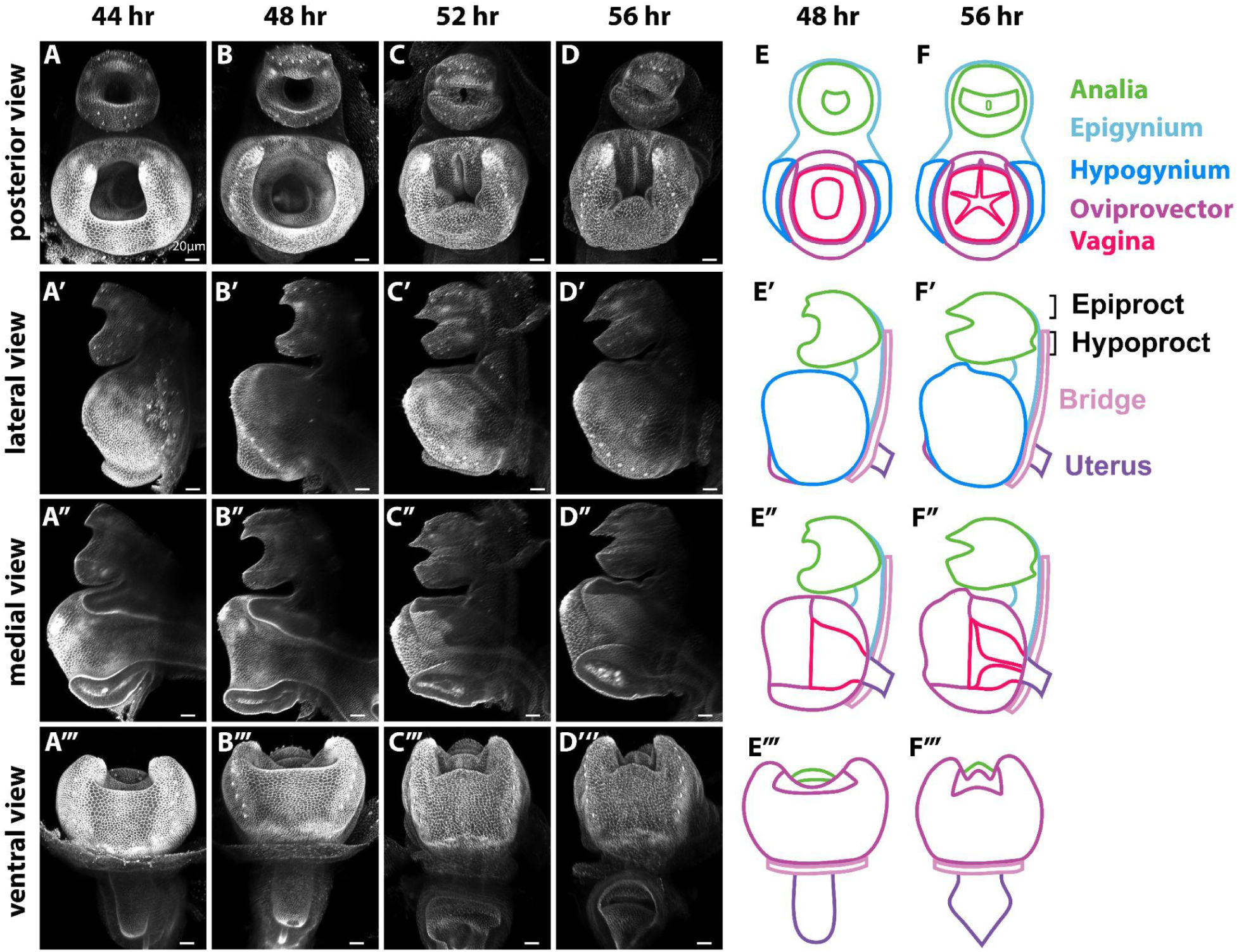
Timecourse of female terminalia development. *armadillo-GFP* fluorescence marks the apical cellular junctions and outlines the overall shape of developing female terminalia. **A, A’, A’’, A’’’:** posterior, lateral, medial, and ventral views of the terminalia at 44 hr APF. At this timepoint, the analia is tubular with a depression between the dorsal analia (epiproct) and ventral analia (hypoproct). The genitalia is also tubular with a depression on the dorsal side where the dorsal oviprovector membrane is found. **B, B’, B’’, B’’’:** At 48 h APF, the epiproct becomes more pointed at its posterior edge. The ventral oviprovector membrane begins to separate from the hypogynium and a small pocket starts to form in the dorsolateral area of the vagina. **C, C’, C’’, C’’’:** At 52h APF, the epiproct becomes more pointed and its distance relative to the hypoproct decreases. The oviprovector membrane begins to form folds that create distinct pockets along its surface. The oviprovector ventral membrane also condenses medially and pulls the hypogynium closer together. **D, D’, D’’, D’’’:** At 56h APF, the epiproct and hypoproct close further and the folds of the oviprovector membrane increase in depth. **E, E’, E’’, E’’’, F, F’, F’’, F’’’:** Schematic representations of 48h APF and 56h APF samples illustrate the analia (green), epigynium (light blue), hypogynium (dark blue), oviprovector (purple) and vagina (red). Black brackets designate the position of the epiproct and hypoproct. Scale bars represent 20μm for all images.

The two halves of the adult female analia exhibit distinct morphologies. At early timepoints, the analia is donut-shaped and divided horizontally into the epiproct on the dorsal side and the hypoproct on the ventral side (Figure 1A’,E’). As development progresses, the epiproct extends and assumes a beak-like shape, while the hypoproct becomes blunted (Figure 1A’-D’). The analia is joined to the genitalia by the epigynium, a tissue that encircles the epiproct and hypoproct (Figure 1E-F’). At 52h APF, the epigynium starts to protrude laterally (Figure 1C’, D’) and will eventually cover the anterior end of the hypogynium.

The most prominent tissue of the adult female genitalia is the hypogynium, which consists of two flat plates that form the lateral sides of the genitalia. During pupal development, the initially rounded hypogyium straightens while increasing in length and area (Green, et al. 2019). The plates of the hypogynium are connected dorsally, ventrally, and inferiorly by the membranous oviprovector (Figure 1E-F’). At 44h APF, the oviprovector is a smooth tissue, and over time, both the dorsal and ventral hypogynial membranes condense and pull the hypogynial plates closer together by 56h APF (Figure S1A-D). The vagina, which represents the anterior internal portion of the genitalia, appears cylindrical at 44h APF (Figure 1A). Over developmental time, five vaginal folds form established: one dorsal (the vaginal furcal dorsal fold), a dorsolateral pair (the vaginal furcal dorsolateral folds), and a lateral pair (the vaginal furcal lateral folds) (Figure 1C-D, Figure S1C’-D’). These folds change the vaginal morphology from a hollow structure to one that has little vacant medial space.

**Figure S1:**
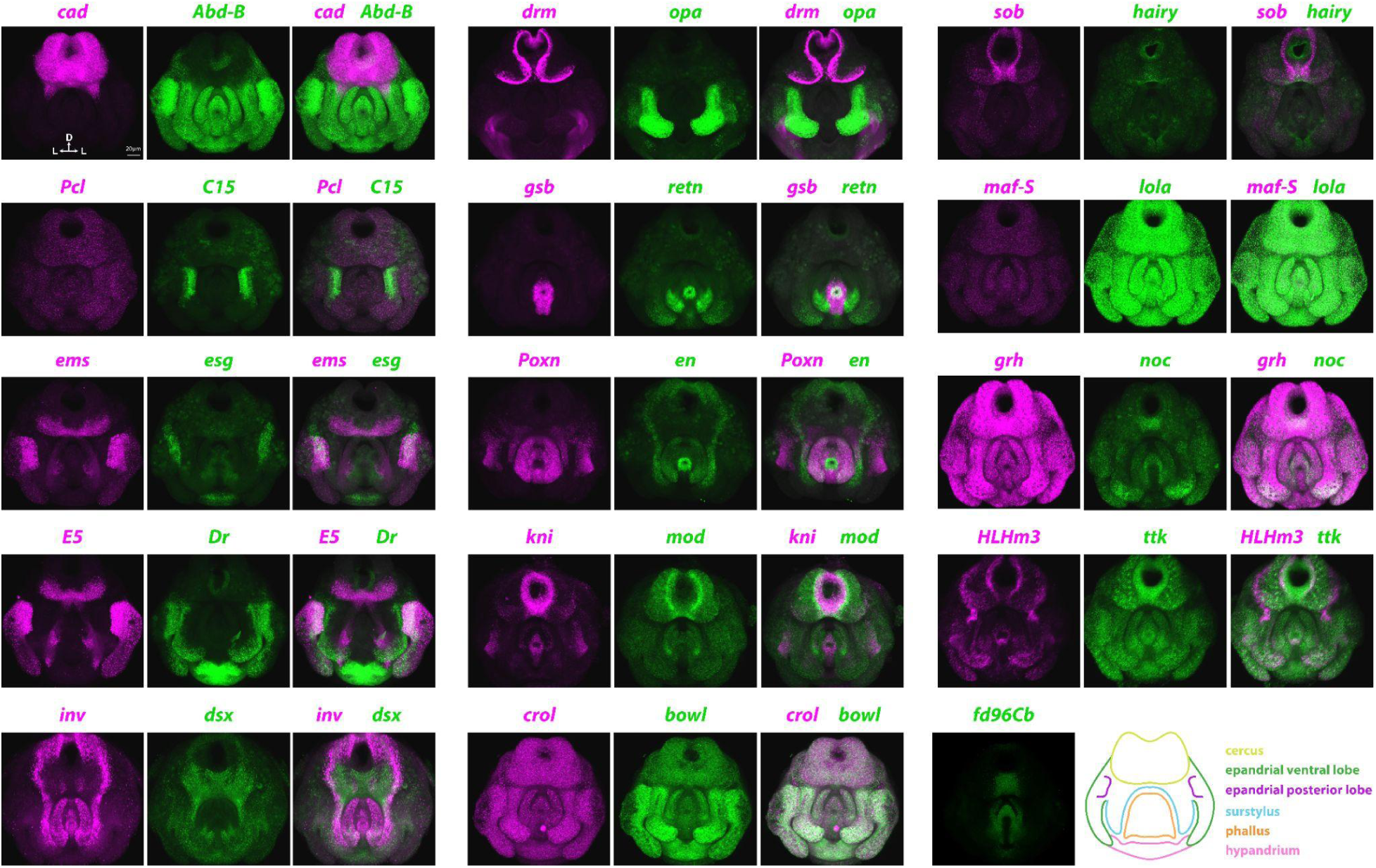
Hybridization chain reaction signal in male pupal terminalia Gene expression patterns for 29 genes were measured in male terminalia at 48h APF using HCR. HCR Probes were multiplexed, which allowed for the acquisition of two gene expression patterns in each sample. Patterns measured using the B1 amplifier are shown in magenta, while patterns measured using the B3 amplifier are shown in green. A schematic representation of the 48h APF male terminalia is shown in the bottom right.

### Validating hybridization chain reaction in male genital tissues

We previously measured the expression patterns of 100 transcription factors in the male pupal genitalia at 48h APF using colorimetric *in situ* hybridization (Vincent et al. 2019). Here, we used hybridization chain reaction (HCR) to measure expression patterns in the female pupal genitalia for 29 transcription factors that exhibited strong spatial patterning in male samples (Dirks & Pierce, 2004; Duckhorn et al., 2022). Among other advantages, HCR enables the collection of 3D images using confocal microscopy. We first validated the efficacy of the protocol by comparing our HCR signal in male genitalia to previous signal by colorimetric *in situ* hybridization (Vincent et al., 2019). At 48h APF, signals for 28 out of 29 genes showed similar expression patterns for both methods (Table S1, Figure S1). The only notable difference was observed for *Polycomblike* (*Pcl*), which had ubiquitous expression in the case of HCR, while our previous results indicated that *Pcl* expression was restricted to the phallus.

Overall, our HCR signal exhibited reduced background and an increased range of detection compared to previous results, and revealed features of genital expression patterns that were not previously noted (Choi et al., 2018). For example, while previous analyses of the *Drop* (*Dr*) expression pattern were restricted to the analia, medial clasper (i.e. surstylis), and the hypandrium (Vincent et al. 2019), HCR revealed additional *Dr* expression in the lateral plate (epandrial ventral lobe), ventral clasper, and posterior lobe (epandrial posterior lobe) (Figure S1). The concordance between our HCR results and the previous *in situ* hybridization analysis suggested that we could successfully deploy HCR to measure 3D gene expression patterns in female pupal genitalia.

**Figure S2:**
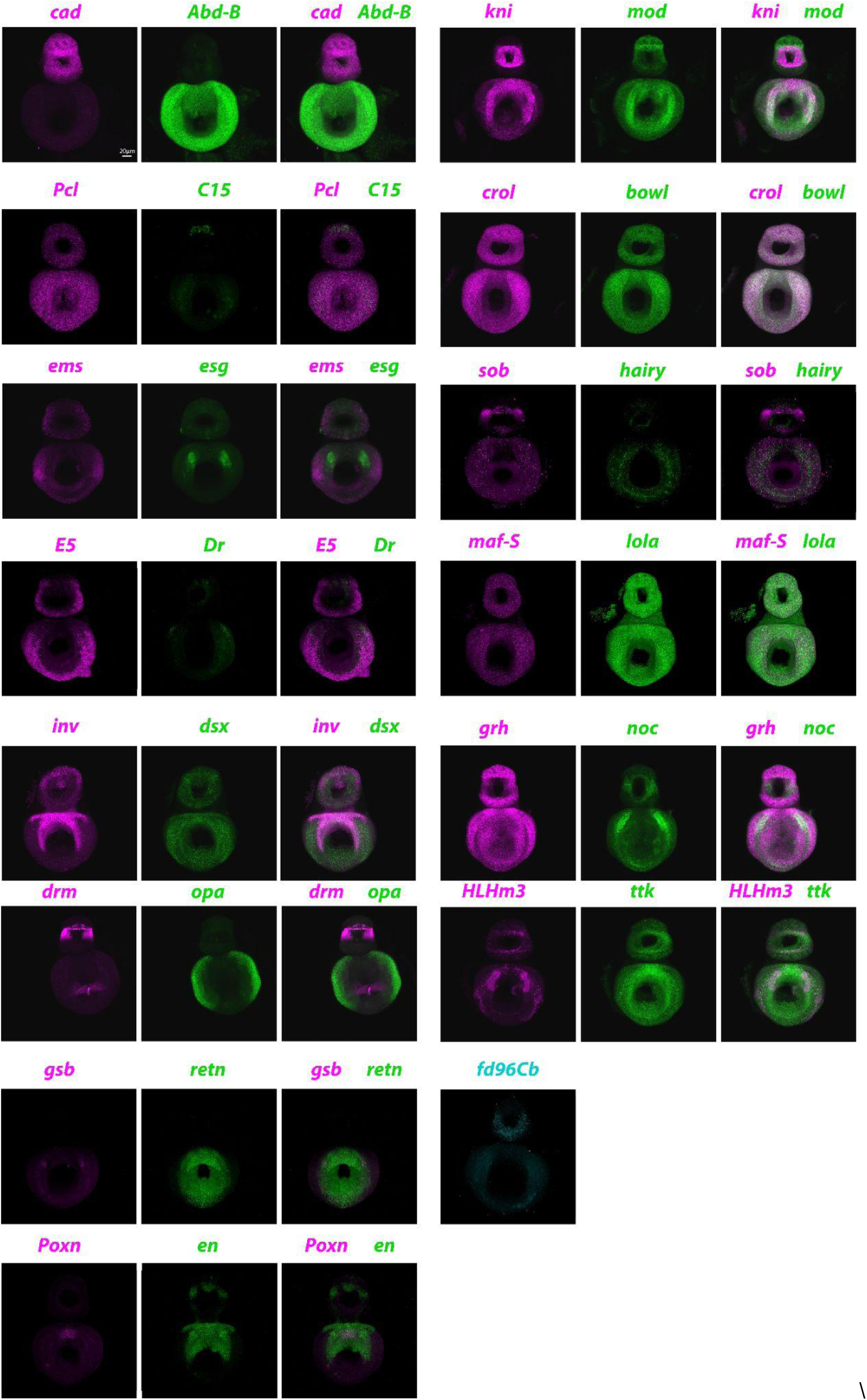
Hybridization chain reaction signal in female pupal terminalia Gene expression patterns for 29 genes were measured in male terminalia at 48h APF using HCR. HCR Probes were multiplexed, which allowed for analysis of two gene expression patterns in each sample. Patterns measured using the B1 amplifier are shown in green, while patterns measured using the B3 amplifier are shown in magenta. *fd96Cb* which used the B4 amplifier is shown in grey.

### Transcription factors with ubiquitous expression patterns

Eight transcription factors displayed ubiquitous or nearly ubiquitous expression patterns in the genitalia of both males and females. In one case, the gene *hairy*, gene expression in the males was ubiquitous, but was notably absent from the epigynium in female samples (Figure S2). In most of these cases, although present in all tissues, gene expression was stronger in some tissues than others. Observation of tissue-level variation in the ubiquitous expression is indicated in **Table 1** and **Table S1**. One illustrative example is the gene *bowl*, which shows low levels of expression across the entire genitalia in both males and females, but is strongly expressed in the dorsal parts of the female genitalia and the external structures of the male genitalia (Figure S1, S2).

**Table 1:**
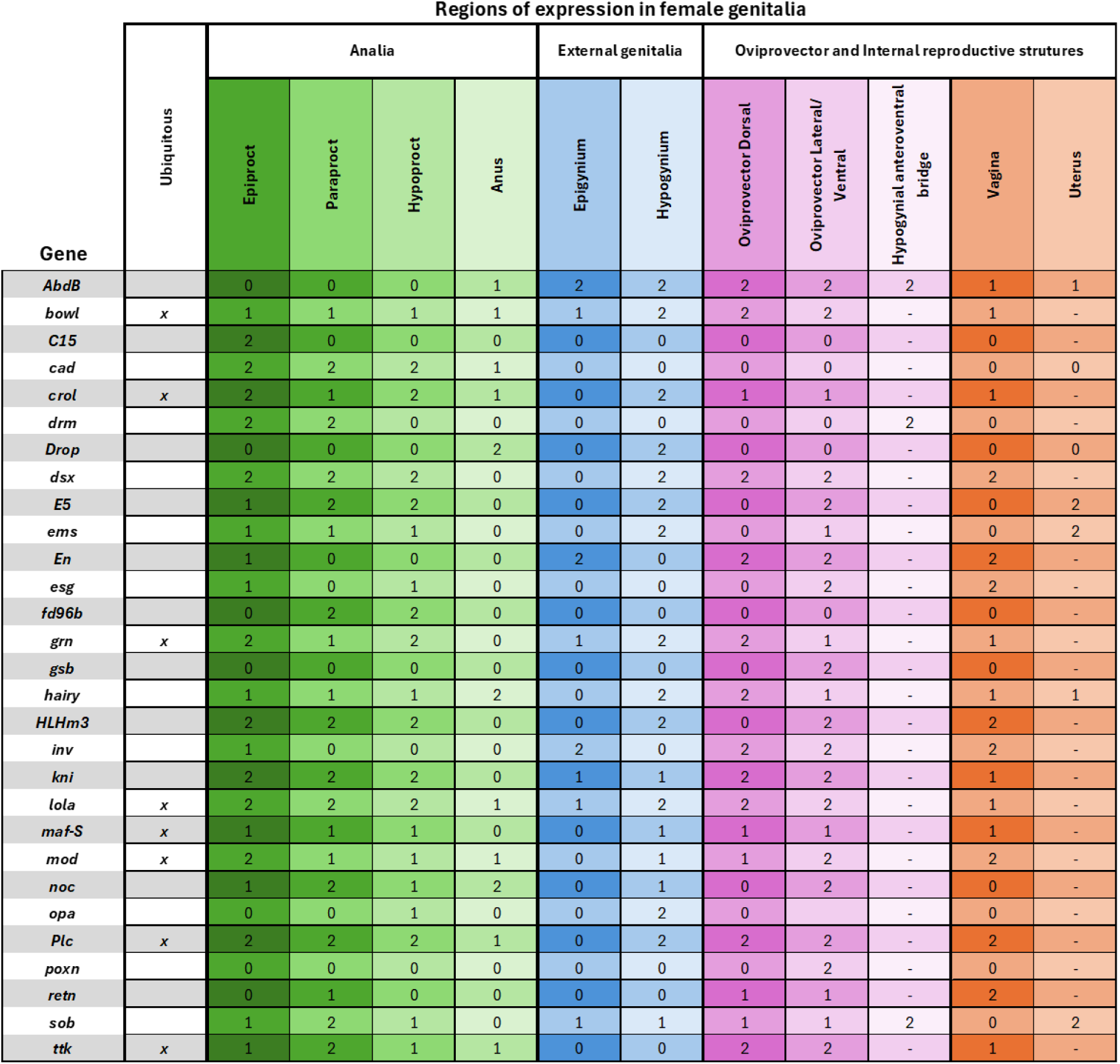
Summary of transcription factor expression in female genital structures. Image stacks of HCR signal were manually inspected, and expression levels were qualitatively assigned to one of four categories for each structure in the developing female genitalia: strongly expressed (2); weakly expressed (1); not expressed (0); and could not be determined from available data (-). Genes exhibiting ubiquitous expression are also noted, with additional information regarding tissue-level variation in expression indicated in the table as needed.

### Expression patterns in the female and male analia

The analia comprise the ventral-most tissues of the terminalia in both sexes. In the analia, 18 out of the 29 transcription factors we screened showed patterned expression in females (Figure 2, Figure S3). The gene *cad* (Figure S1) was previously identified as a marker of the anal plate in males (Moreno & Morata, 1999; Vincent et al., 2019). Likewise, *cad* exhibits strong expression throughout the female analia (Figure 2C). Although the analia of both sexes is derived from the tenth abdominal segment, in females the anal plate is divided horizontally, which produces two distinct substructures. In males, the anal plate is divided vertically and exhibits bilateral symmetry (Green et al., 2019) (Figure 2B). Despite this difference, some of the genes we examined are patterned similarly in both sexes. For example, *Dr* is expressed in the anus in both males and females, (Table 1, S1, Figure 2C, S1), as is *hairy, Abd-B, cad,* and *noc* (Figure S1, S3). *HLHm3* exhibits medial expression in both the female epiproct and hypoproct (Figure 2C), which mimics its similar expression pattern in the in male anal plate, except that the pattern is rotated 90° (Figure S1). Moreover, in both males and females, the genes *drm*, *sob,* and *kni* are expressed in a third, distinct tissue located between the two lobes of the anal plate in males, and between the epiproct and hypoproct in females. We assign the term “paraproct” to this structure (Figure 2C). Some genes, including *E5*, *ems*, and *inv*, are expressed distinctly in either the epiproct or hypoproct in females and also exhibit dorsal/ventral patterning in males (Figure S1, S3, S6). Conversely, some genes are markers of specific subsections of the female analia and have distinct patterns when compared to the males. For instance, *C15* is restricted to the epiproct (Ridgway, Hood, et al., 2024), while the *fd96Cb* marks the hypoproct (Figure 2C). We did not detect C15 in the analia of males, whereas *fd96Cb* is present in males but only in the ventral paraproct (Figure S1).

**Figure 2:**
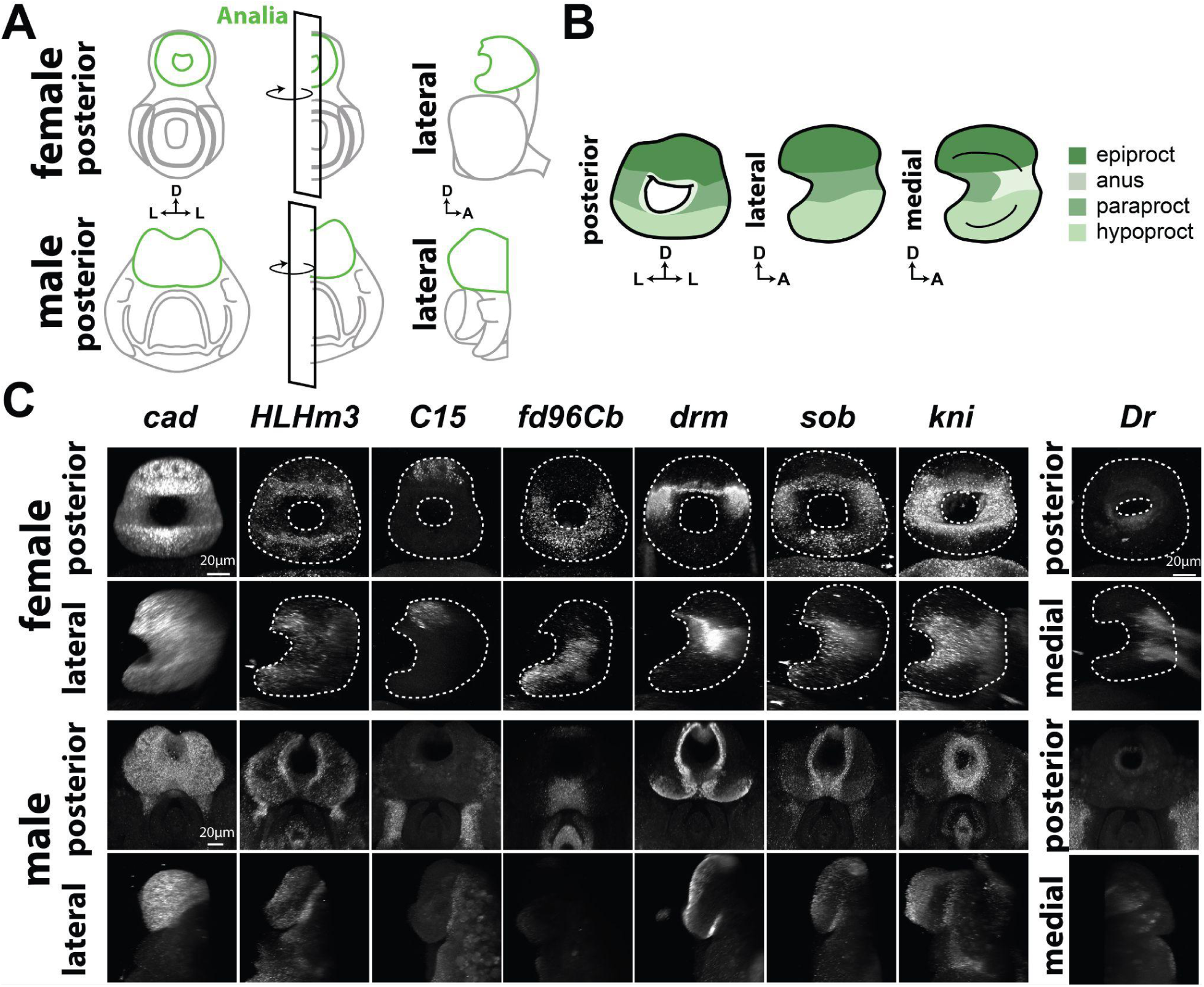
Transcription factors expressed in the analia. **A)** Schematics of the *Drosophila melanogaster* female and male terminalia with the analia highlighted in green. For lateral views the left half of the terminalia is masked and the sample is turned to face the lateral side. **B)** Schematic of the tissues within the female terminalia: epiproct (dark green), anus (grey-green), paraproct (sage green), and hypoproct (light green)**. C)** HCR signal in female and male analia, with detected signal rendered in white. *cad* is expressed throughout the epiproct and hypoproct. *HLHm3* is expressed in both the epiproct and hypoproct. *C15* is restricted to the epiproct, while *fd96Cb* is restricted to the paraproct and hypoproct. *drm*, *sob*, and *kni* are mainly expressed in the paraproct. *Dr* is expressed in the anus, which connects the analia to the rectum.

**Figure S3:**
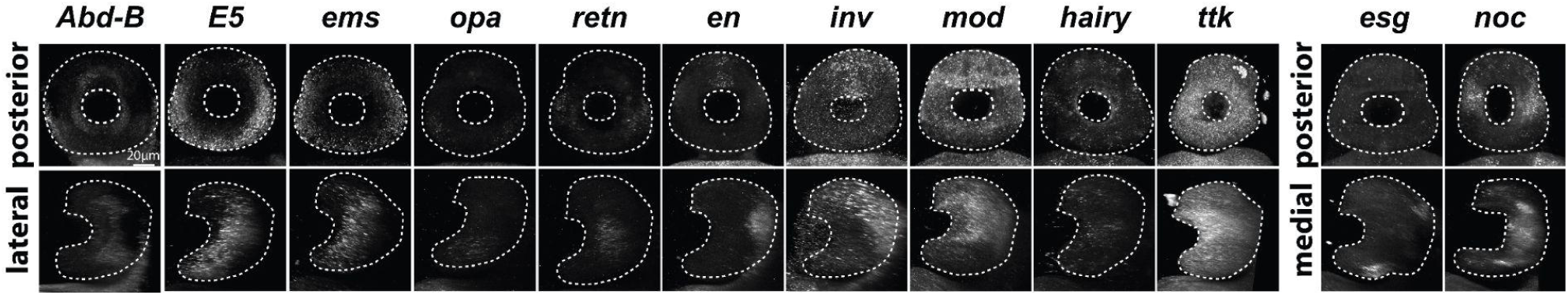
Additional transcription factors expressed in the female analia HCR signal for transcription factors in female analia, with detected signal rendered in white. *Abd-B* is expressed in the anus, while *E5* and *ems* show strong expression in the edge of the hypoproct and the lateral region of the epiproct, respectively. *opa* is expressed in the lateral region of the hypoproct, while *retn* is expressed in the paraproct. *noc* is expressed in the paraproct and anus, and *en* and *inv* are expressed in the medial region of the epiproct. *mod* is expressed throughout the analia, but weaker expression can be detected in the hypoproct. *esg* is expressed in the ventromedial hypoproct and posterodorsal epiproct. Medial views for *esg,* and *noc* are shown on the bottom right.

### Expression patterns in the female external genitalia: hypogynium and epigynium

The female analia is connected to the genitalia via a cuticular structure called the epigynium (Figure 3A). Aside from ubiquitously expressed genes, we found that few of the screened transcription factors exhibited patterned expression in the epigynium (**Table 1**). The transcription factors *en* and *inv* (Figure 3B) are expressed strongly throughout the epigynium; these patterns wrap around the anterior side of the analia to the hypogynial plates (Figure S4). Notably, this pattern in which *en* and *inv* expression wraps around the anal plate is also observed in males (Figure S1). The gene *kni* is also weakly expressed in parts of the epigynium (Figure S2).

**Figure 3:**
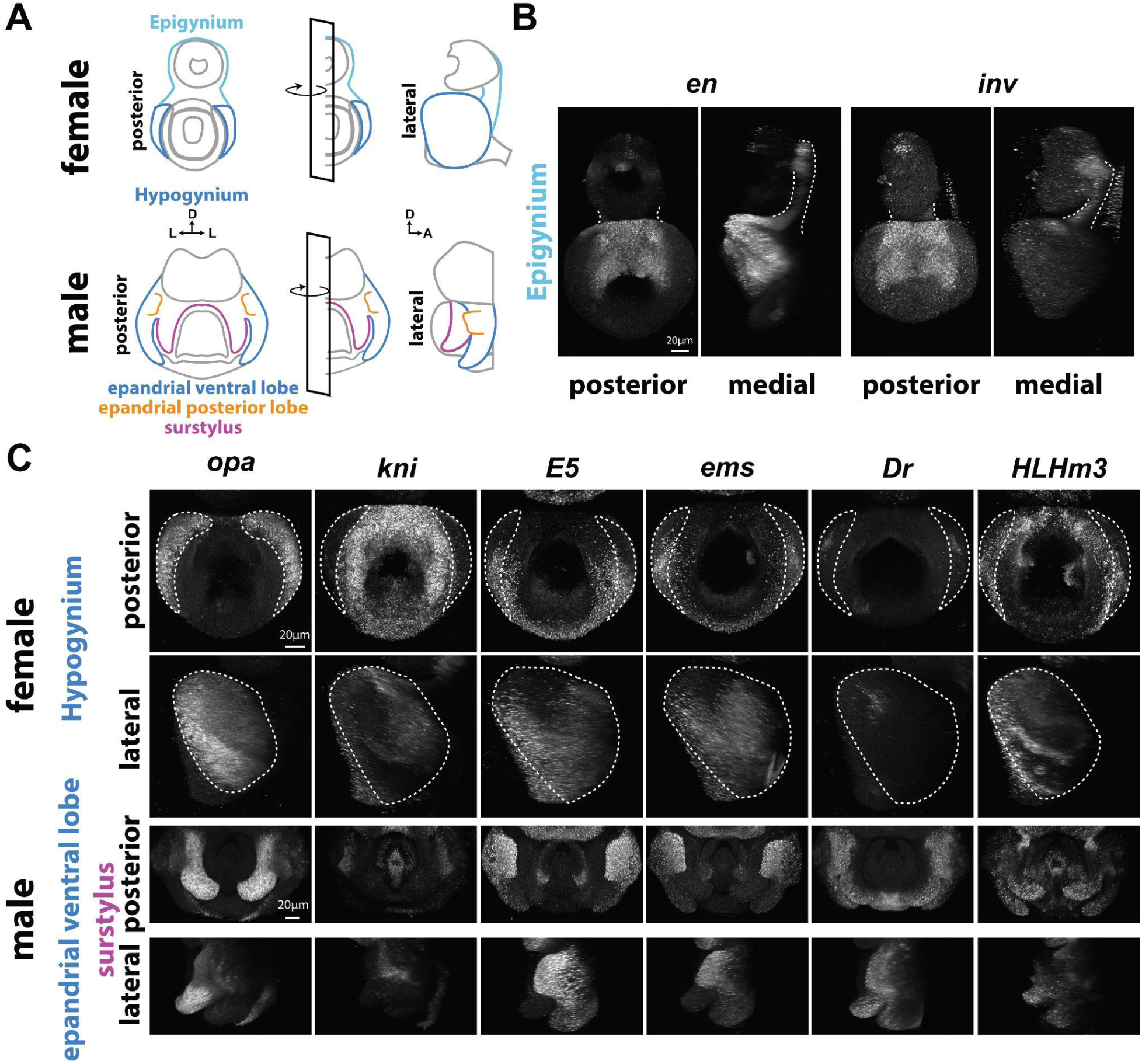
Transcription factors expressed in the epigynium and hypogynium. **A)** Schematics of the female and male terminalia with the female epigynium (light blue), and hypogynium (dark blue), as well as the male epandrial ventral lobe (dark blue), epandrial posterior lobe (orange), and surstylus (purple) highlighted. **B)** HCR signal in the female epigynium, where the detected signal appears white. *en* and *inv* are expressed across the hypogynium. **C)** HCR signal in the female hypogynium and male epandrial ventral lobe and surstylus. Posterior (top) and lateral views (bottom) are shown. *opa* is expressed in the posterior edge of the hypogynium, while *kni* expression is restricted to the anterior region of the hypogynium. *ems* and *E5* are expressed broadly in the epandrial ventral lobe but decrease in expression midway through the anterior-posterior axis. *Dr* is expressed in the posteroventral hypogyium. *HLHm3* is expressed throughout the dorsal hypogynium and in a punctate pattern in the ventral hypogynium.

**Figure S4:**
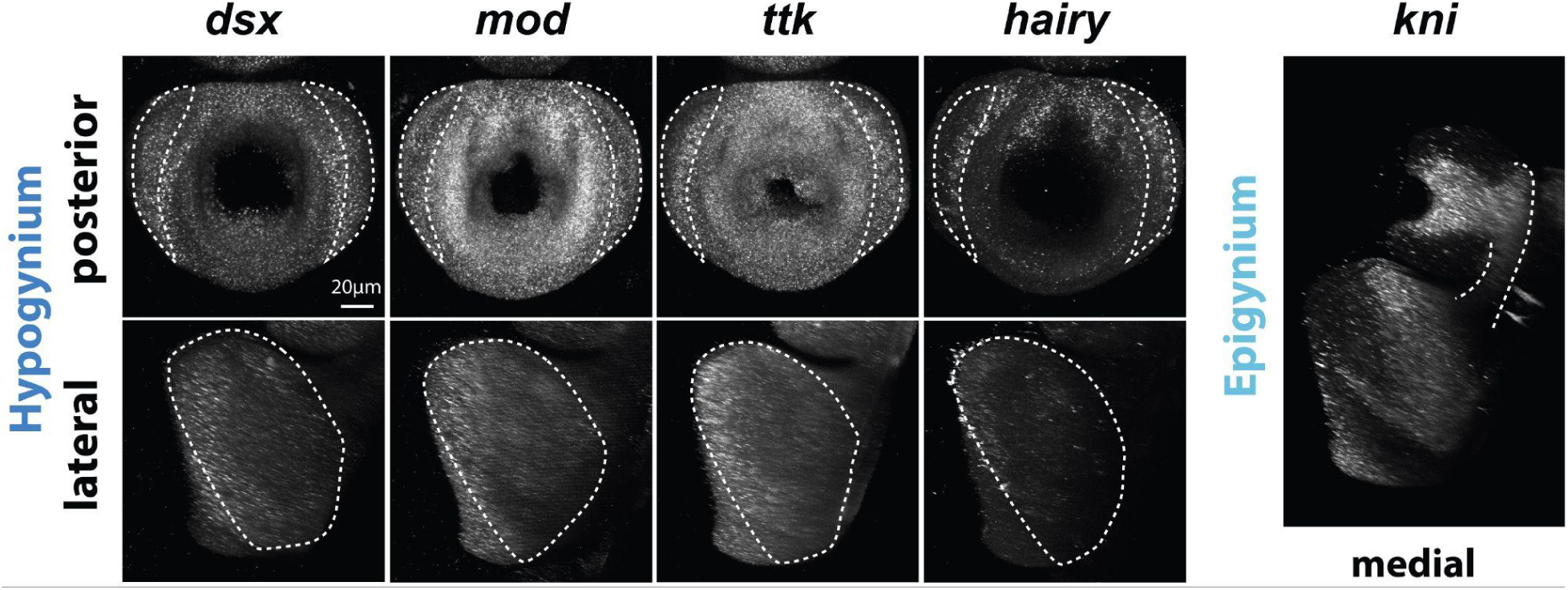
Additional transcription factors expressed in the epigynium and hypogynium. HCR signal in the female hypogynium and epigynium, where the detected signal appears white. *dsx, mod,* and *ttk* are expressed throughout the hypogynium but show reduced expression in the anterior region. *hairy* expression is restricted to the anterior region of the hypogynium. *kni* is expressed throughout the epigynium.

The hypogynium plays a key functional role during oviposition in many pest species, and is one of the best-studied female genital structures in *Drosophila (Atallah et al., 2014)*. For instance, previous work traced *Drosophila suzukii*’s increased hypogynial length–a derived state that facilitates more penetrant oviposition in that species–to altered cellular morphogenesis (Green et al., 2019). Understanding the developmental landscape of this structure is therefore of substantial interest. We found that many transcription factors in our panel exhibit patterned expression in the hypogynium. For example, o*pa* exhibits strong expression along the entire posterior edge of each hypogynial plate, while *kni* is restricted to the anterior region (Figure 3C). The genes *E5* and *ems* are also expressed throughout the anterior and ventral parts of the hypogynium, with some weak expression along the ventral posterior edge (Figure 3C). *Drop* expression is restricted to a small dorsolateral patch on each plate (Figure 3C). Interestingly, co-staining of *Dr* and *E5* reveals that the *Dr* expression pattern is anticorrelated with *E5*, except for a very small section of overlap between the two patterns (see Figure 5C). A similar phenomenon occurs between *E5* and *Dr* in males, where the two genes are expressed in separate regions except for a small region of overlap in the epandrial ventral lobe (see Figure 5C). The expression pattern for *HLHm3* in the hypogynium is restricted to a stripe of expression along the posterior edge of each plate (Figure 3C). This pattern begins at the dorsal posterior tip of the plates and then bifurcates part way along the posterior edge, with one stripe of expression continuing along the posterior edge and the other continuing in a ventral trajectory slightly further towards the anterior portion of the tissue. In adults, the cuticle in this area of each hypogynial plate is partly, but not entirely, bisected–a feature called the hypogynial mid-dorsal incision (McQueen et al, 2022). The expression pattern of *HLHm3* may be associated with the patterning of the mid-dorsal incision, which could be investigated in samples at later time points.

### Expression patterns in the oviprovector and vagina

The oviprovector is a membranous tissue interior to the hypogynium that connects the hypogynium to the vagina (Figure 4, A and D). Although contiguous at early stages of development, the adult oviprovector comprises two parts–a dorsal membrane and a ventral membrane. The ventral membrane also extends dorsally and harbors a set of scales (McQueen et al, 2022). In *Drosophila erecta* and *D. teissieri,* the dorsal oviprovector has evolved pocket and shield-like structures that may protect against copulatory harm (Kamimura, 2016; Kamimura & Mitsumoto, 2012).

**Figure 4:**
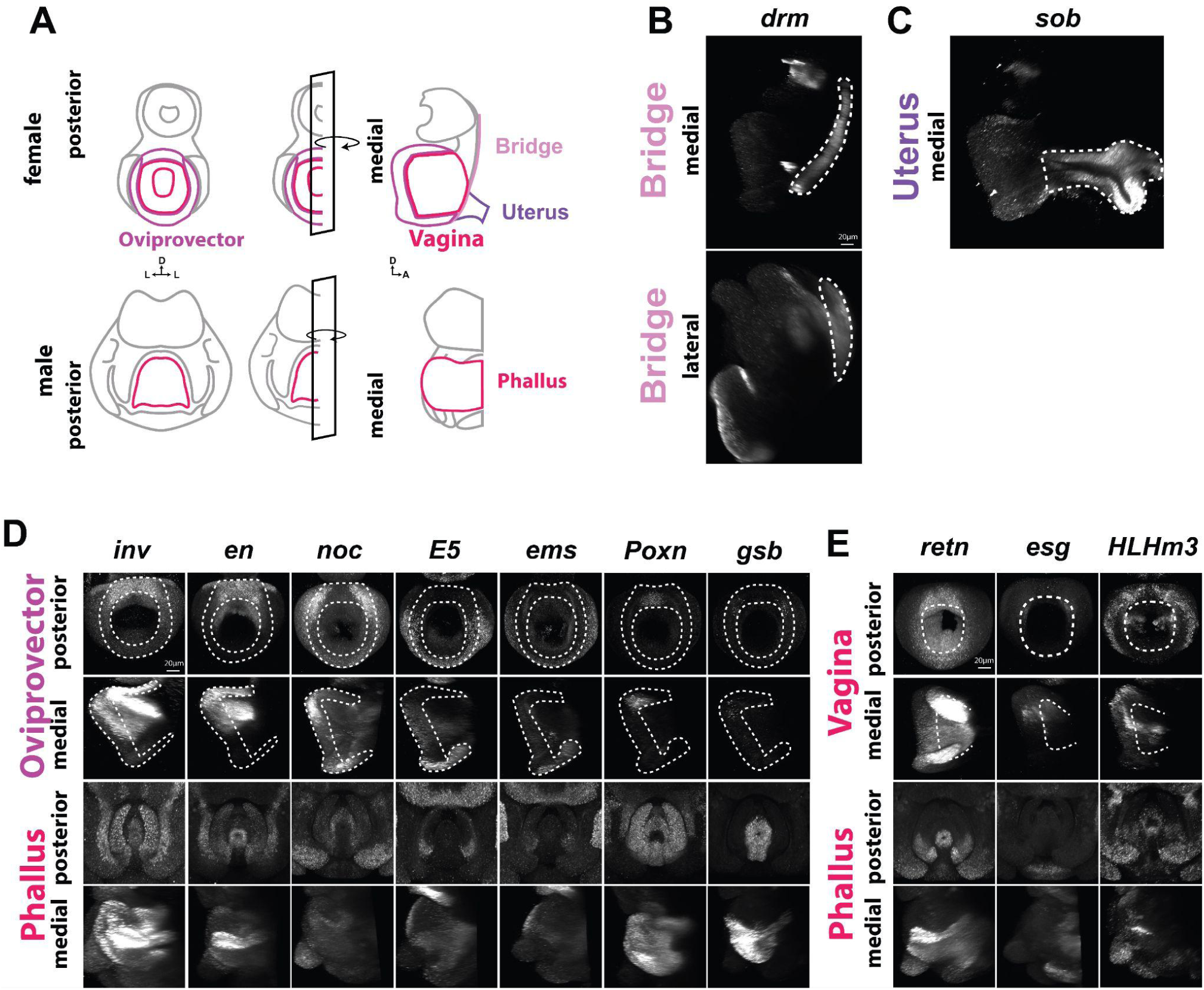
Transcription factors expressed in the oviprovector and internal genitalia. **A)** Schematics of the female and male terminalia with the female oviprovector (light purple), vagina (red), bridge (pink), and uterus (dark purple) highlighted in females and the male phallus highlighted in red. **B)** HCR signal in the bridge (dashed line) for both sexes. **C)** HCR signal in the female uterus (dashed line). **C)** HCR signal in the female oviprovector (dashed line) and male phallus, where the detected signal appears in white. *inv, en*, and *noc* are expressed in the dorsal oviprovector, while *inv* and *en* are also expressed in the dorsal vagina. *E5*, and *ems* are expressed in the lateral and ventral regions of the oviprovector. *Poxn* is expressed in the dorsomedial oviprovector and *gsb* is expressed in the dorsolateral oviprovector. **D)** HCR signal in the female vagina (dashed line) and male phallus. *retn* is expressed throughout the vagina. *esg* and *HLHm3* are expressed in the dorsal lateral portion of the vagina.

**Figure S5:**
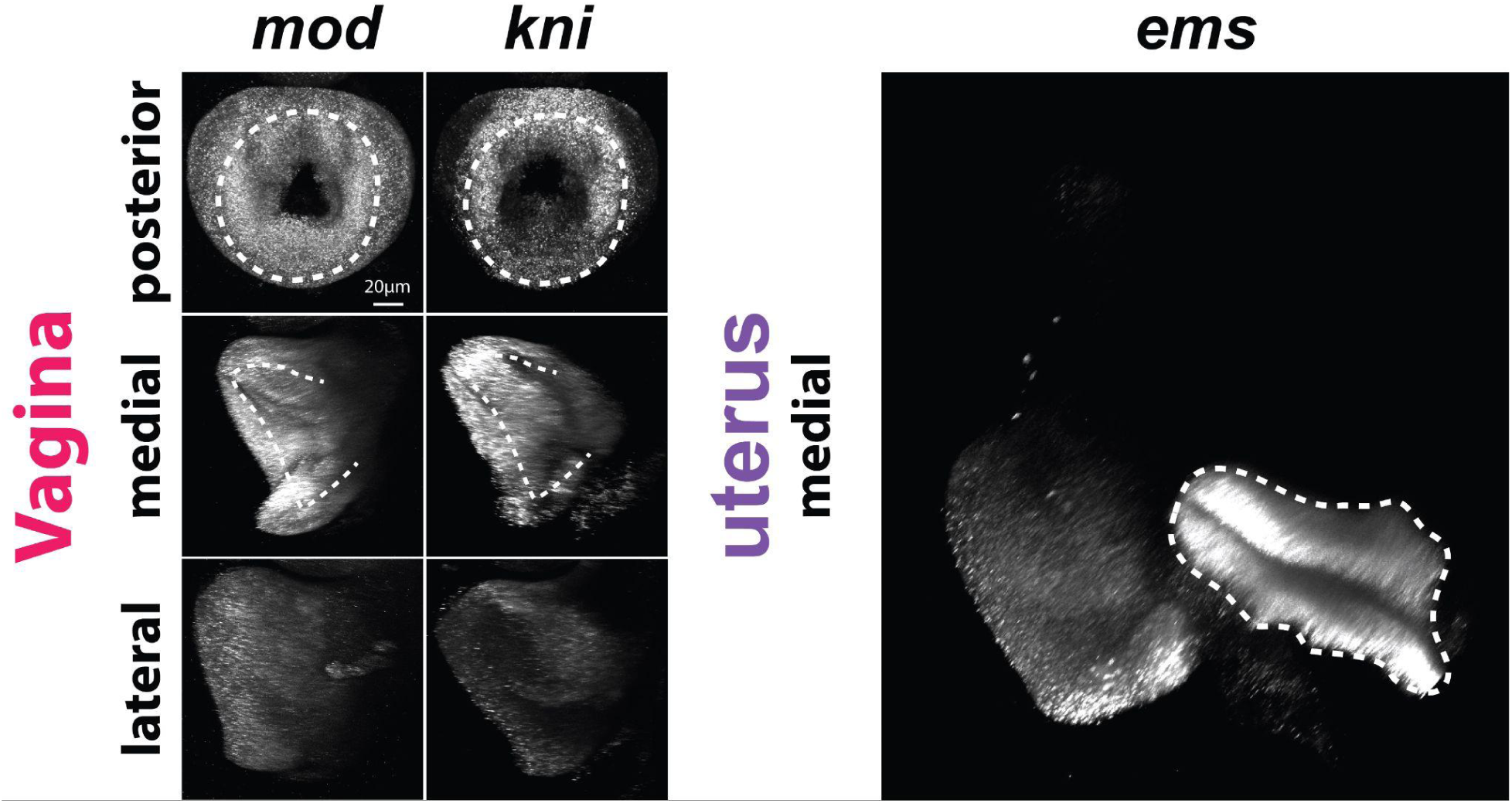
Additional transcription factors expressed in the oviprovector and internal genitalia HCR signal in the female vagina and uterus, where detected signal appears in white. *mod* and *kni* are expressed throughout the vagina and in the oviprovector. *ems* is expressed throughout the uterus.

Many of the transcription factors we surveyed are expressed in the developing oviprovector. For example, *en* and *inv* are expressed throughout the dorsal oviprovector and partially extend ventrally (Figure 4D), whereas the *Poxn* expression pattern is restricted to the medial portion of the dorsal oviprovector (Figure 4D). *gsb* exhibits a highly specific expression pattern at a location that may represent the boundary between the dorsal and lateral/ventral regions of the oviprovector (Figure 4D). *noc* is strongly expressed in the dorsolateral portion of the oviprovector, while *E5* and *ems* are expressed in the ventral oviprovector (Figure 4D).

The vagina is located anterior (with respect to the whole animal) to the developing oviprovector. Although it is difficult to determine the exact boundary between the developing oviprovector and vagina at 48h APF, some transcription factors are clearly expressed in the anterior region associated with vaginal tissue. A key example is *retn* (Figure 4E). In contrast, *esg* is expressed in regions of the anterior (more internal) oviprovector as well as the most posterior portions of the vagina (Figure 4E) in a pattern that suggests a possible association with some of the eventual vaginal furcal folds. In these internal tissues, *HLHm3* is expressed in two lateral stripes that extend anteriorly from the oviprovector into the vagina (Figure 4E). This expression pattern may mark a boundary between tissue types that differentiate later in development.

We found that two transcription factors, *drm* and *sob*, serve as markers for the hypogynial anteroventral bridge, a tissue that connects the terminalia to the abdominal tergite 7 (Figure 4B). Although this named structure in adults is restricted to a ventral fragment (McQueen et al, 2022), our results suggest that the hypogynial anteroventral bridge shares developmental origins with the anterior edge of the epigynium, which wraps around the anterior portion of the analia. For many transcription factors that we screened, we were unable to determine with confidence whether they were expressed in the ventral portion of the hypogynial anteroventral bridge (see Table 1).

In this study, the uterus was only retained in a subset of dissected samples, which allowed us to determine whether some transcription factors were expressed in this tissue. For example, *sob* (Figure 4C), *ems* (Figure S5), and E5 were all strongly expressed in the uterus.

**Figure 5:**
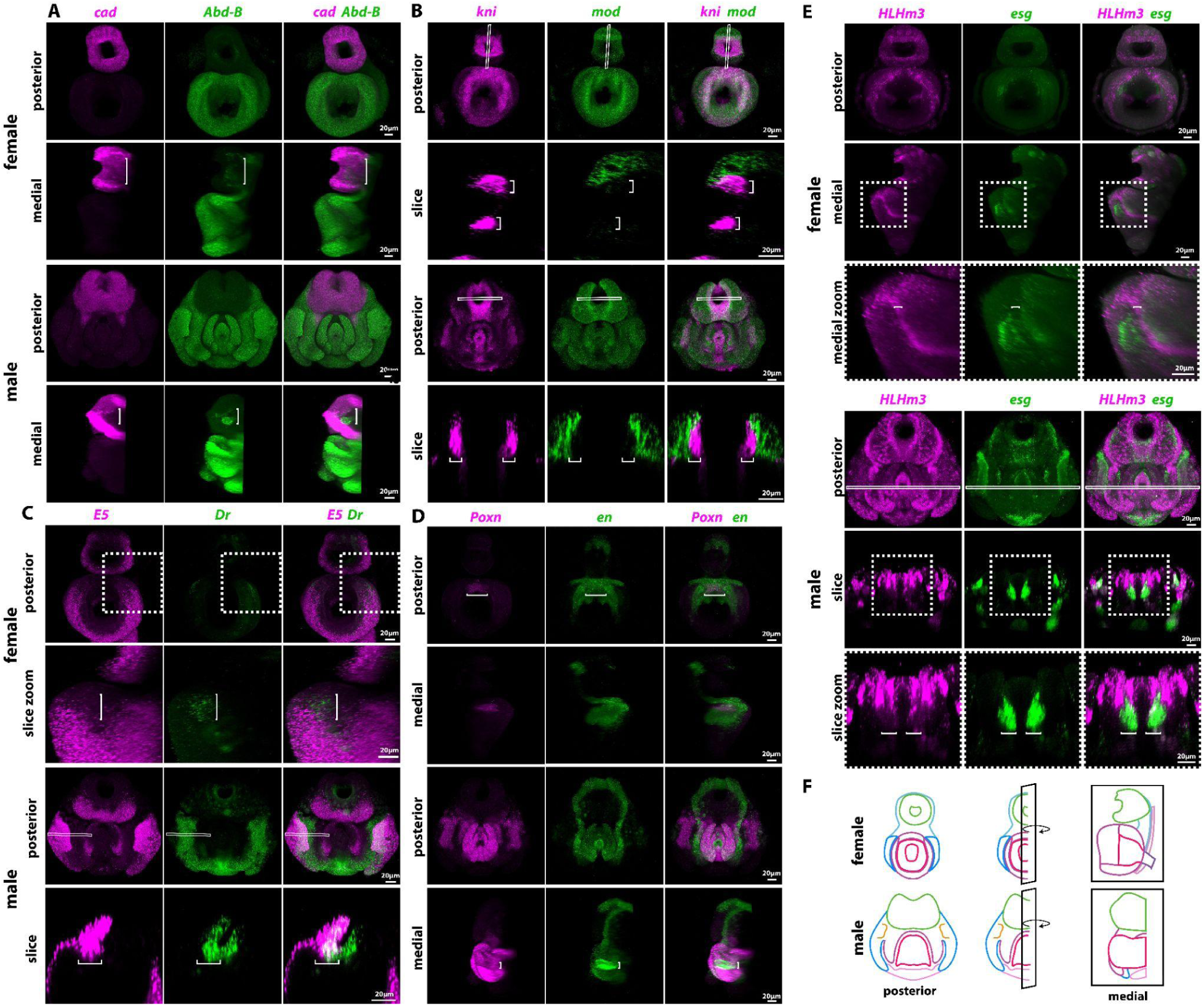
Multiplexed HCR reveals fine scale relationships between gene expression patterns. **A)** HCR siganl for *cad* (magenta) and *Abd-B* (green) in female and male pupal genitalia. In females, *cad* is expressed in the hypoproct and epiproct but not in the anus (white brackets). *Abd-B* is expressed in the genitalia as well as the anus (brackets) in both sexes. **B)** HCR signal for *kni* (magenta) and *mod* (green). *kni* is expressed in the female inner epiproct (brackets) but not in the outer portion (brackets), while *mod* is expressed in the outer epiproct and not the internal portion. The male analia shows a similar pattern to the females, but rotated 90 degrees. **C)** HCR siganl for *E5* (magenta) and *Dr* (green). *Dr* is expressed in the hypogynal posterior lobe, while *E5* is expressed ventrally relative to the *Dr* expression domain (brackets). In the male epandrial posterior lobe, both *E5* and *Dr* are co-expressed along the medial edge, while only *E5* is expressed on the lateral edge. **D)** HCR siganl for *Poxn* (magenta) and *en* (green). *PoxN* expression is restricted in the medial portion of the female oviprovector dorsal membrane (brackets), while *en* is expressed throughout the oviprovector dorsal membrane. *Poxn* is expressed throughout the male phallus but is restricted from the tube that runs through it (known as the phallotrema) while *en* shows strong expression in the phallotrema (brackets). **E)** HCR signal for *HLHm3* (magenta) and *esg* (green). *HLHm3* and *esg* are both expressed in the female oviprovector ventral membrane in close proximity. A close up of the indicated region (dashed box) reveals that these expression patterns do not overlap. *HLHm3* expression splits two *esg* expression domains (brackets). The male phallus shows anterior expression of *esg* (brackets) directly adjacent to posterior *HLHm3* expression pattern. **F)** Schematic of the posterior and medial views of the *Drosophila melanogaster* terminalia. For female terminalia, we highlight the analia (green), epigynium (light blue), hypogynium (dark blue), oviprovector (light purple), vagina (red), bridge (pink), and uterus (dark purple). For male terminalia, we highlight the analia (green), epandrial ventral lobe (dark blue), epandrial posterior lobe (orange), surstylus (purple), phallus (red), and hypandrium (pink). Solid boxes indicate the location of an optical slice through the samples.

### Co-expression of transcription factors in males and females

HCR accommodates multiplexing–i.e. detecting RNA for multiple genes in the same sample. Multiplexing allows researchers to define precise expression domains of genes relative to each other with high confidence. For instance, multiplexed *AbdB* and *cad* HCR signals reveal that these two transcription factors are directly adjacent and mutually exclusive (Figure 5A). Previous expression data suggested that *cad* expression was limited to the anal plate, and *AbdB* was absent in that tissue (Vincent et al., 2019). We find that *AbdB* also has low-level expression in the anus and the inner tissue of the analia of females and males, but this pattern remains adjacent to and distinct from *cad*-expressing cells (Figure 5A). In addition to this mutually exclusive gene pair, *kni* expression in the inner ring of the analia is adjacent to a *mod* expression domain in the epiproct (Figure 5B), while *Dr* is expressed directly dorsal to *E5* in the hypogynial posterior lobe (Figure 5C). Finally, in the vagina, *HLHm3* is expressed between two adjacent *esg* expression domains (Figure 5D). We also find examples of co-expressed transcription factors, as the *Poxn*’s medial expression is entirely within the broader *en* expression in the oviprovector dorsal membrane (Figure 5E). These genes are also co-expressed in males, specifically in tissues surrounding the phallus and the ventral portion of the clasper (Figure S1). These examples of mutual exclusion and co-expression may indicate negative and positive regulatory interactions respectively, which can be interrogated further in future experimental work.

**Figure S6:**
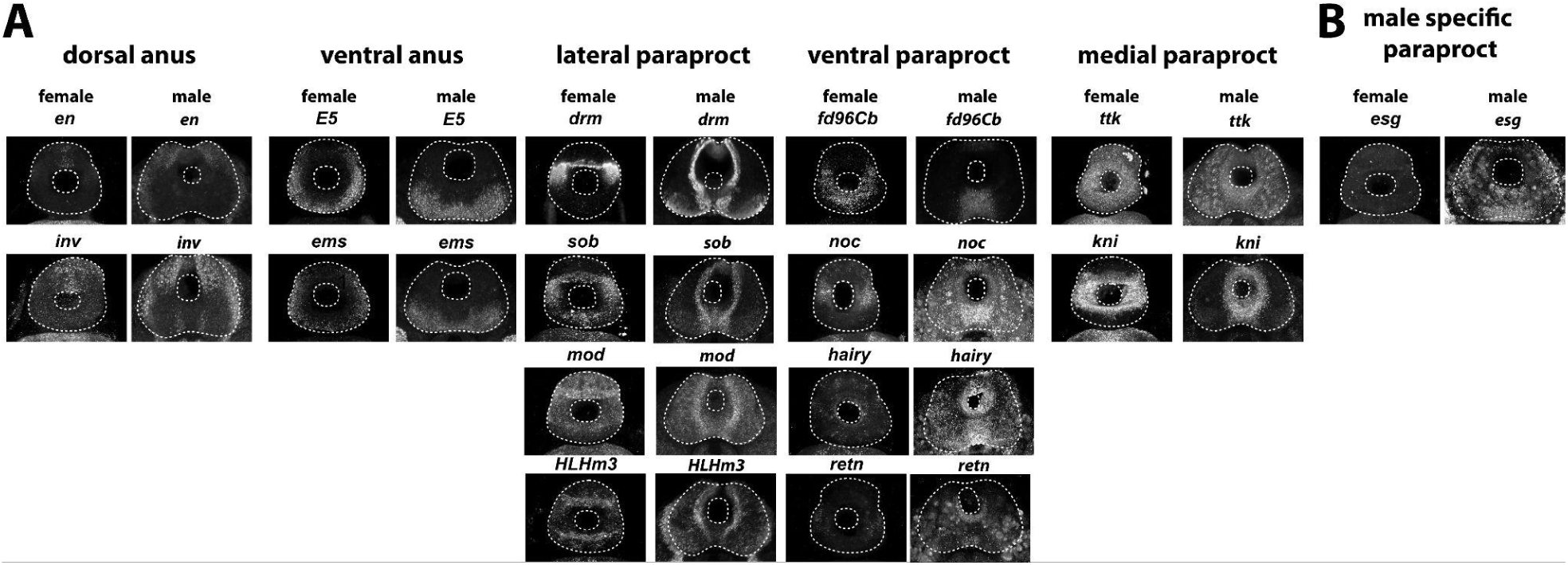
Shared male and female analia expression patterns HCR images where white represents expression. **A)** *en*, and *inv* are expressed in the dorsal anus of males and females. *E5*, and *ems* are expressed in the dorsal anus of males and females. *drm*, *sob*, *mod*, and *HLHm3* are expressed in the lateral paraproct region of males and females. *fd96Cb*, *noc*, *hairy,* and *retn* are expressed in the ventral paraproct region of males and females. *ttk*, and *kni* are expressed in the medial paraproct region of males and females. **B)** *esg* is expressed only in the paraproct of the male.

## Discussion

In this work, we present a detailed 3D atlas of transcription factors expressed in developing genital structures in *Drosophila melanogaster*. We collected 3D images of genital morphology at four time points during pupal development in females, and we measured the expression patterns of 29 transcription factors at 48h APF in both females and males. This analysis confirmed and extended previous results found using colorimetric *in situ* hybridization in males (Vincent et al., 2019) and revealed a complex gene regulatory landscape in female tissues. Our results suggest that some morphological observations in a recent detailed analysis of adult female genitalia may have molecular underpinnings (McQueen et al., 2022). For example, we found that certain transcription factors can serve as marker genes in the analia (*cad*), hypogynium (*opa*), and vagina (*retn*). We also found that expression patterns in the female analia are subdivided along the dorsal/ventral axis, compared with the medial/lateral organization of the male analia. Finally, multiplexing of hybridization chain reaction signal allowed us to label expression domains of transcription factor pairs in single tissues, which suggested putative regulatory interactions between them. These results suggest that the same transcription factors may have functional roles in genital development in both sexes, and further investigation into their regulation may provide genetic reagents for future work on morphogenesis, sexual dimorphism, and co-evolution.

Drosophila appendages have long been powerful model systems to study how gene regulatory networks operate during development. Recent work has illuminated the cellular mechanisms underlying the formation and evolution of genital structures in females and males: changes in ovipositor morphology have been traced to cellular expansion and rearrangements in females (Green et al., 2019), while the male posterior lobe forms as a result of extreme cell elongation mediated by the apical extracellular matrix (Smith et al., 2020). Transcription factors like *Pox neuro* and *Sox21b* have been functionally characterized in male genital structures (Glassford et al., 2015; Ridgway, Hood, et al., 2024), but their role in females is comparatively understudied. As we measured the 3D expression patterns for 29 transcription factors in female samples, future work may investigate how those genes regulate their target effector proteins to coordinate the complex cellular processes operating during genital development.

Of note, our gene expression signals are currently restricted to a single time point during the pupal stage in both sexes, and these expression patterns likely shift as development progresses (Vincent et al., 2019). The functional role of these transcription factors on genital development in both sexes can be assessed using genetic perturbations (Ridgway, Figueras Jimenez, et al., 2024; Tanaka et al., 2022), although these experiments may be hindered by the limited number of drivers that have been characterized in these tissues (Chatterjee et al., 2011; Green et al., 2019; Ridgway, Figueras Jimenez, et al., 2024; Ridgway, Hood, et al., 2024; Robinett et al., 2010). Genetic experiments may also confirm putative regulatory interactions that we have inferred by observing mutually exclusive or overlapping gene expression patterns. Finally, a complete picture of the gene regulatory landscape in female genital tissues is possible through single-cell transcriptomics, and this current atlas may prove useful for interpreting such data by providing marker genes for annotating cell populations after tissue dissociation.

This study shows that many transcription factors that have been found to pattern the male terminalia also pattern the female terminalia, even though the adult structures take on radically different morphologies. Future work will be required to determine how the same regulatory genes produce these divergent body parts. The imaginal discs that produce most appendages of the fly share serial homology. Homeotic transformations can cause both large and small shifts in morphology. Legs can be transformed into antennae (Percival-Smith et al., 1997), and forelegs can be transformed into midlegs through the same homeotic process (Lewis et al., 1980). In contrast to all other fly imaginal discs, the genital disc is composed of primordia from three different abdominal segments, each of which forms separate compartments. Abdominal segment 10 produces the analia of both males and females, while Abdominal segment 9 will produce the genitalia in males and is reduced to only form the accessory glands in females (Keisman et al., 2001). Abdominal segment 8 produces the genitalia in females and is reduced to only the epandrial anterodorsal phragma in males (Keisman et al., 2001). Comparing directly homologous abdominal segments between the sexes is therefore unlikely to be fruitful due to these extreme reductions. Determining top-tier patterning genes that establish the unique genetic cascades in the male and female genitalia will help us better understand if their level of serial homology is high, like forelegs and midlegs, or low, like legs and antennae.

One of our most striking findings is the parallel and perpendicular expression patterns of the male and female analia, respectively. We find two transcription factors that pattern the dorsal (*en* and *inv*) and ventral (*E5* and *ems*) regions of the analia of both sexes (Figure S6). Conversely, even though the plates of the analia are divided along different axes – horizontally for females and vertically for males – nine transcription factors show specific expression in these corresponding regions (Figure S6). These results indicate that these two structures may be homologous even if they are found in perpendicular planes. Therefore, we suggest that the region between the anal plates in males also be referred to as the paraproct. It remains to be seen at what point the male analia and female analia form these divisions and how this orientation is rotated between the two sexes. There are also transcription factors that pattern the paraproct in only one sex: *retn* is expressed in females, while *esg* is expressed in males. These differences in expression may indicate sex-specific differences in tissue organization or morphology. The stark change in the orientation of the analia makes the paraproct an exciting model of sexual dimorphism. Furthermore, the males of some species, such as *Drosophila ananassae*, show a division between the dorsal and ventral regions of their cercus, which raises the question of whether the horizontal or vertical orientation is ancestral (Bock, I.R. & Wheeler, M.R., 1972; Polak & Rashed, 2010).

As the knowledge gap between female and male terminalia closes, we will gain a more nuanced view of how these two tissues are connected to one another developmentally. A recent study found that the transcription factors *C15* (also analyzed in this study), *Al*, and *lim* were all expressed in both female and male terminalia (Ridgway, Figueras Jimenez, et al., 2024). A male-specific enhancer for *C15* was identified, indicating that the regulation of the male and female terminalia expression patterns is independent (Ridgway, Figueras Jimenez, et al., 2024). The separation of female and male regulatory sequences would decrease pleiotropy, allowing for the expression pattern of each sex to evolve independently. Similarly, an analysis of single-nuclei RNA-seq of the adult female reproductive tract revealed that 40% of the previously characterized male-specific seminal fluid genes are expressed in the female reproductive tract (Thayer et al., 2024). The lack of attention to female traits has led us to mislabel genes as male-specific and provided us with an overly simplistic view of sexual dimorphism. We must continue to build resources in female specific systems if we hope to understand how sexually dimorphic traits evolve.

## Materials and Methods

### Drosophila strains

We obtained our lines from the Bloomington Drosophila Stock Center *yellow white,* y^1^w^1^, (Bloomington Stock Center #1495) and *armadillo*-green fluorescent protein (GFP), arm-GFP, (Bloomington stock number #8556).

## Sample Collection and Dissection

Female and male *D. melanogaster* pre-pupa were collected at room temperature and incubated in a petri dish with moisture for 48 hours prior to dissection. The posterior tip of the pupa was then separated, and the terminalia flushed out using a micropipette. The samples were then collected in Phosphate Buffered Saline (PBS) and then fixed in PBT and 4% paraformaldehyde (PFA). Samples were finally washed three times in PBT and stored at 4⁰C for less than 12 hours before being processed for the HCR reaction. Arm-GFP samples were mounted and imaged after fixation.

## Hybridization Chain Reaction (HCR)

Samples were taken from storage and blocked by incubating in antibody buffer for 2h at room temperature. They were then incubated with rat anti-E-cadherin, 1:100 in PBT (DSHB Cat# DCAD2, RRID:AB_528120) overnight at 4°C. The samples were then washed 3 times in PBT, then washed 4X for 15 minutes in PBT and incubated with Molecular Instruments antibodies anti-rat B4 secondary antibody at 1 ug/mL for 2 hours at room temperature. Samples were then washed with PBT 3X and post-fixed for 10 minutes in 4% PFA in PBT before being washed 3 more times in PBT. It was then washed once in 5X sodium chloride, sodium citrate with 0.1% Tween (5X SSCT). Probe solutions made with 8 uL HCR initiator probe set per 250 uL of incubation volume. All HCR probes were designed and synthesized by Molecular Instruments Inc. Transcript accession numbers and sequences that were sent to Molecular Instruments, Inc. for HCR probe design of each gene can be found in Sup Table 2. Samples were prehybridized in probe hybridization buffer for 30 minutes at 37 °C before being incubated in 250 uL of the probe solution for 48 hours at the final hybridization step. Samples are then washed 4 times with probe wash buffer and 3 times with 5X SSCT. B1 488, B3 647, and B4 546 amplifiers were used. h1 and h2 amplifier hairpins are snapped cooled by heating 5 uL of hairpin stock per sample to 95 degrees C for 90 seconds, then cooling for 30 minutes in the dark to room temperature. Samples are then pre-amplified for 30 minutes in an amplification buffer and then amplified overnight in the dark in an amplification solution made of snap-cooled hairpins and amplification buffer. Finally, samples are washed 3 times in PBT and once in 50:50 PBT Glycerol solution. Samples are then moved to glycerol and mounted on microscope slides coated with poly-L-lysine.

## Imaging and Image Processing

Samples were imaged at ×20 on a Leica TCS SP8 confocal microscope. For HCR the expression patterns of two genes and ECAD, which gave only weak expression, were taken for all samples. As the imaged structures are three-dimensional in nature, we used the MorphoGraphX program (Barbier de Reuille et al., 2015) to render and manipulate images in three dimensions. This allowed us to rotate the samples to better present the most informative perspectives of the various phallic structures. Raw image files are available on request.

## Acknowledgments and Funding

We thank Amir Yassin, Troy Shirangi, and the entire Rebeiz lab for their comments on this project and manuscript. This work was supported by the National Institutes of Health grants R35GM14196 (to M.R.) and K99GM147343-01 (to G.R).

## Tables

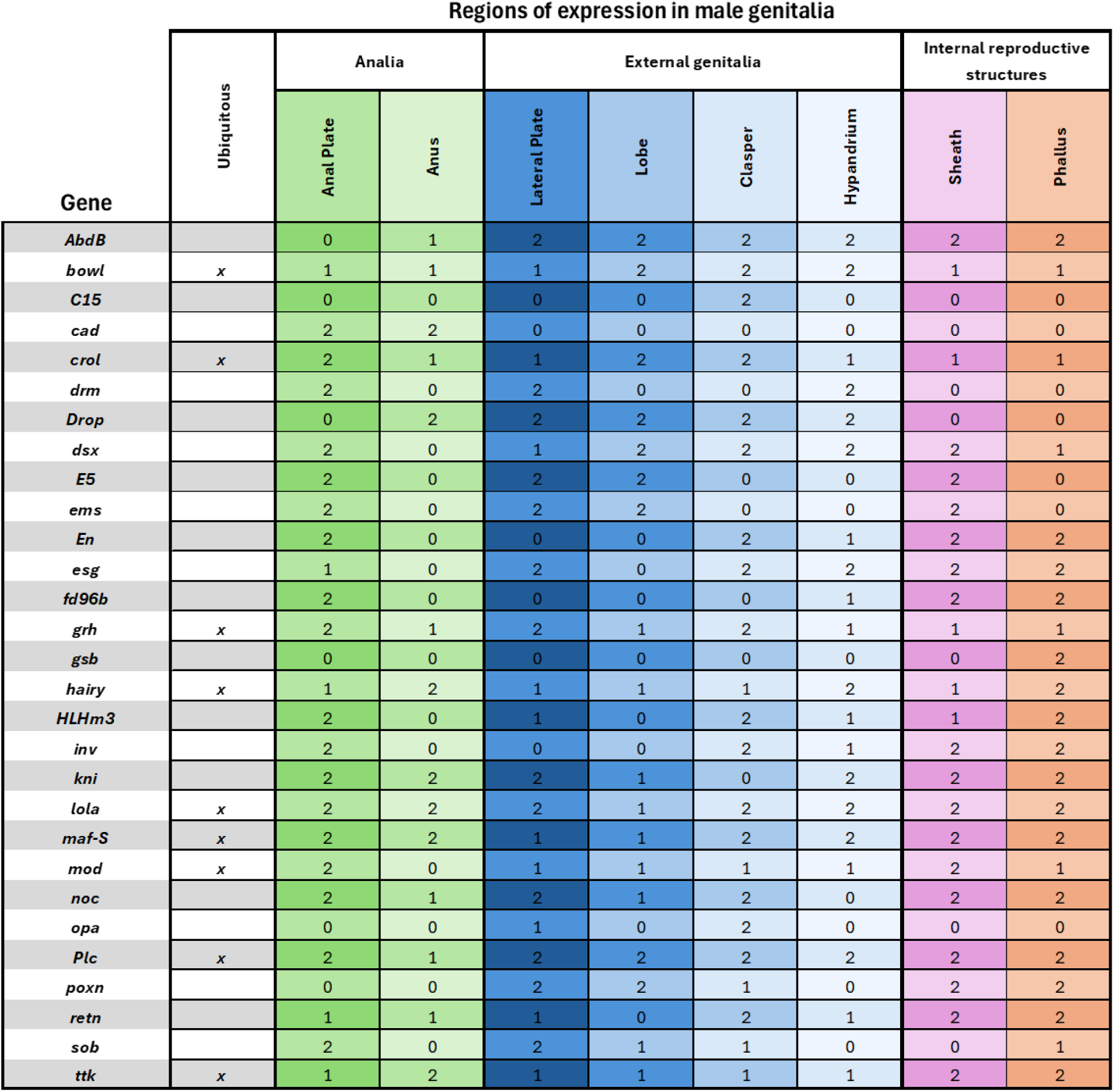
Sup Table 1.

**Sup Table 2:**
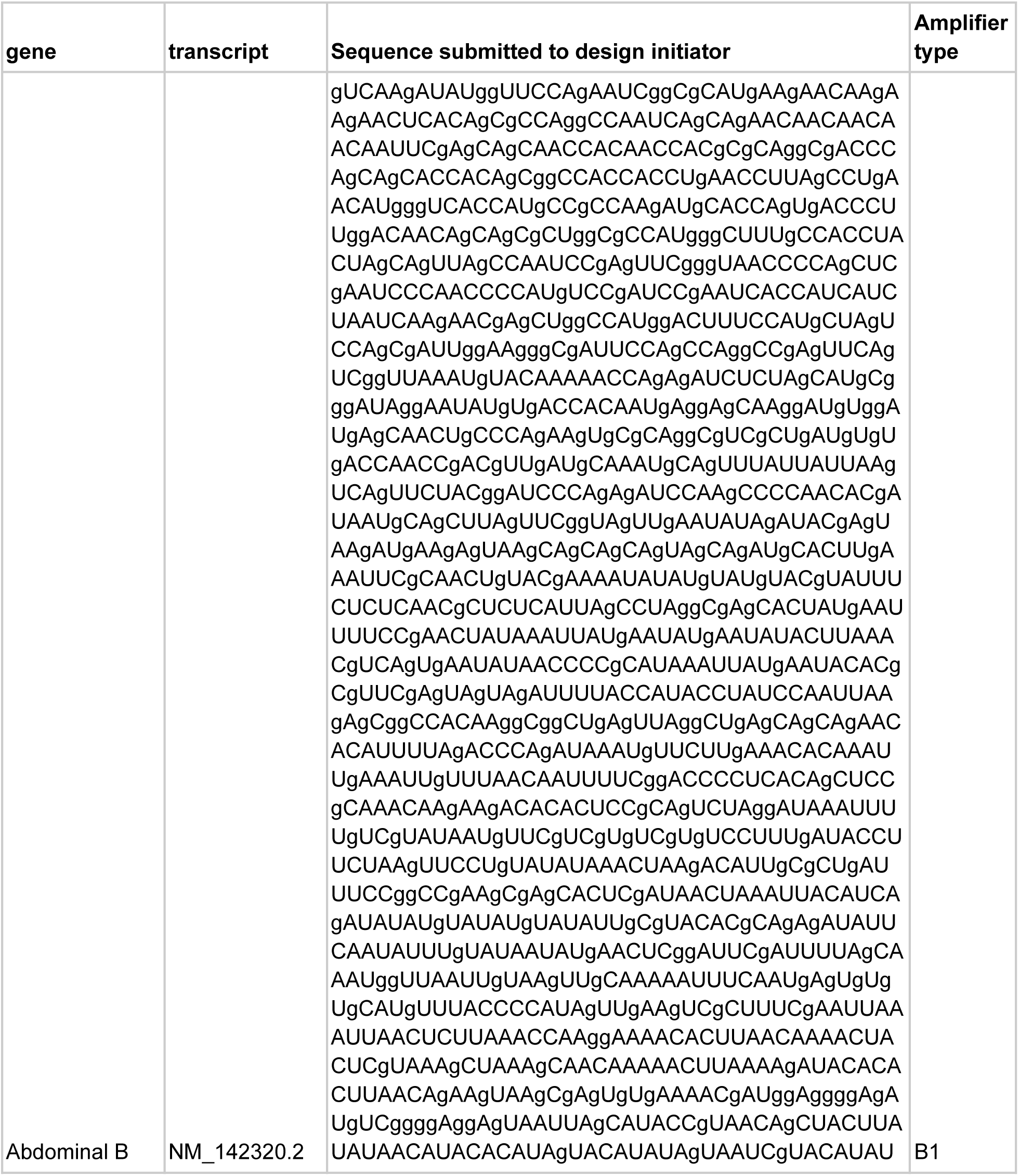
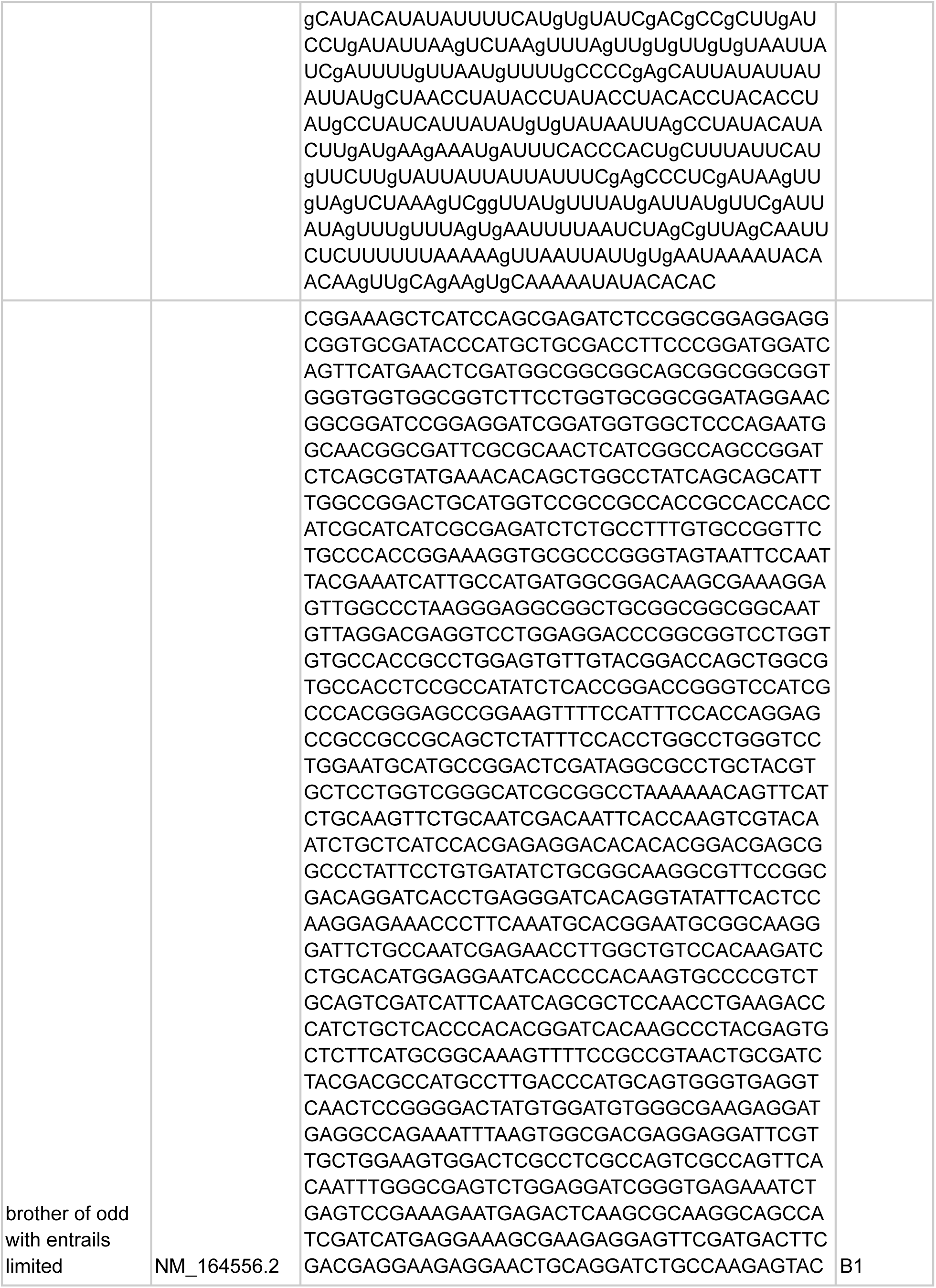
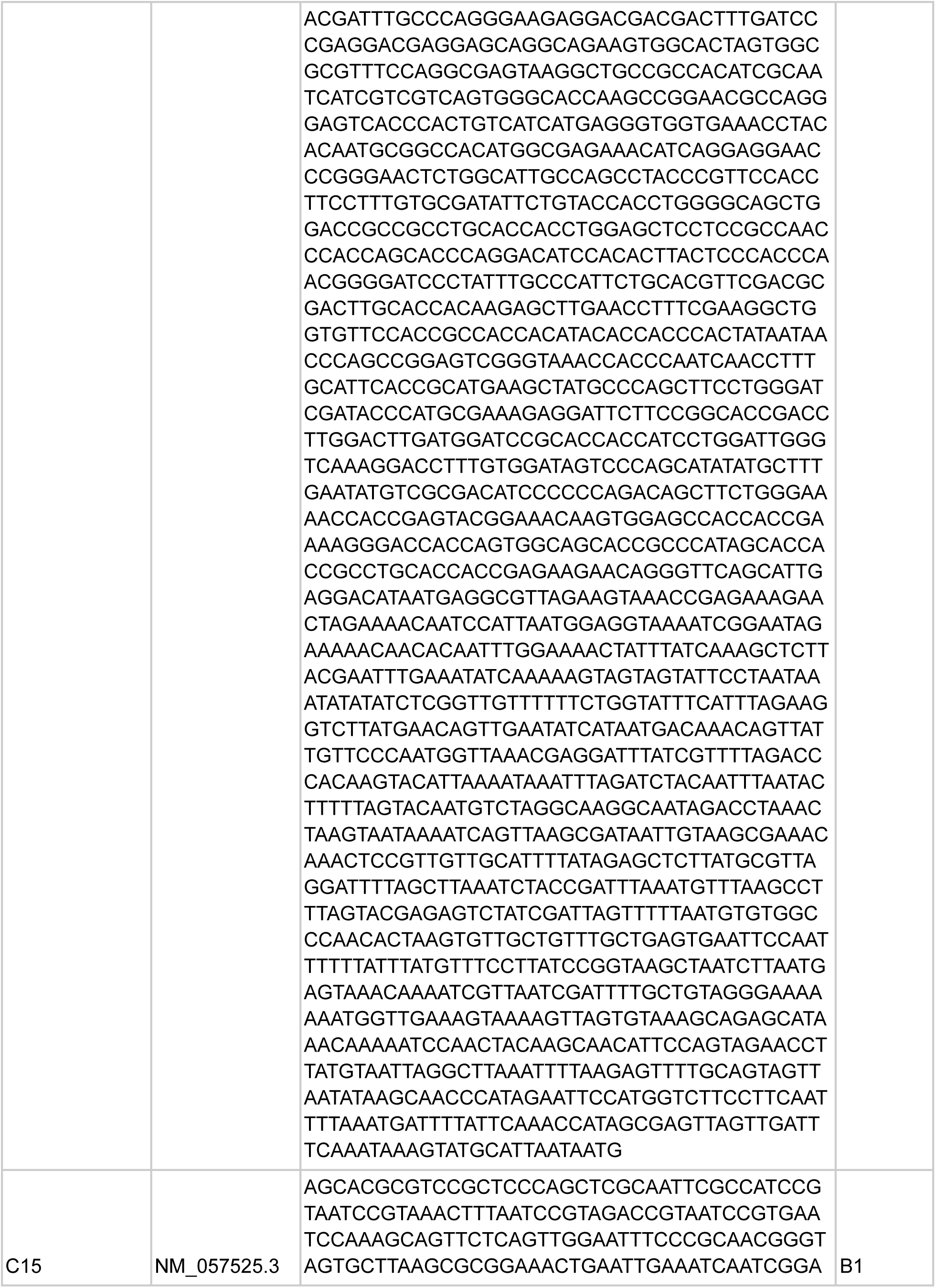
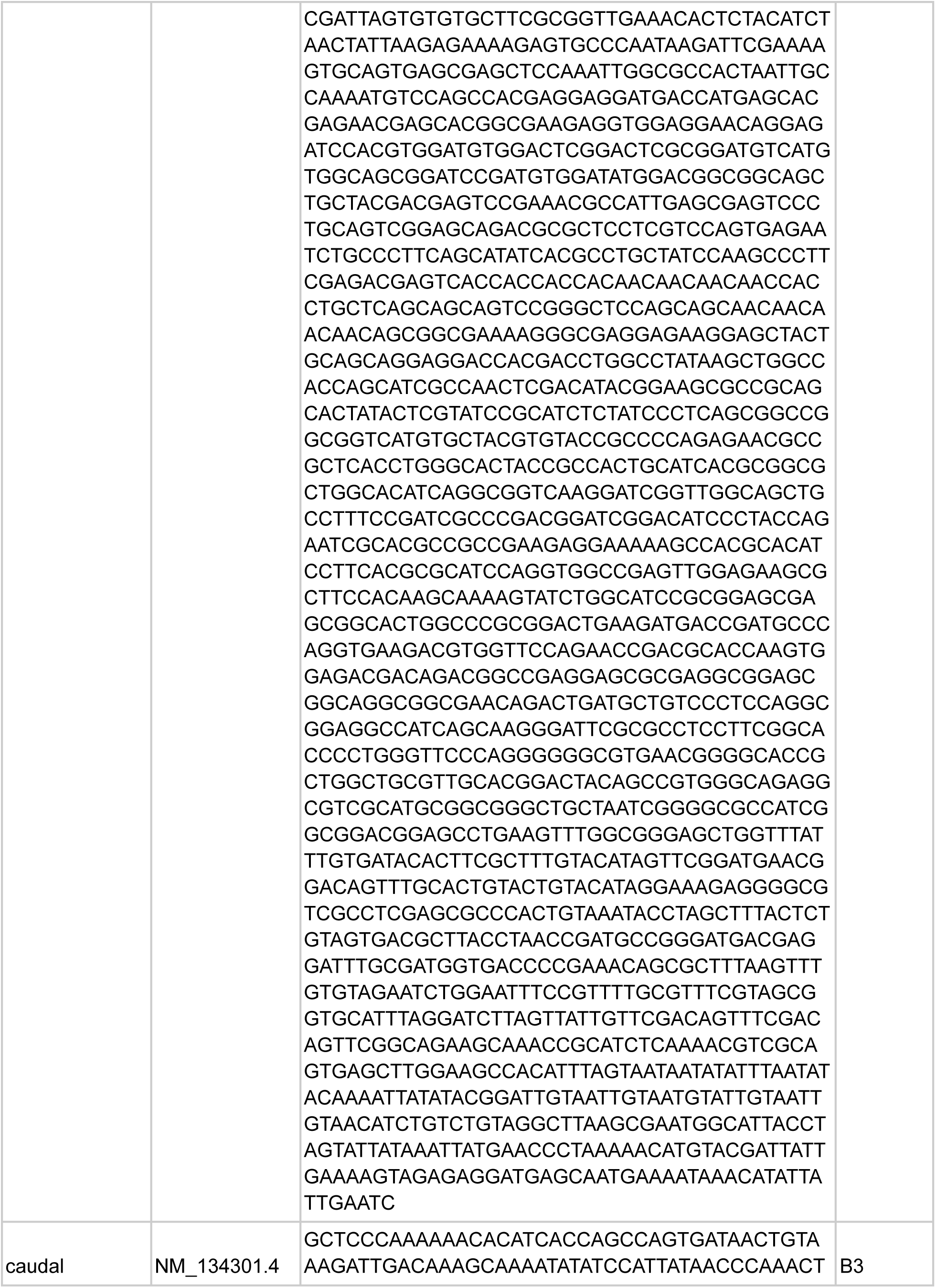
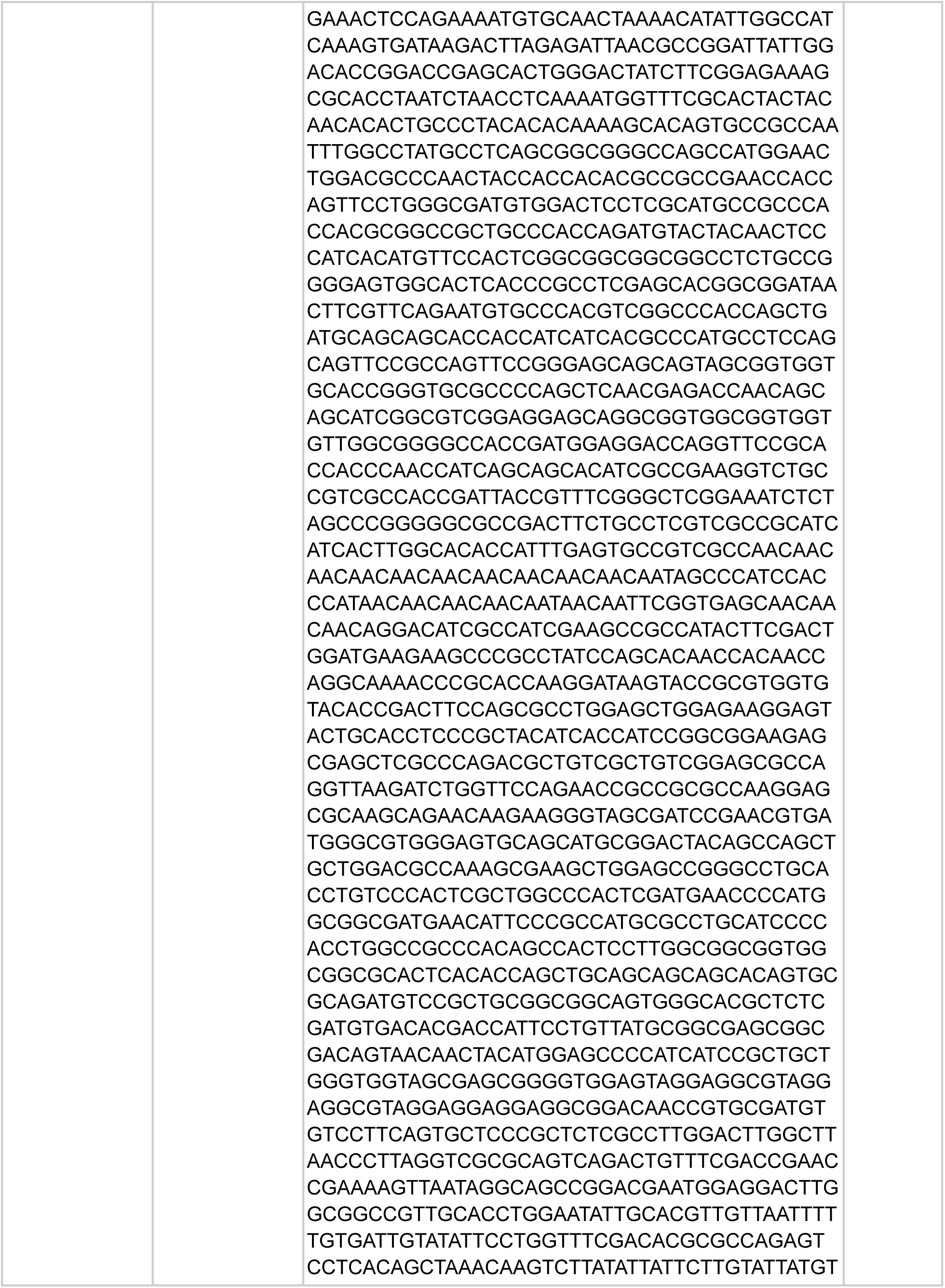
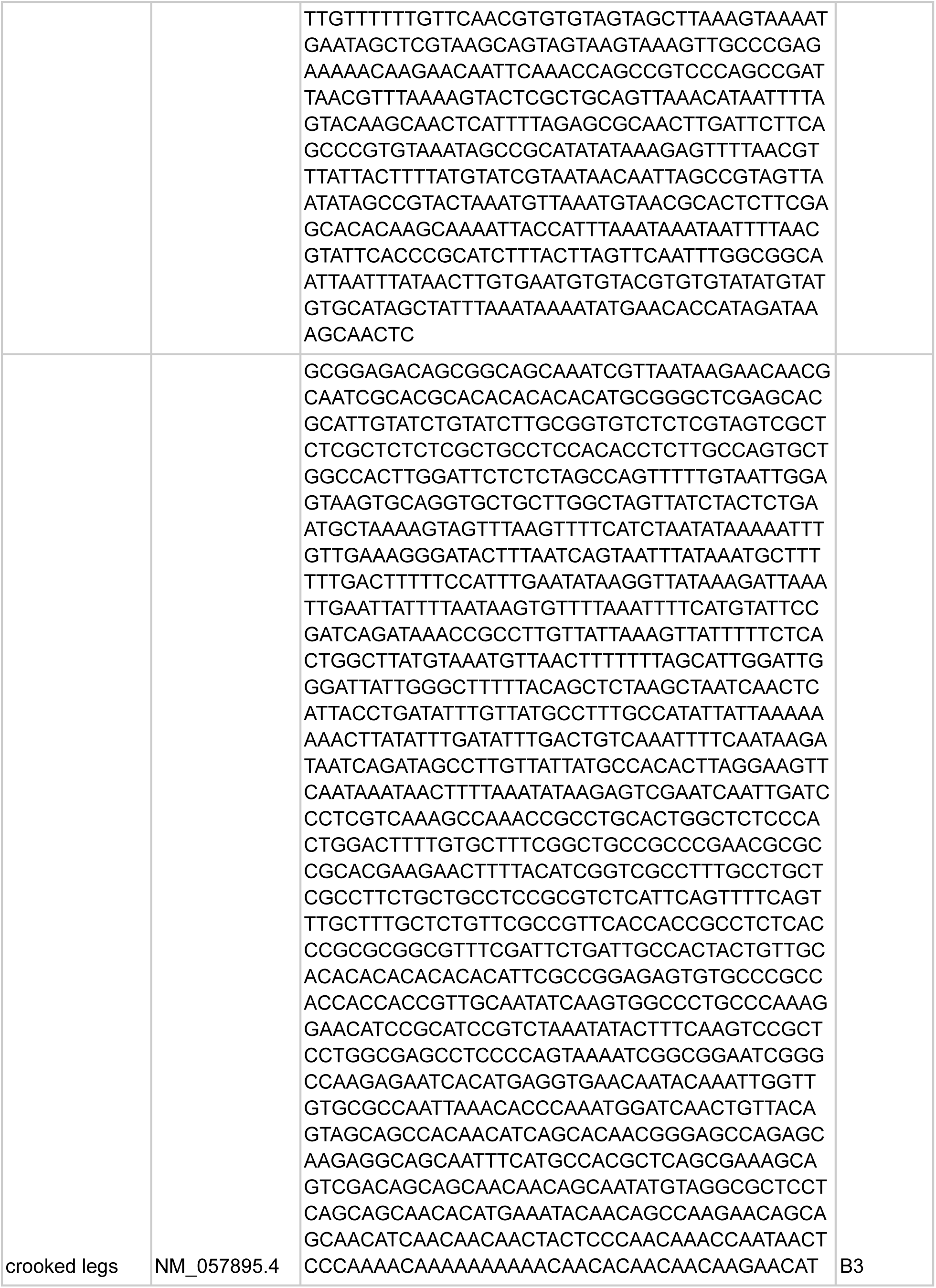
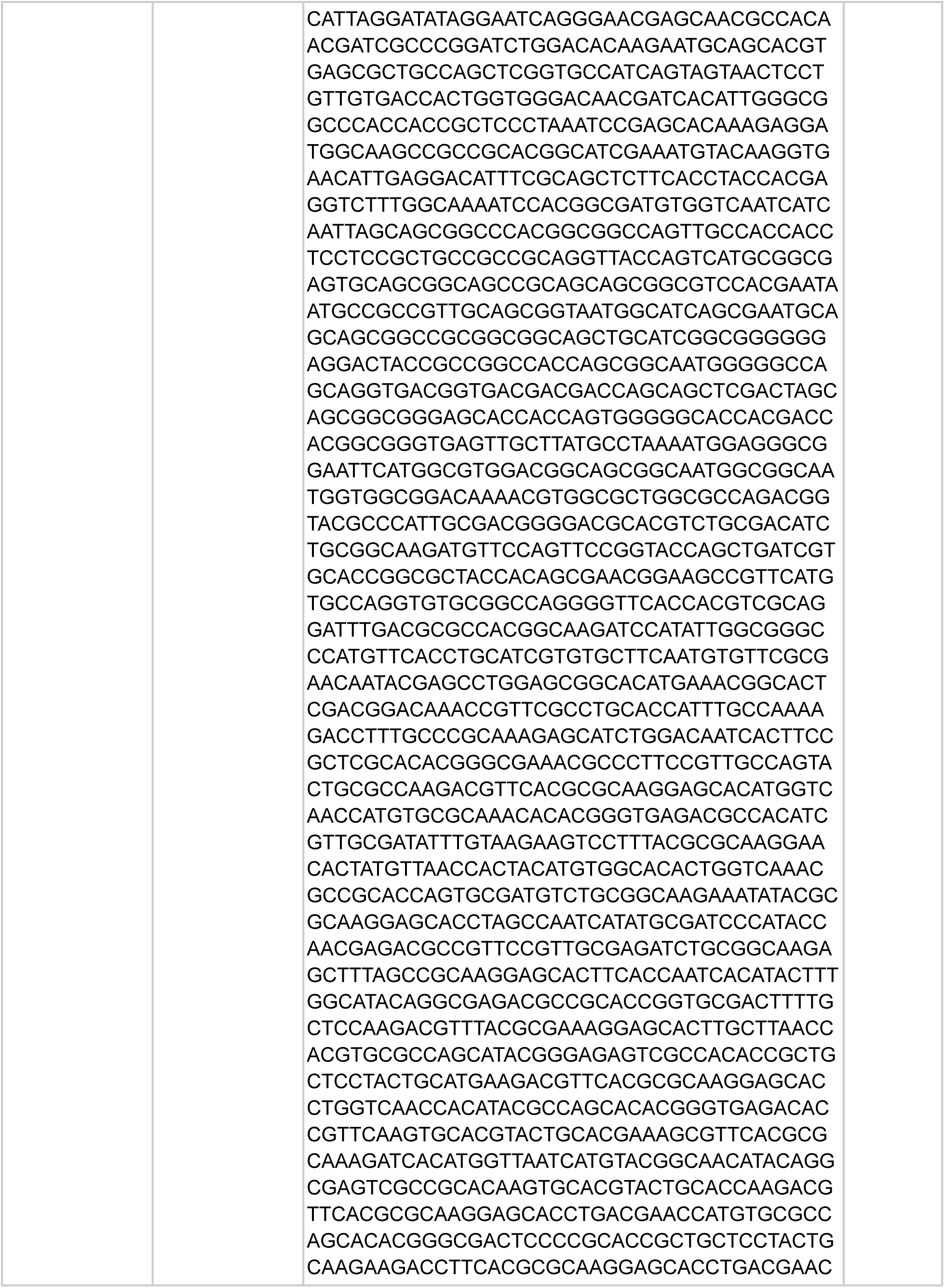
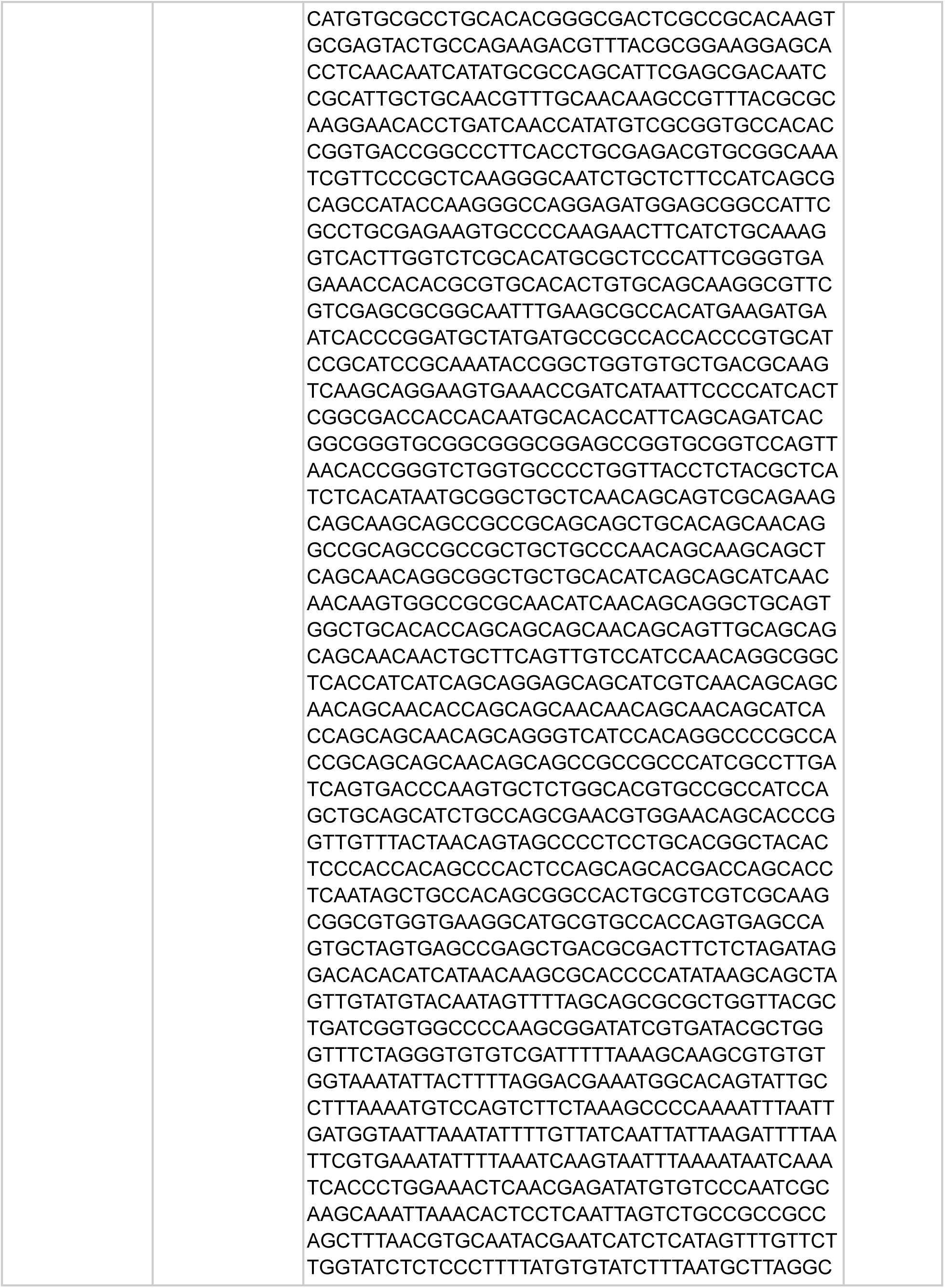
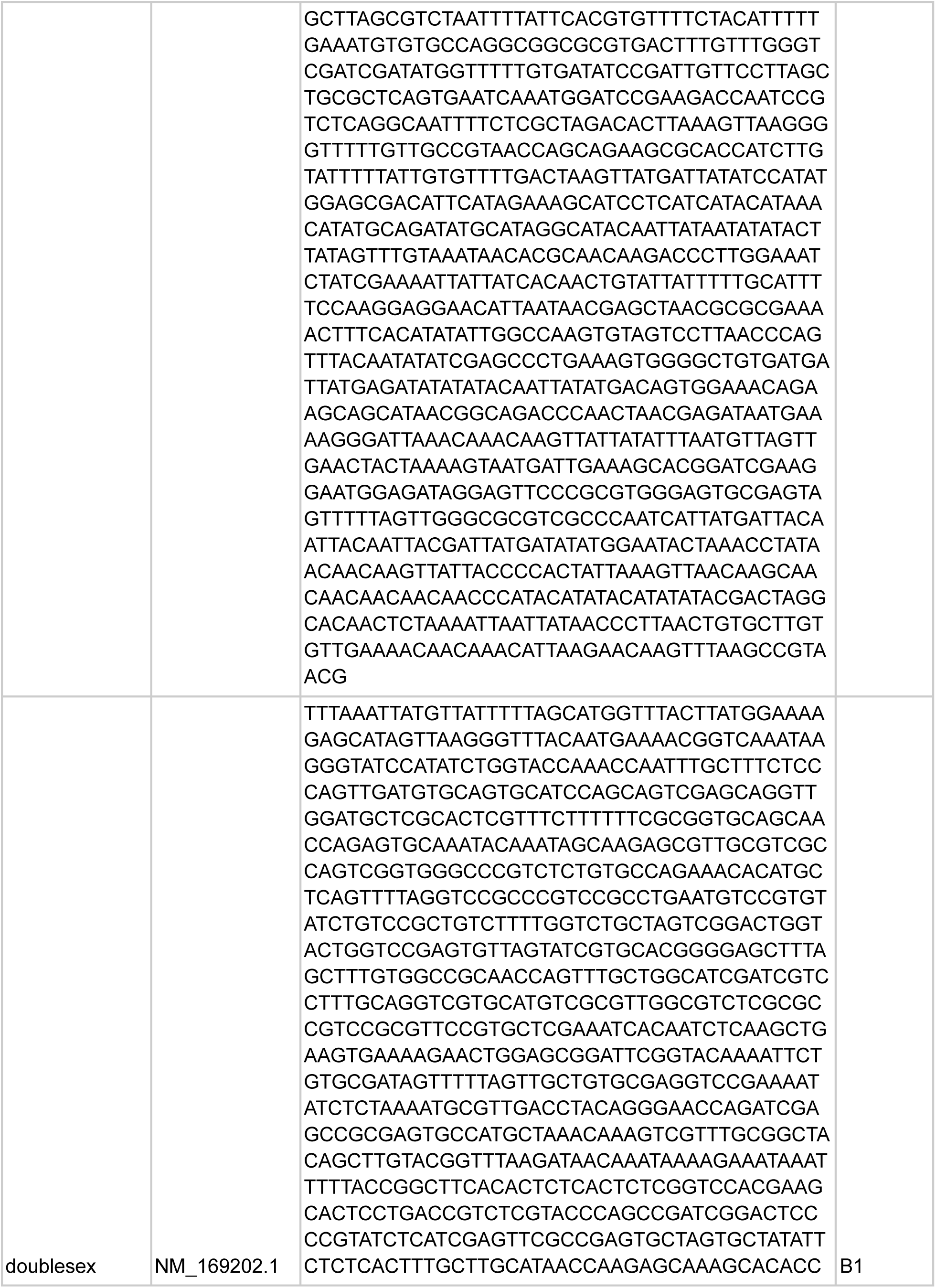
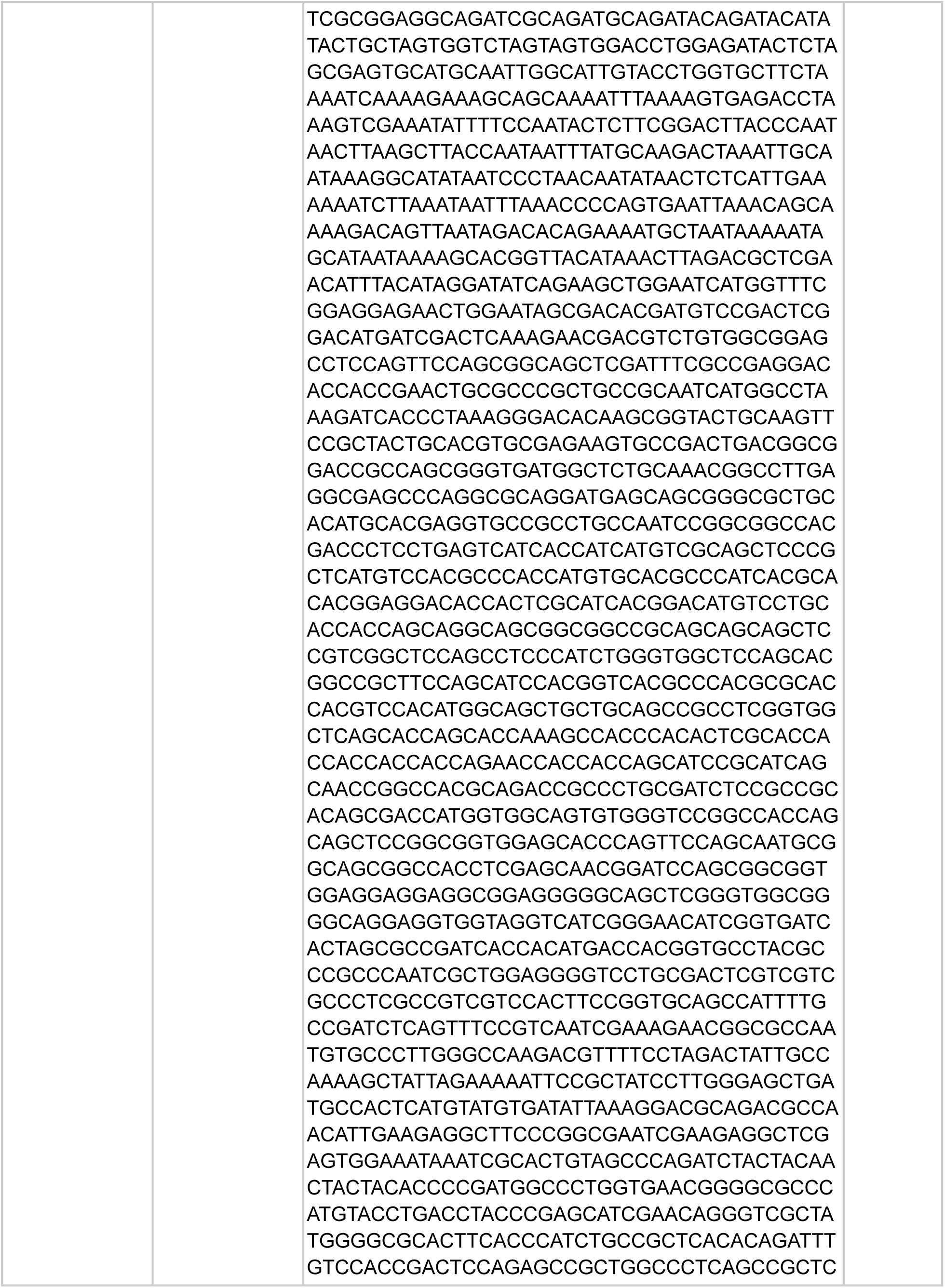
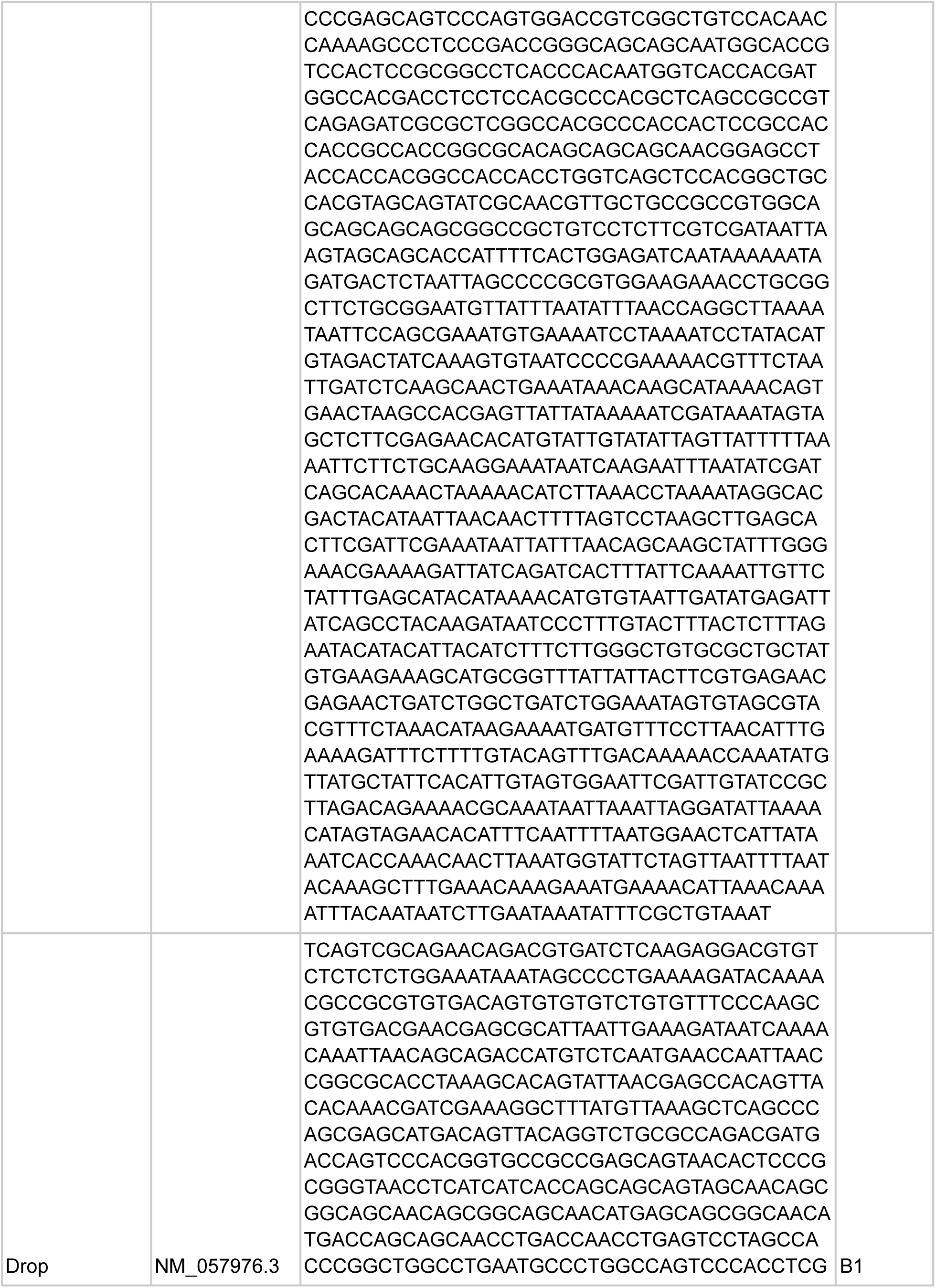
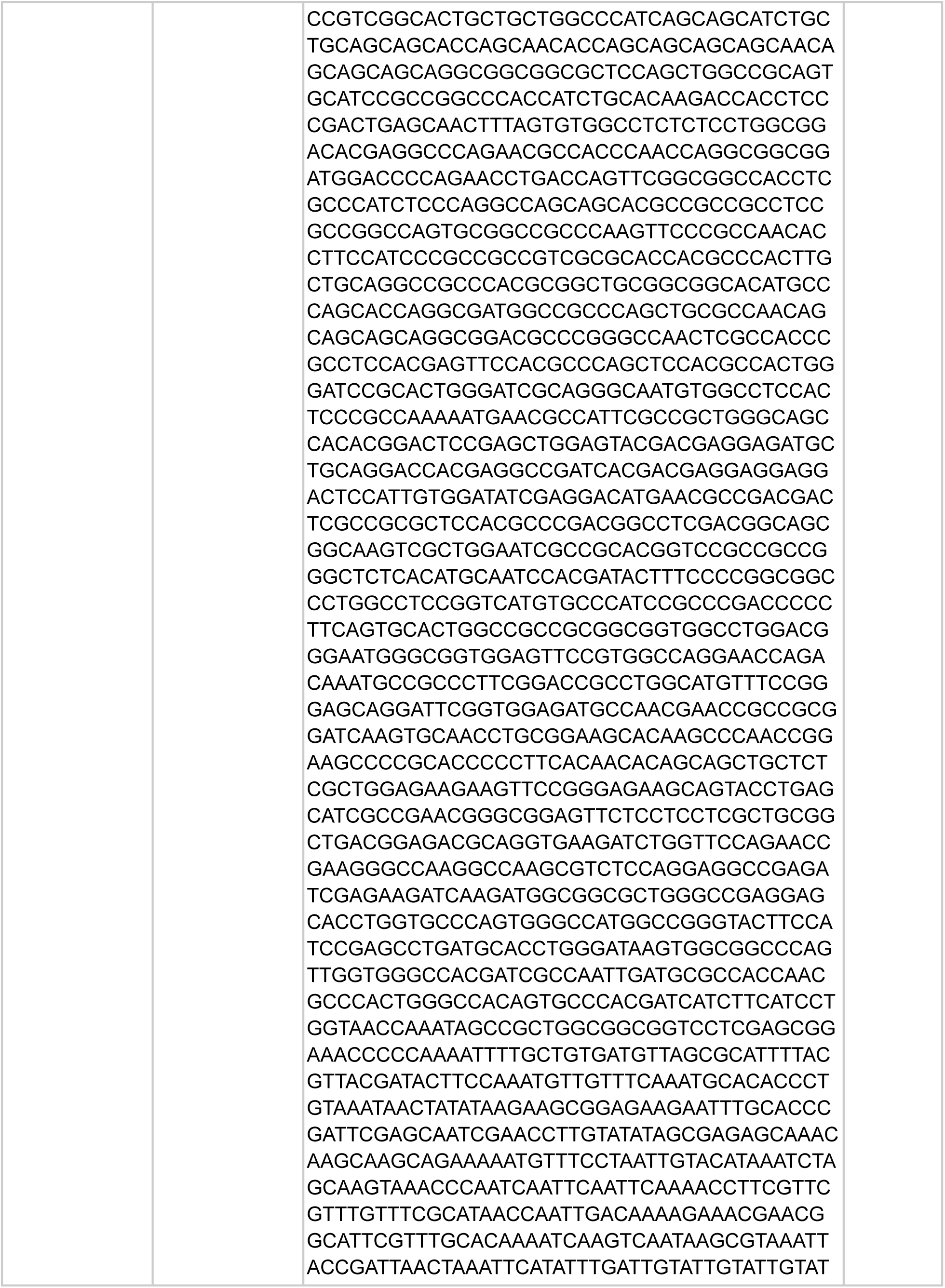
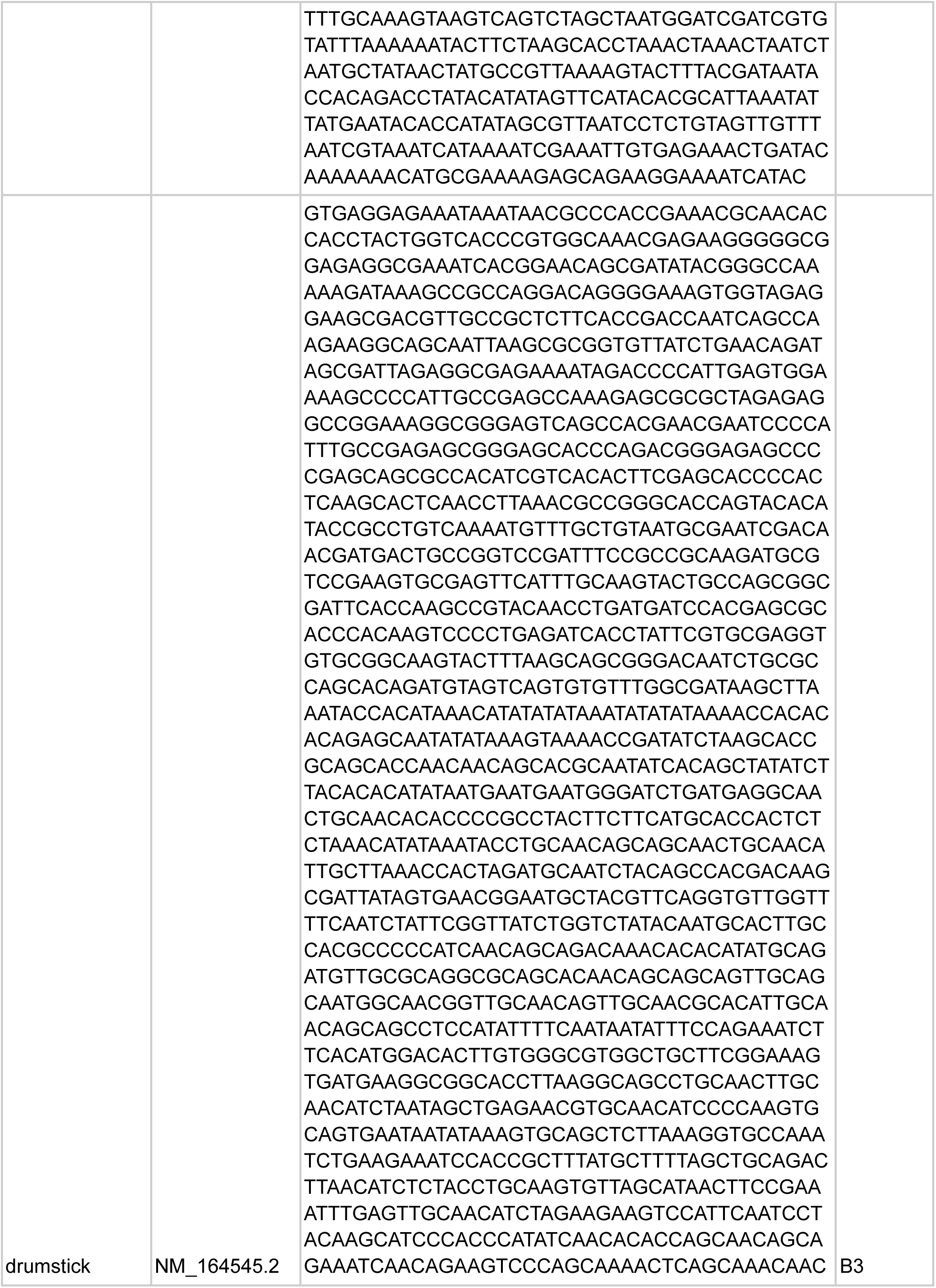
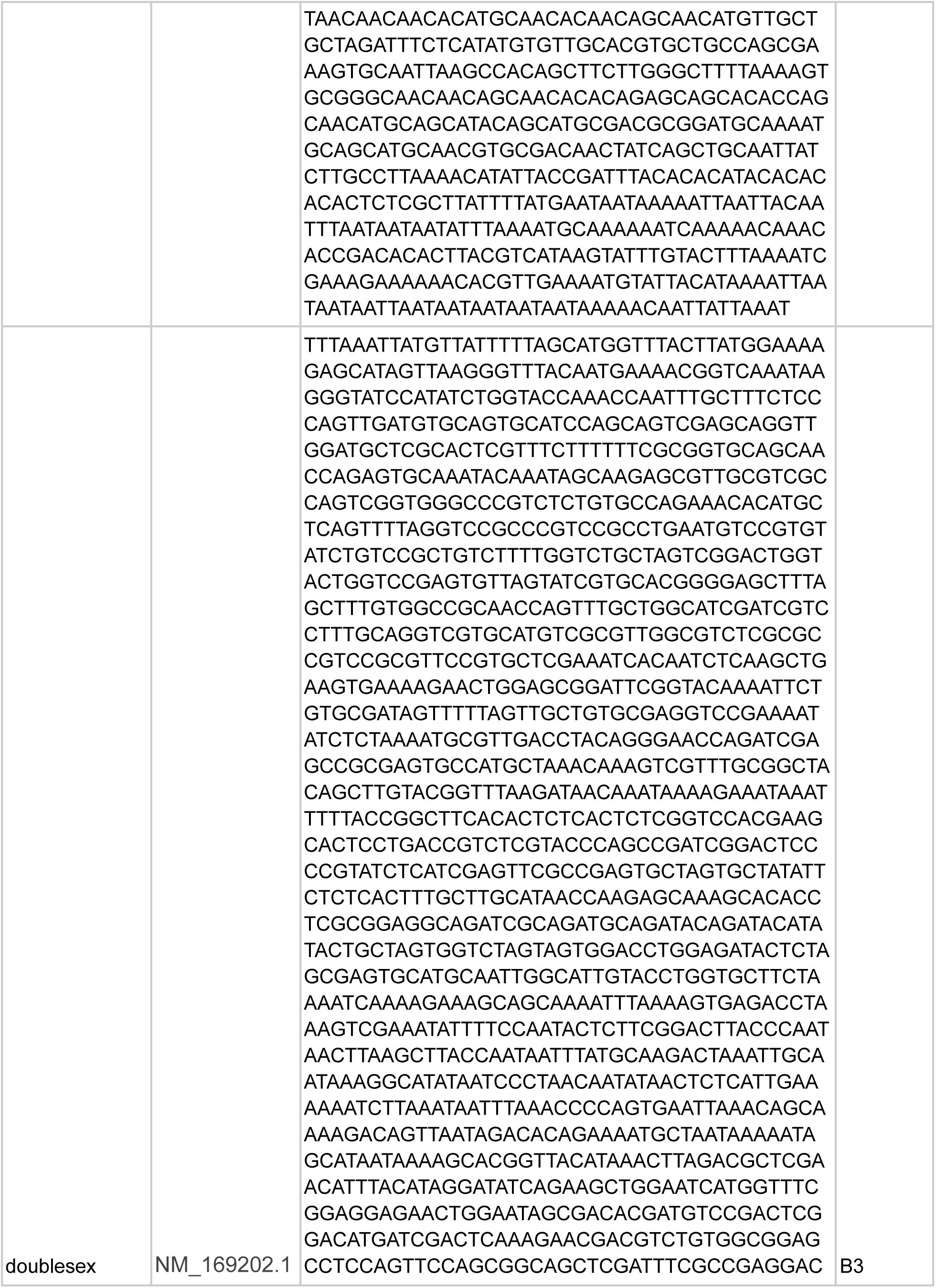
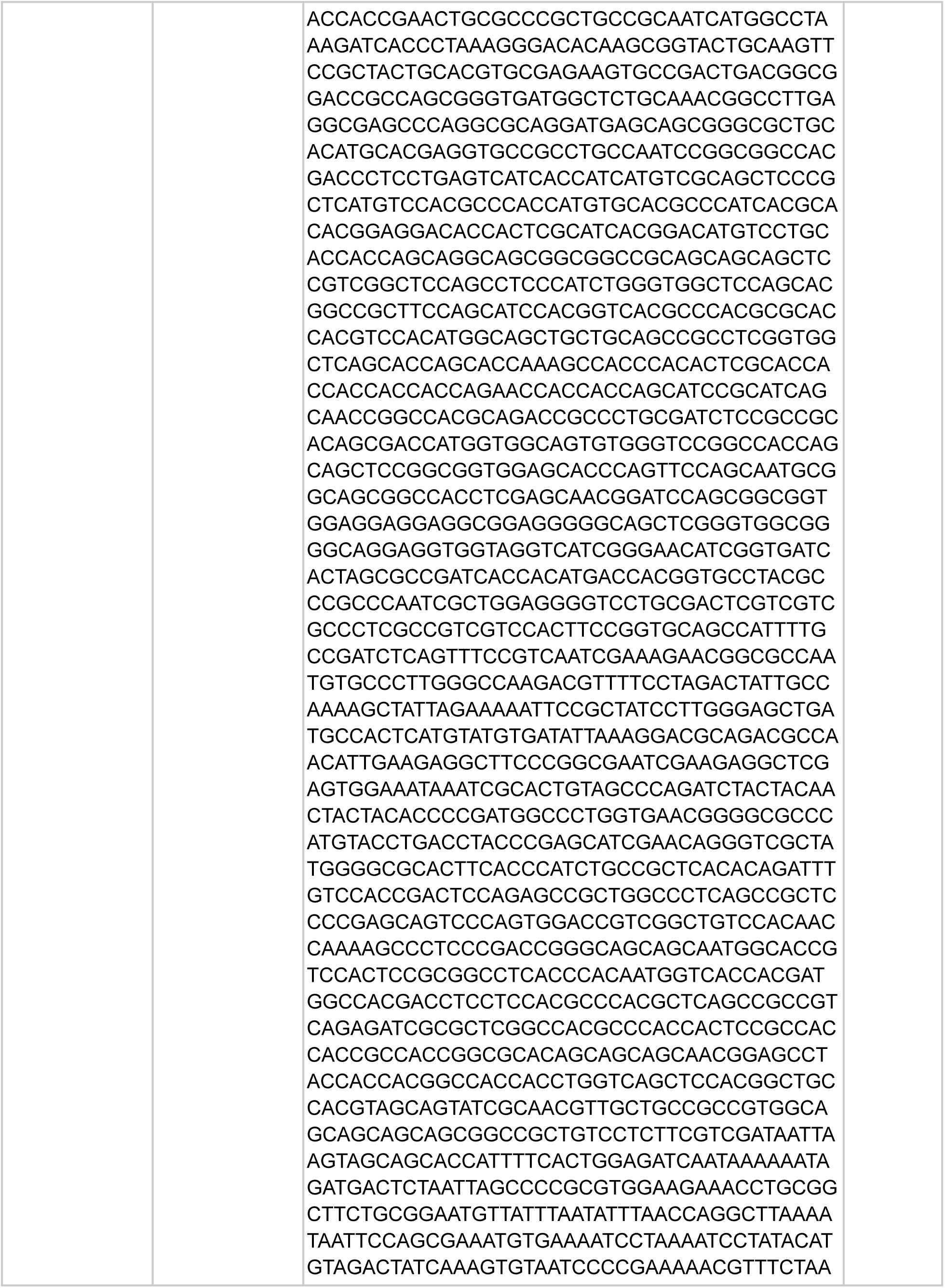
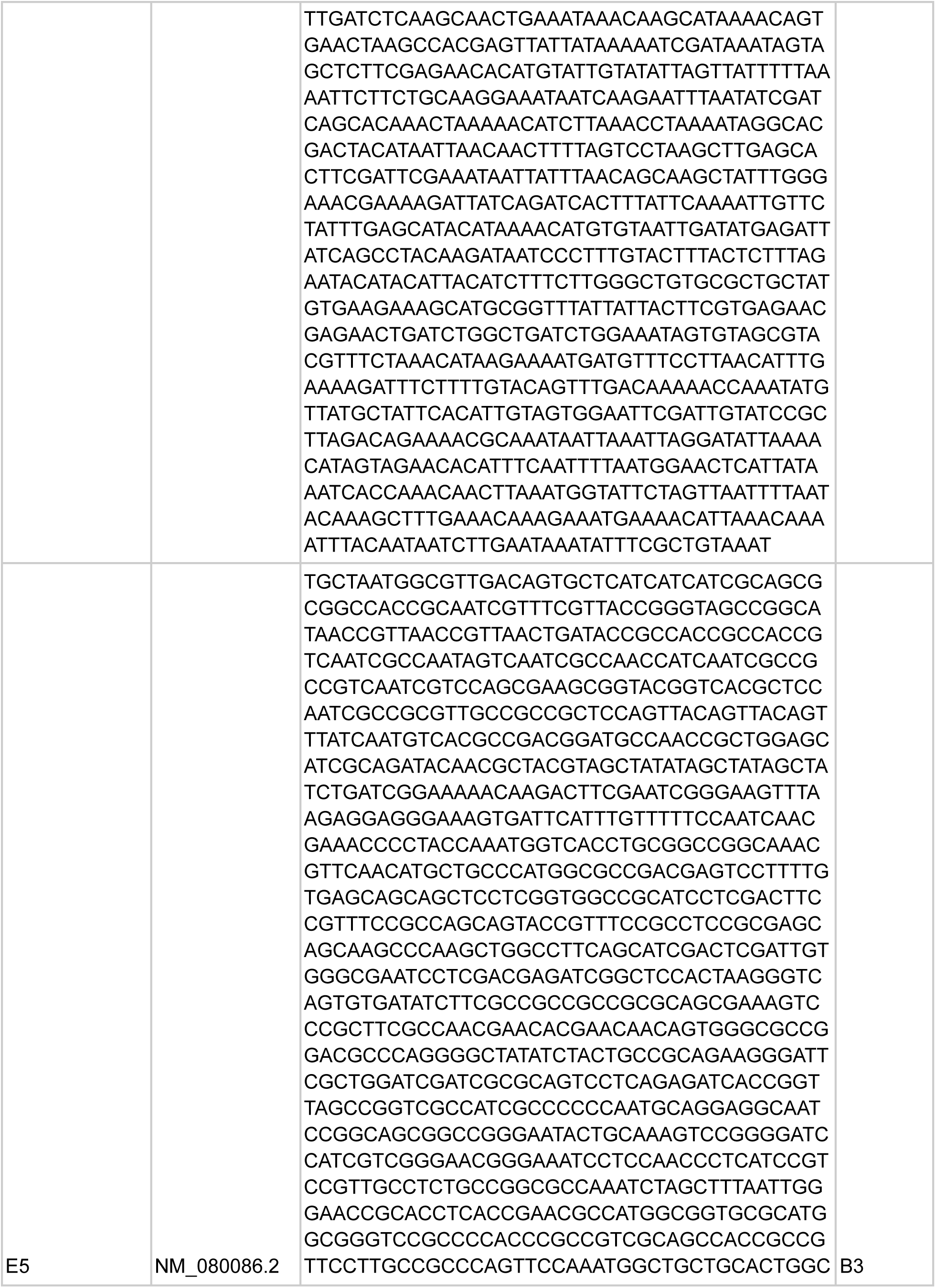
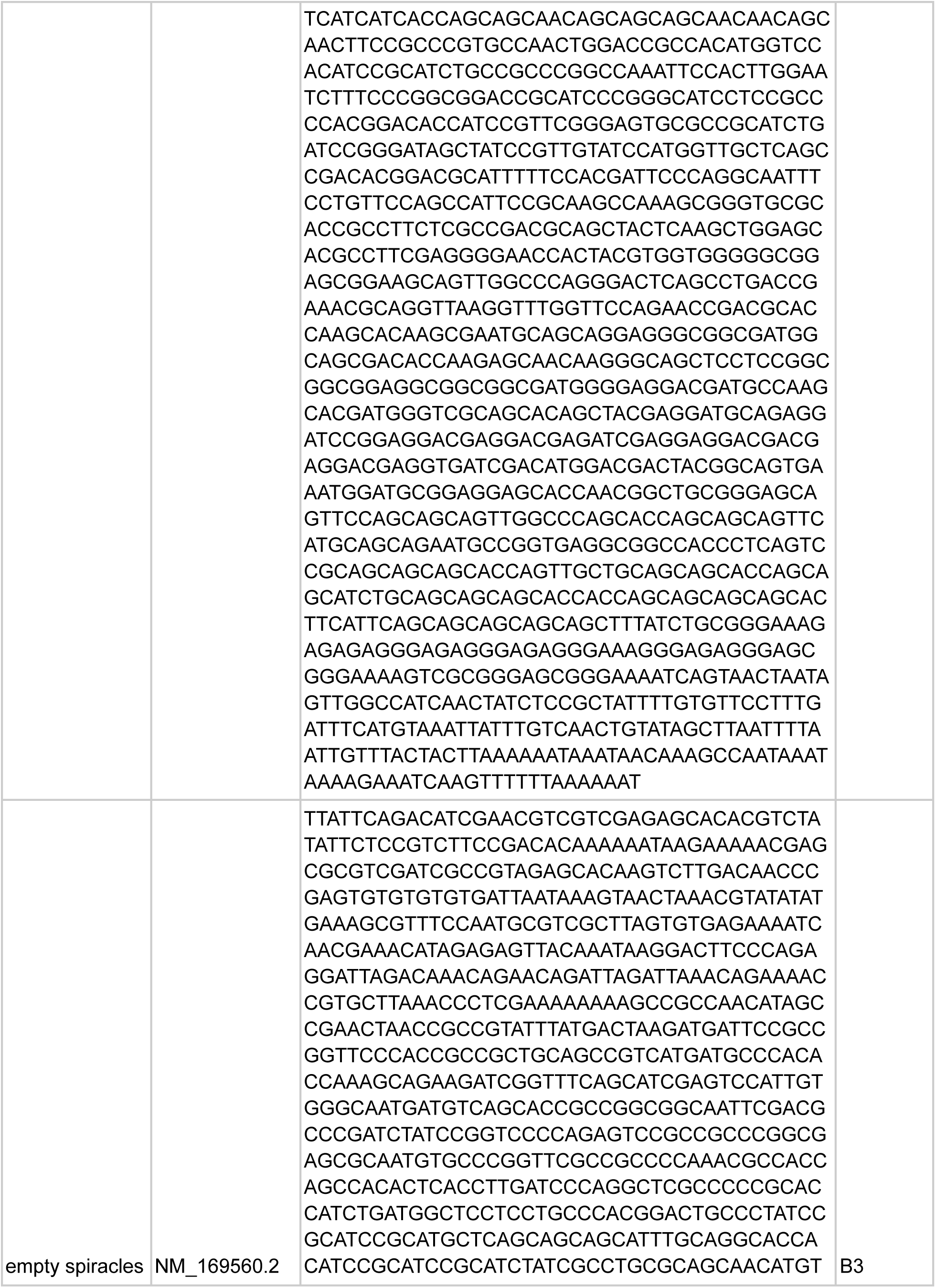
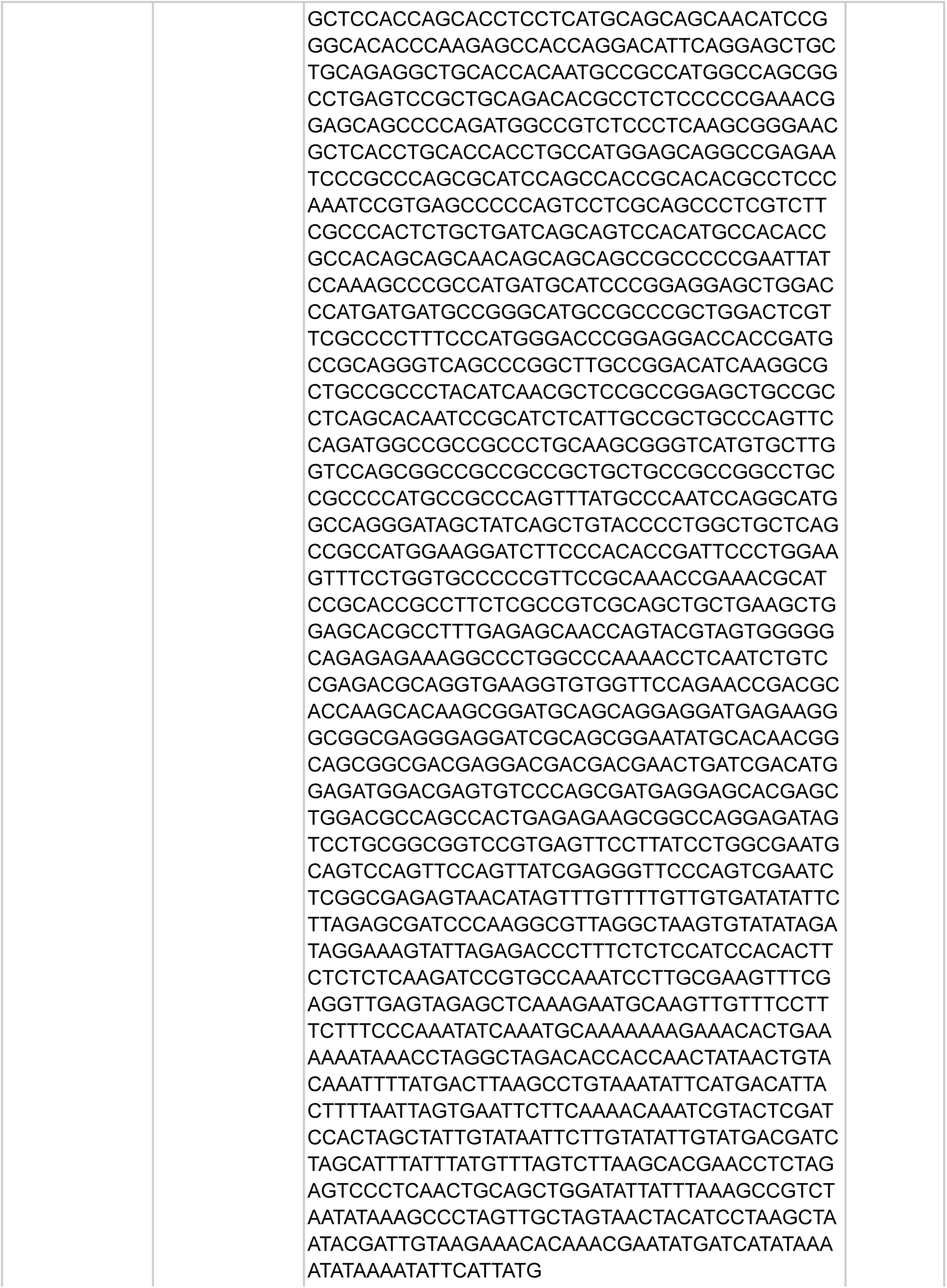
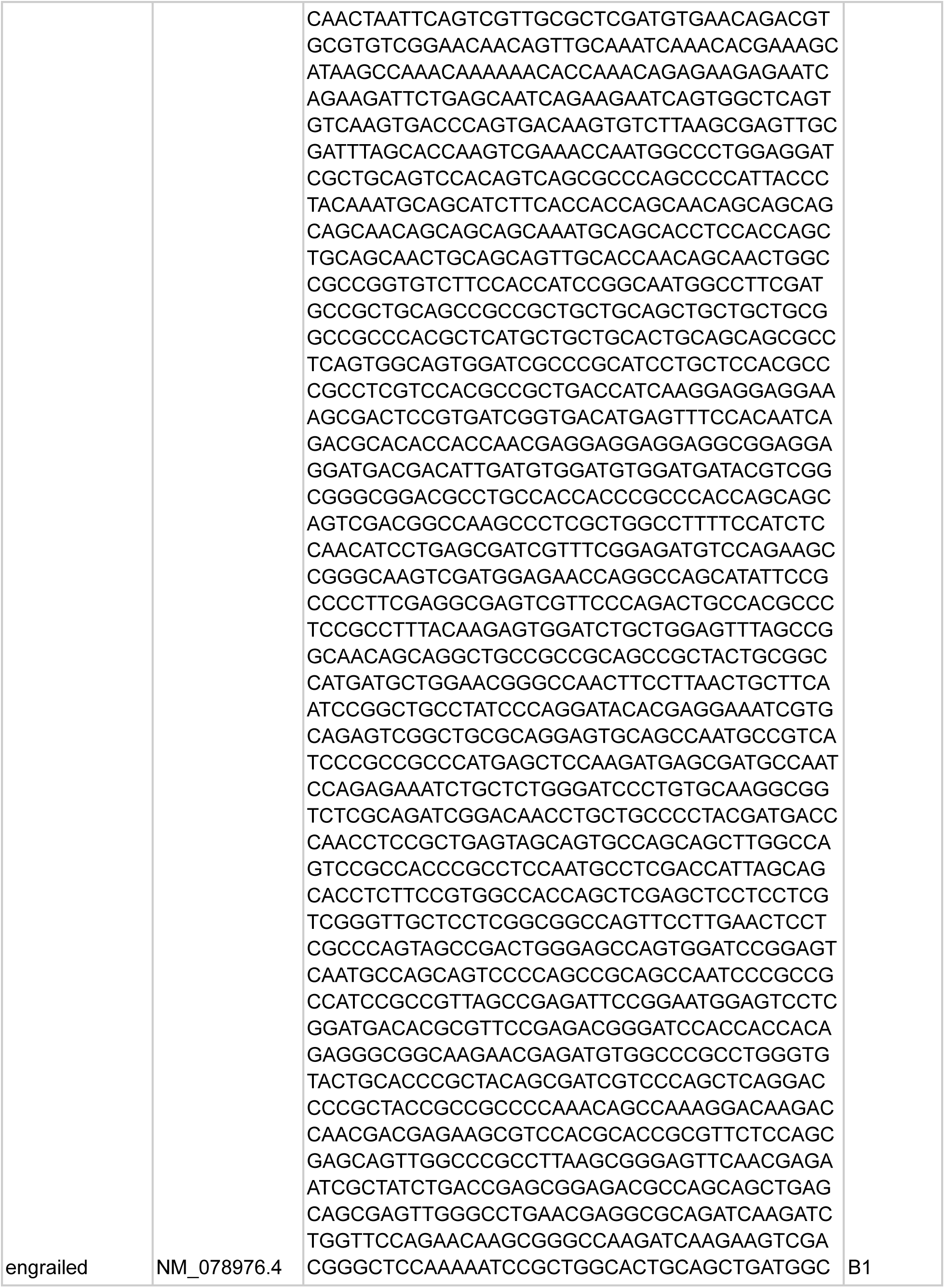
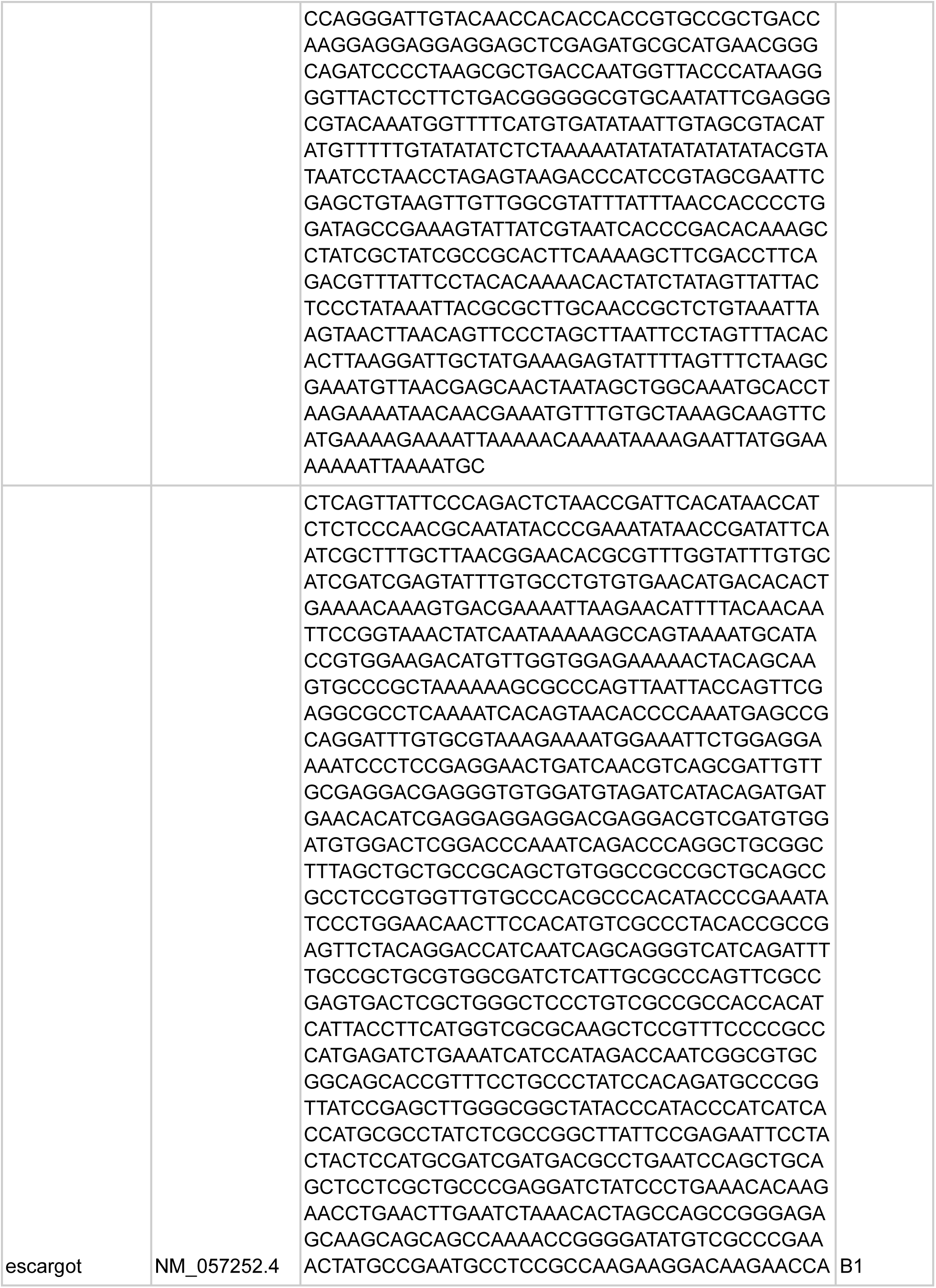
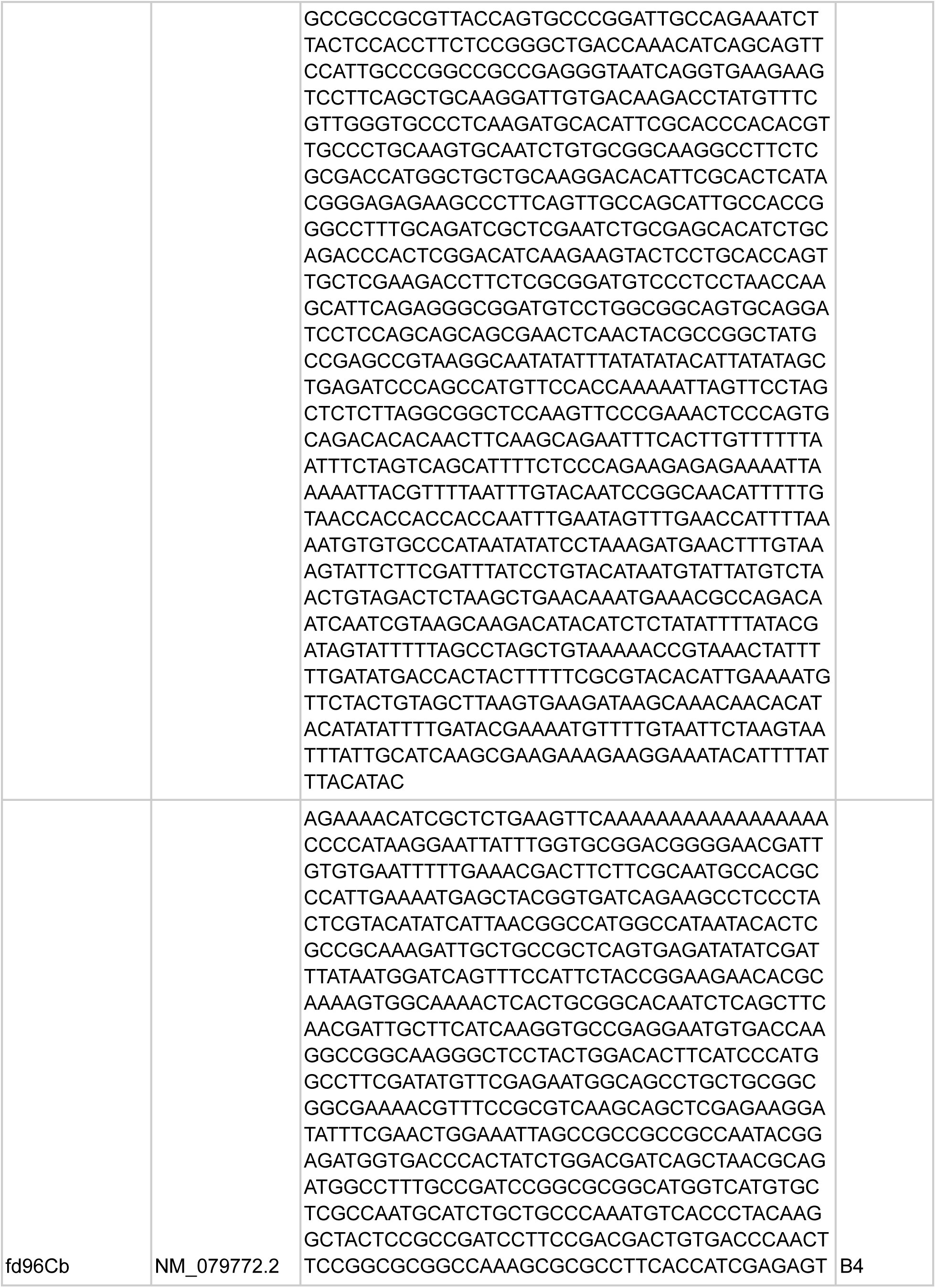
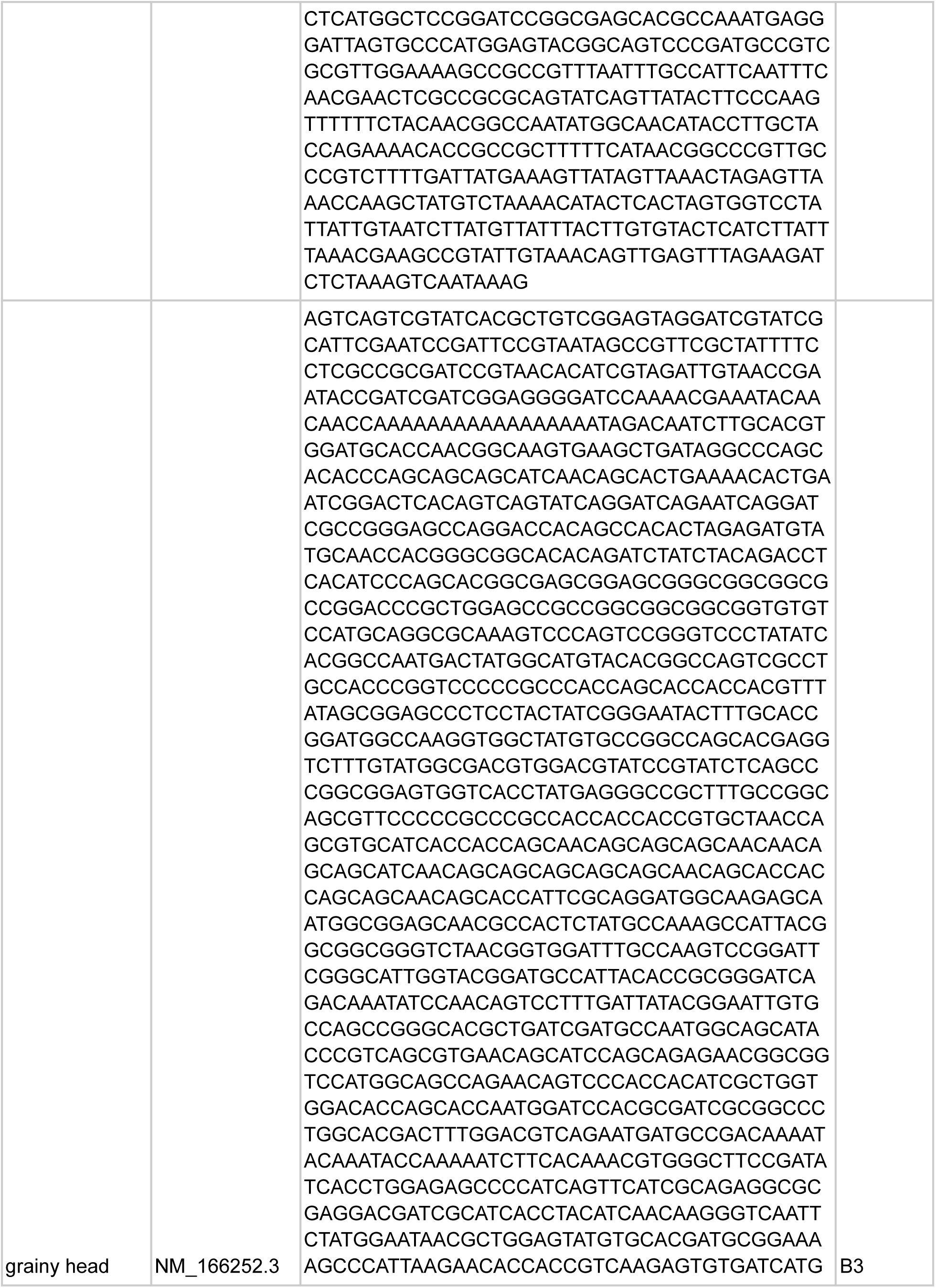
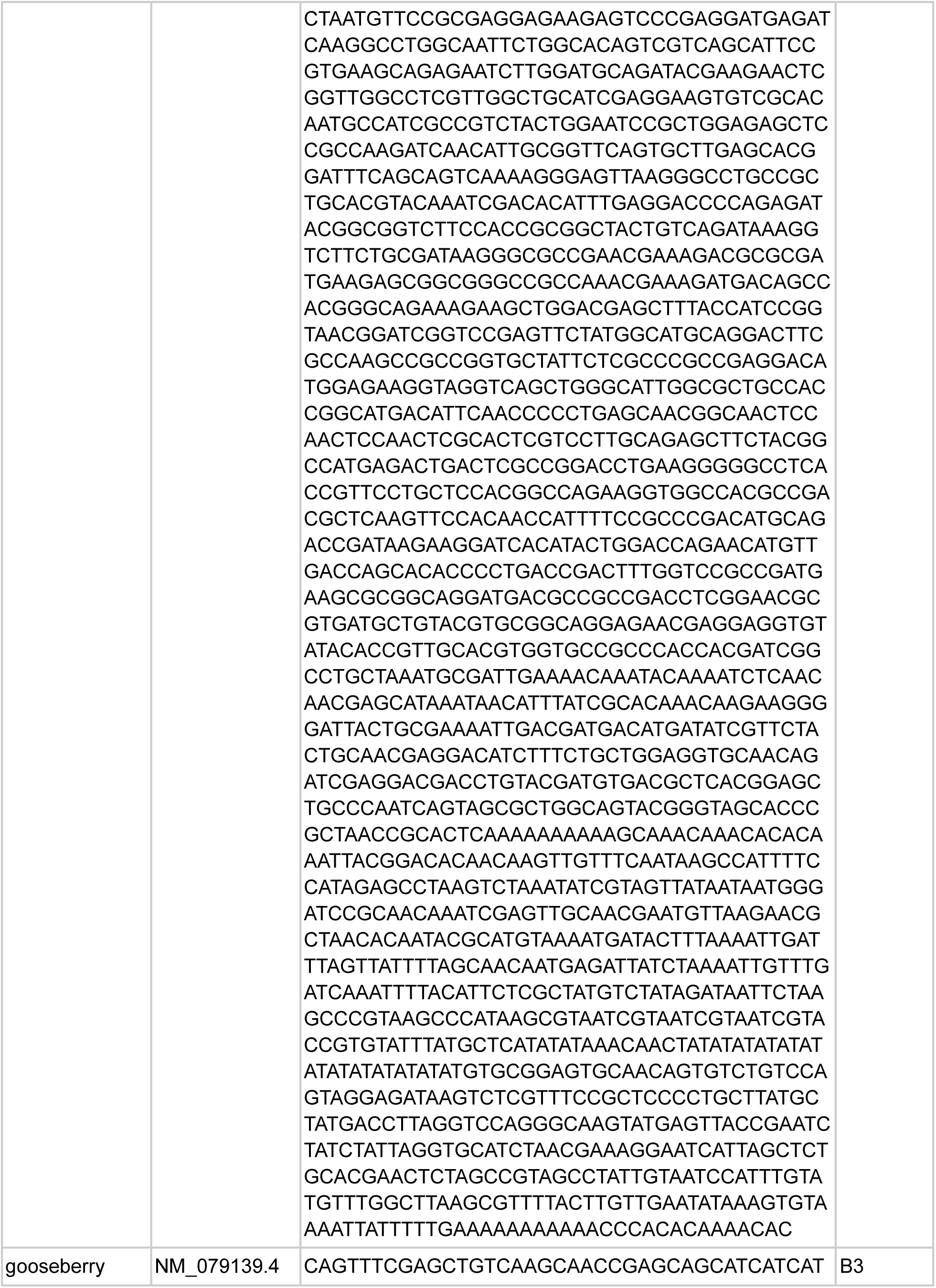
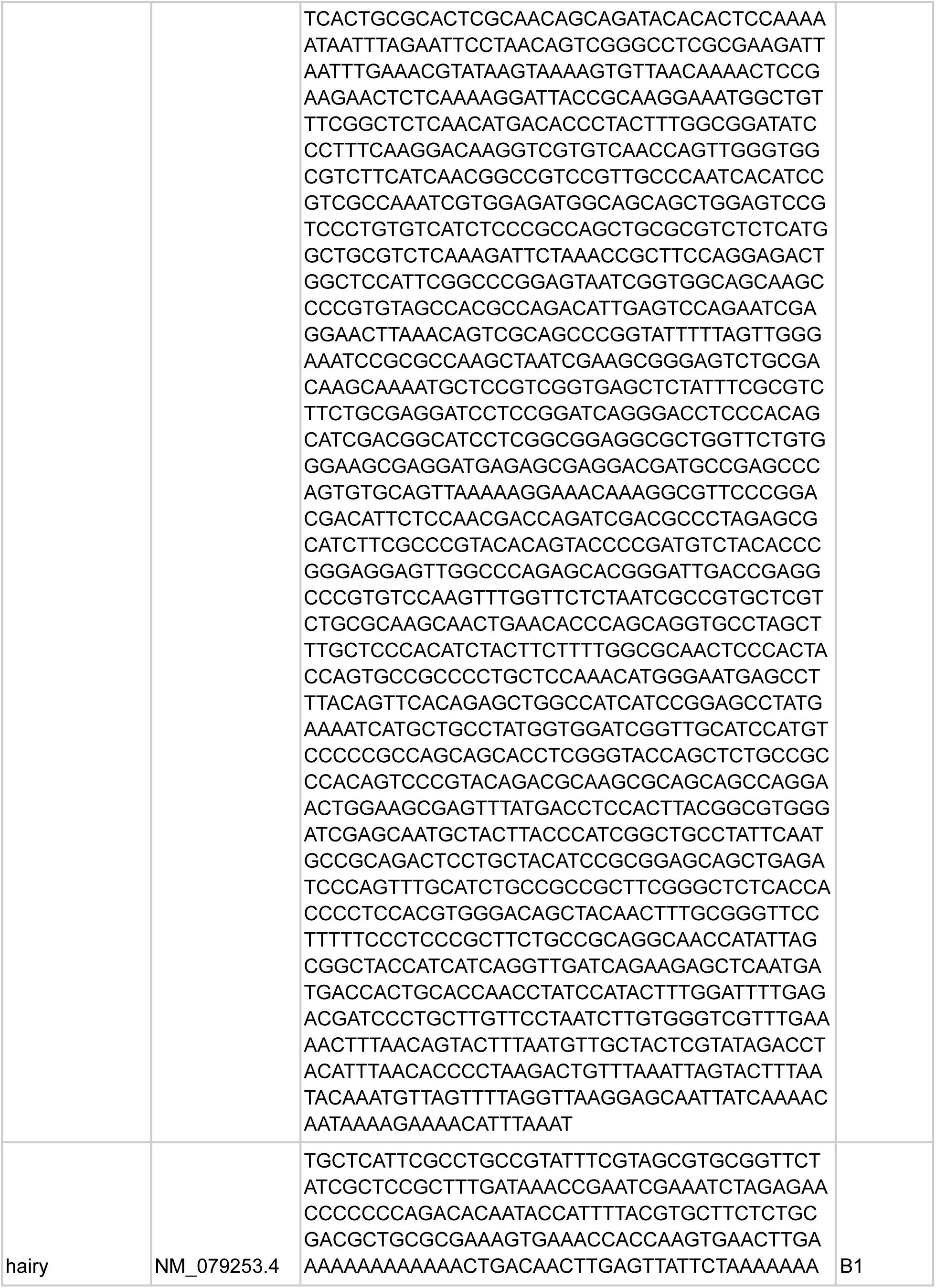
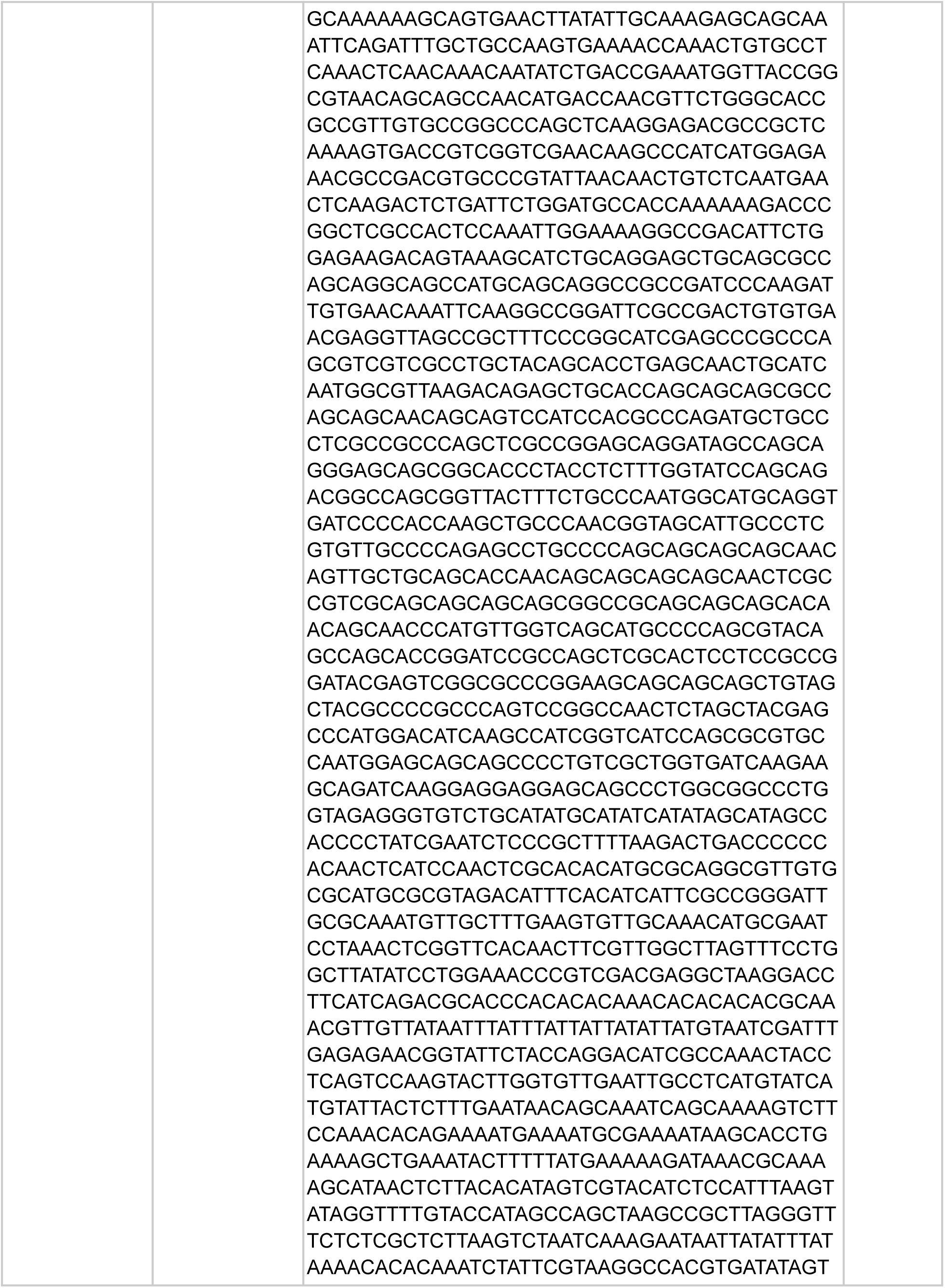
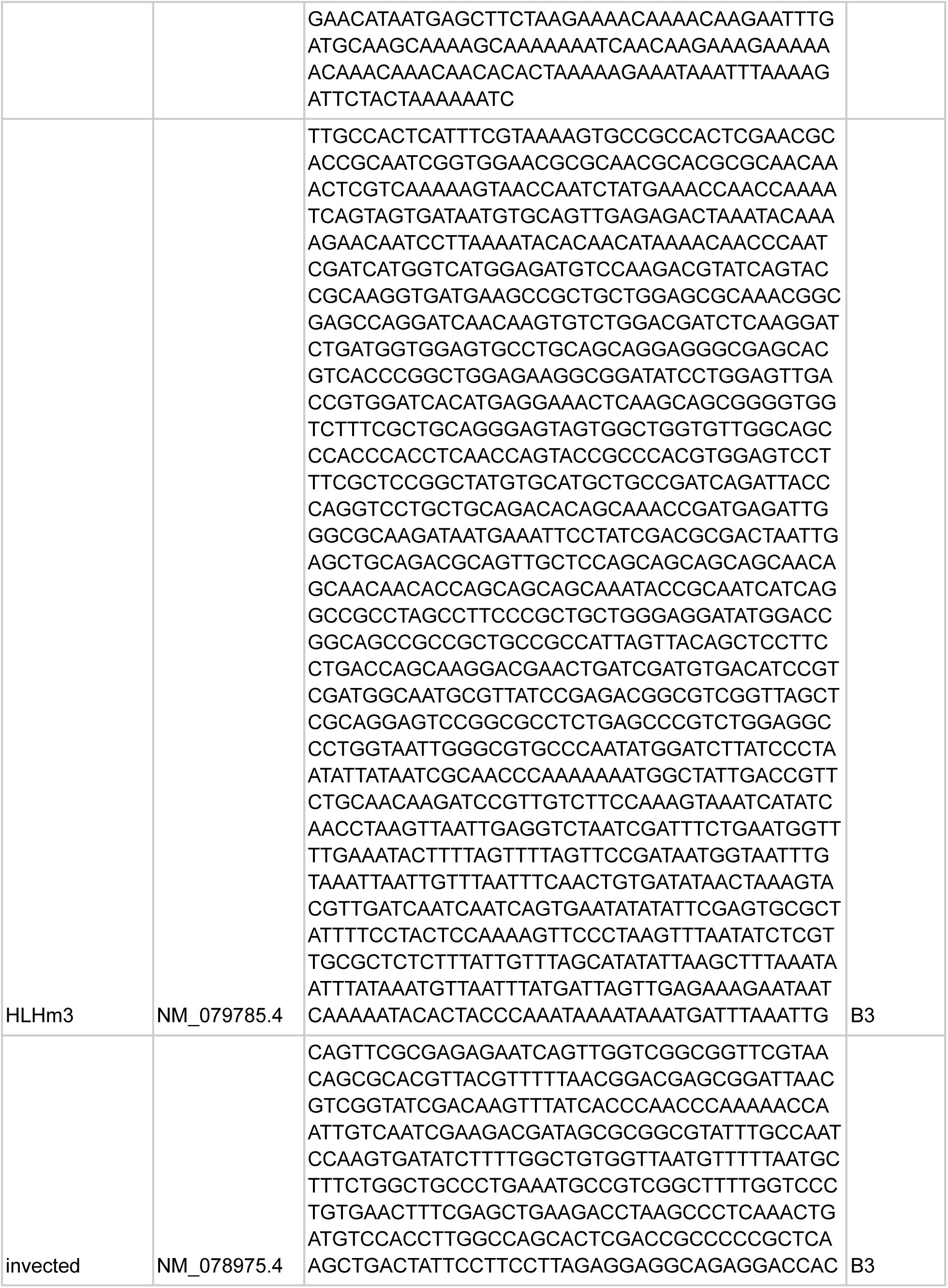
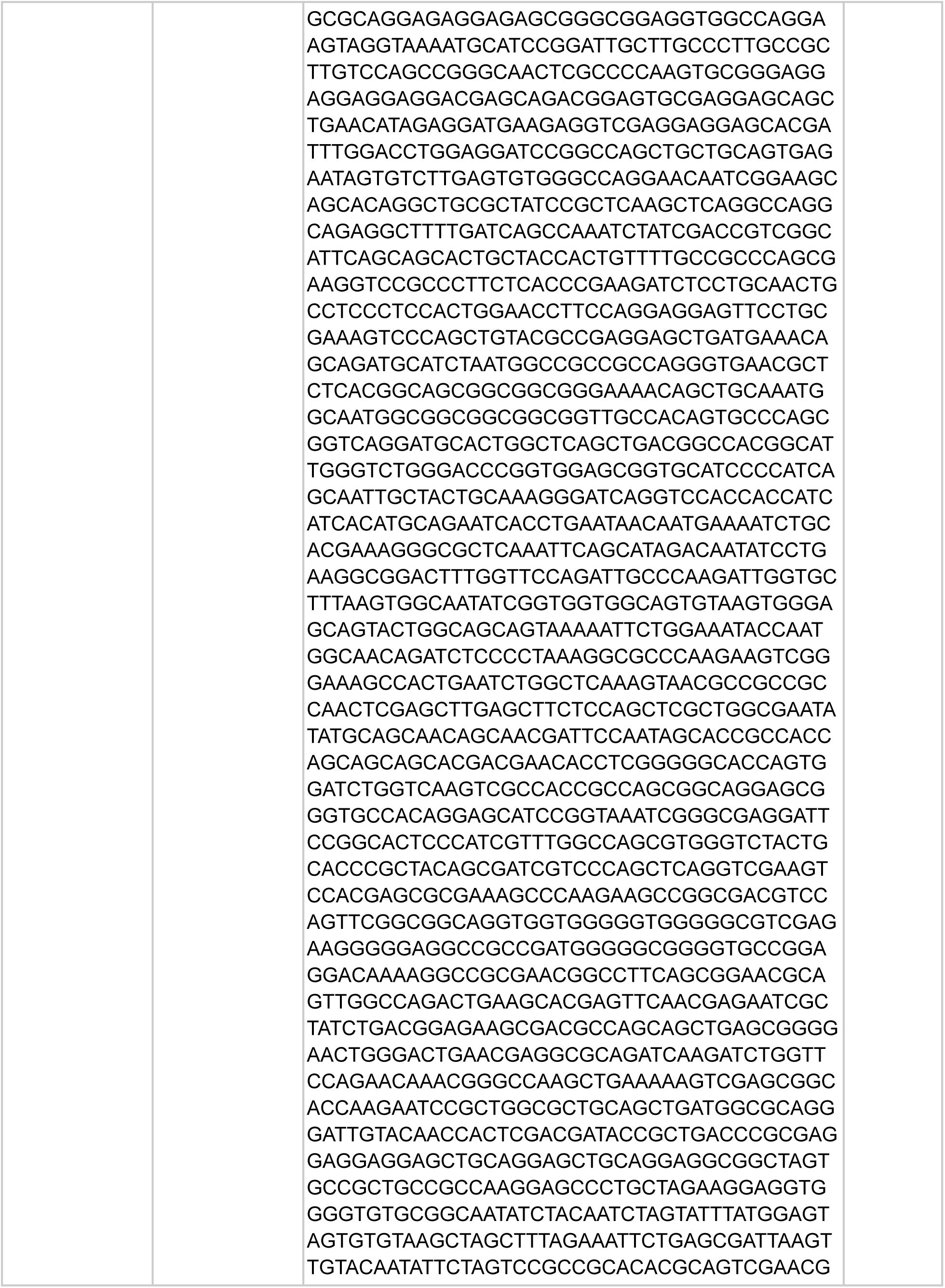
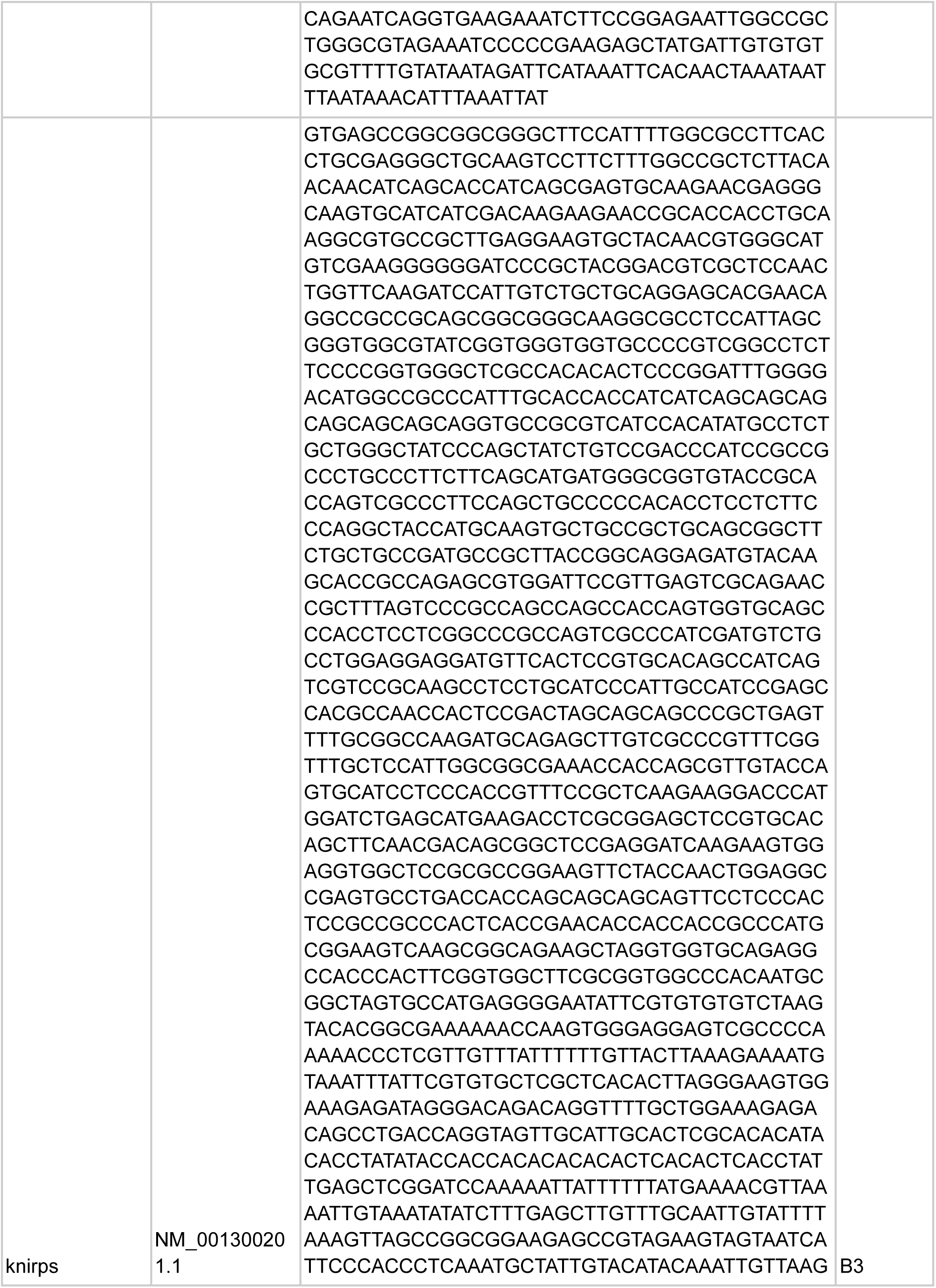
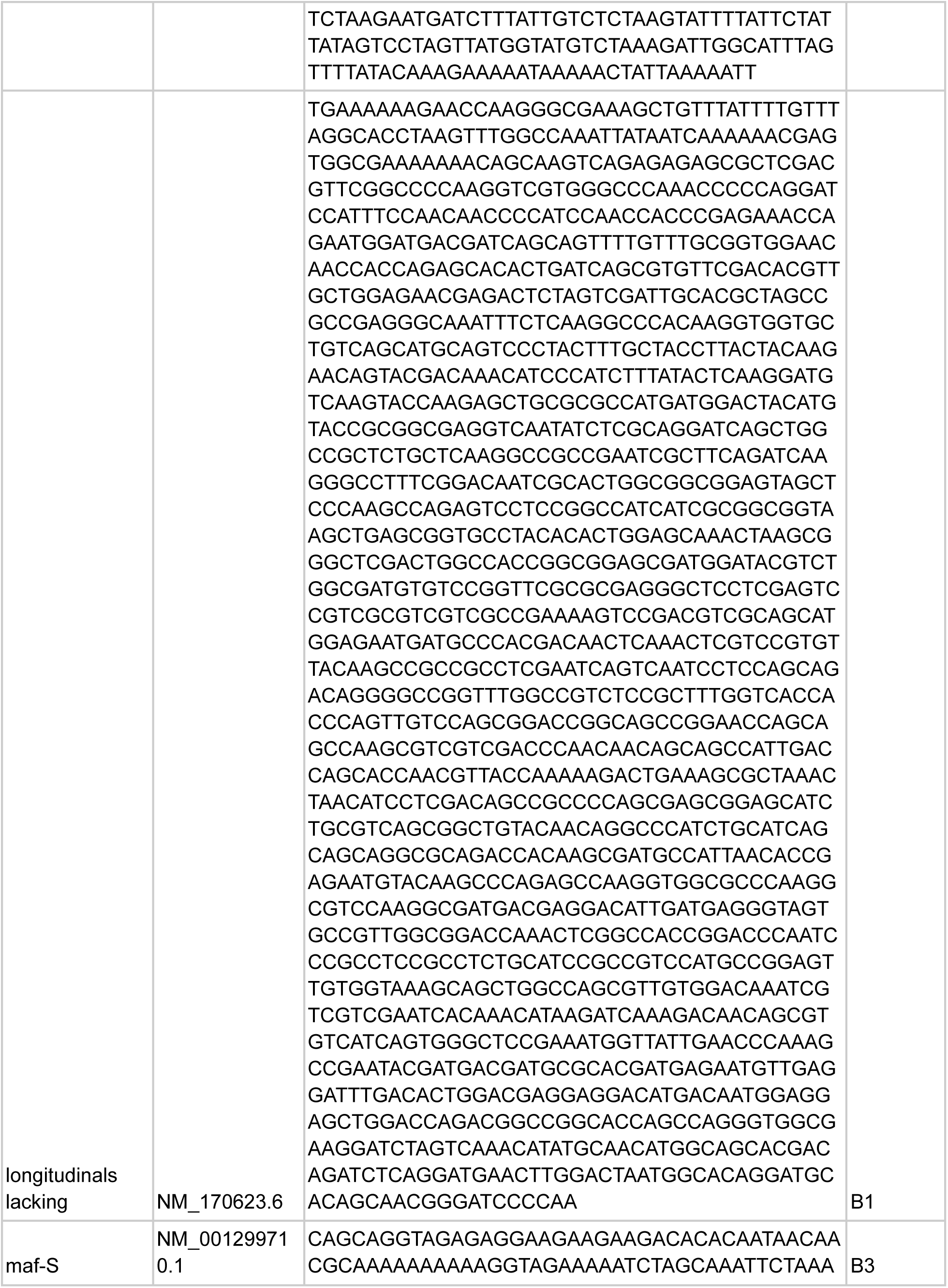
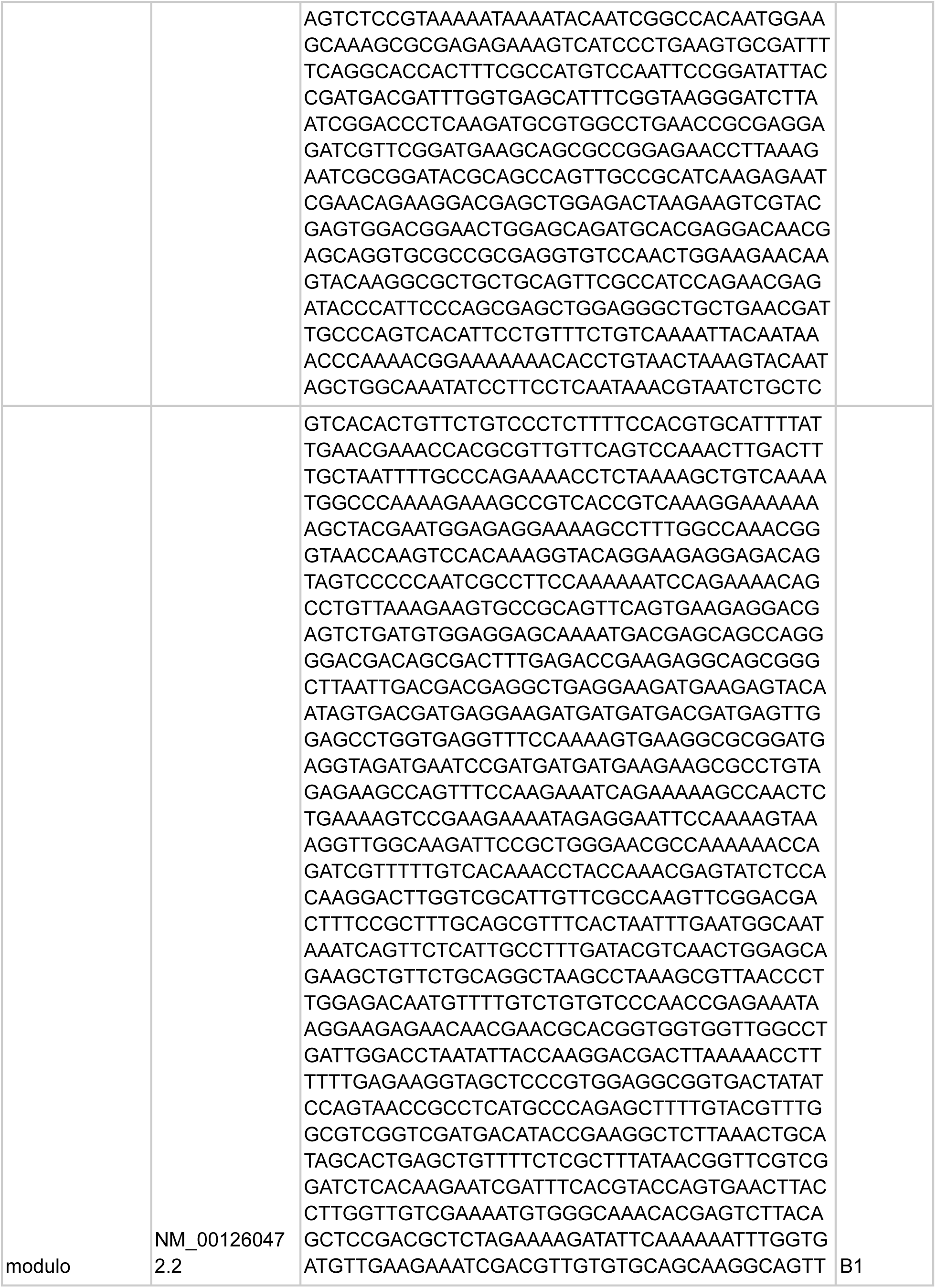
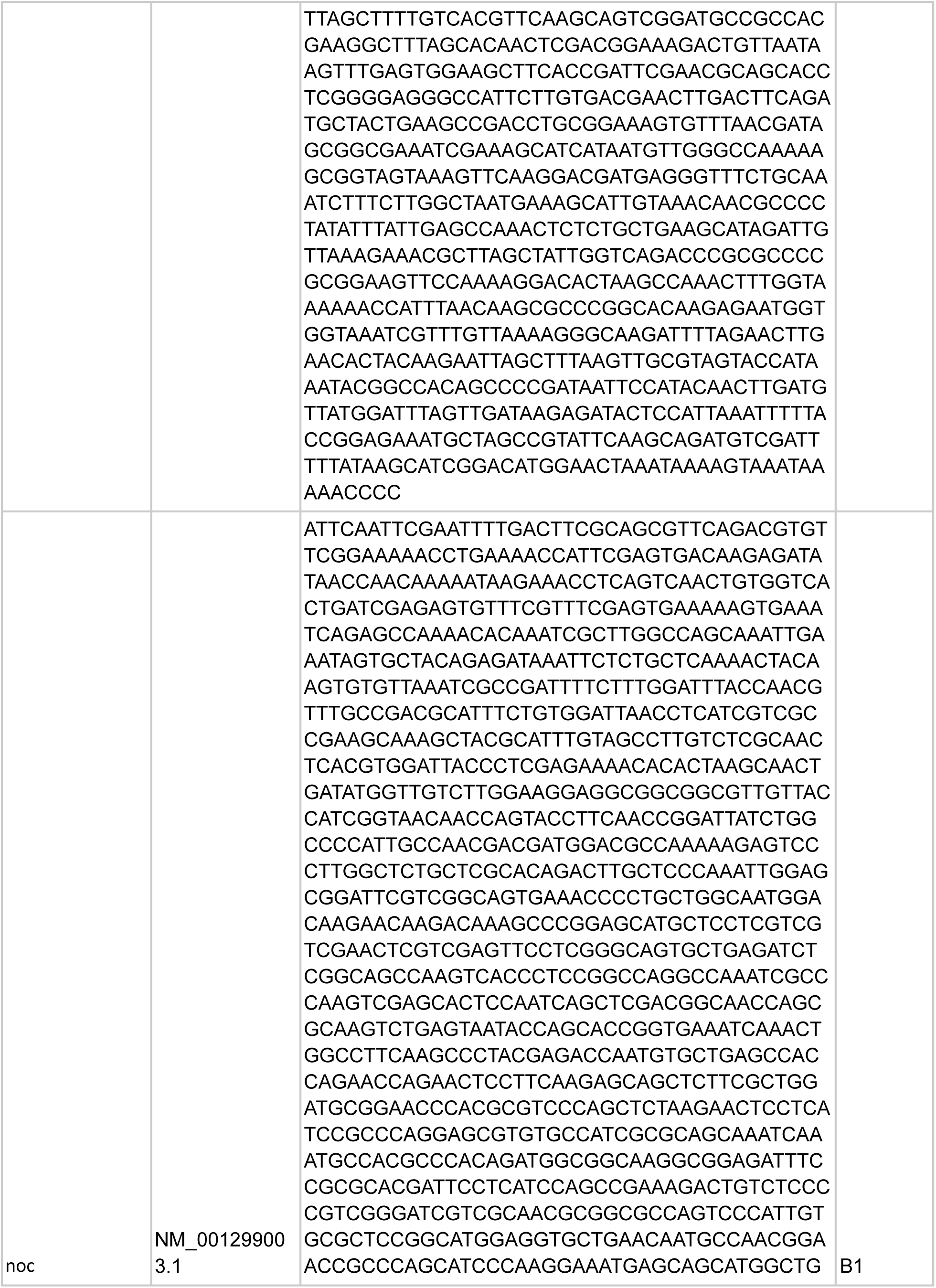
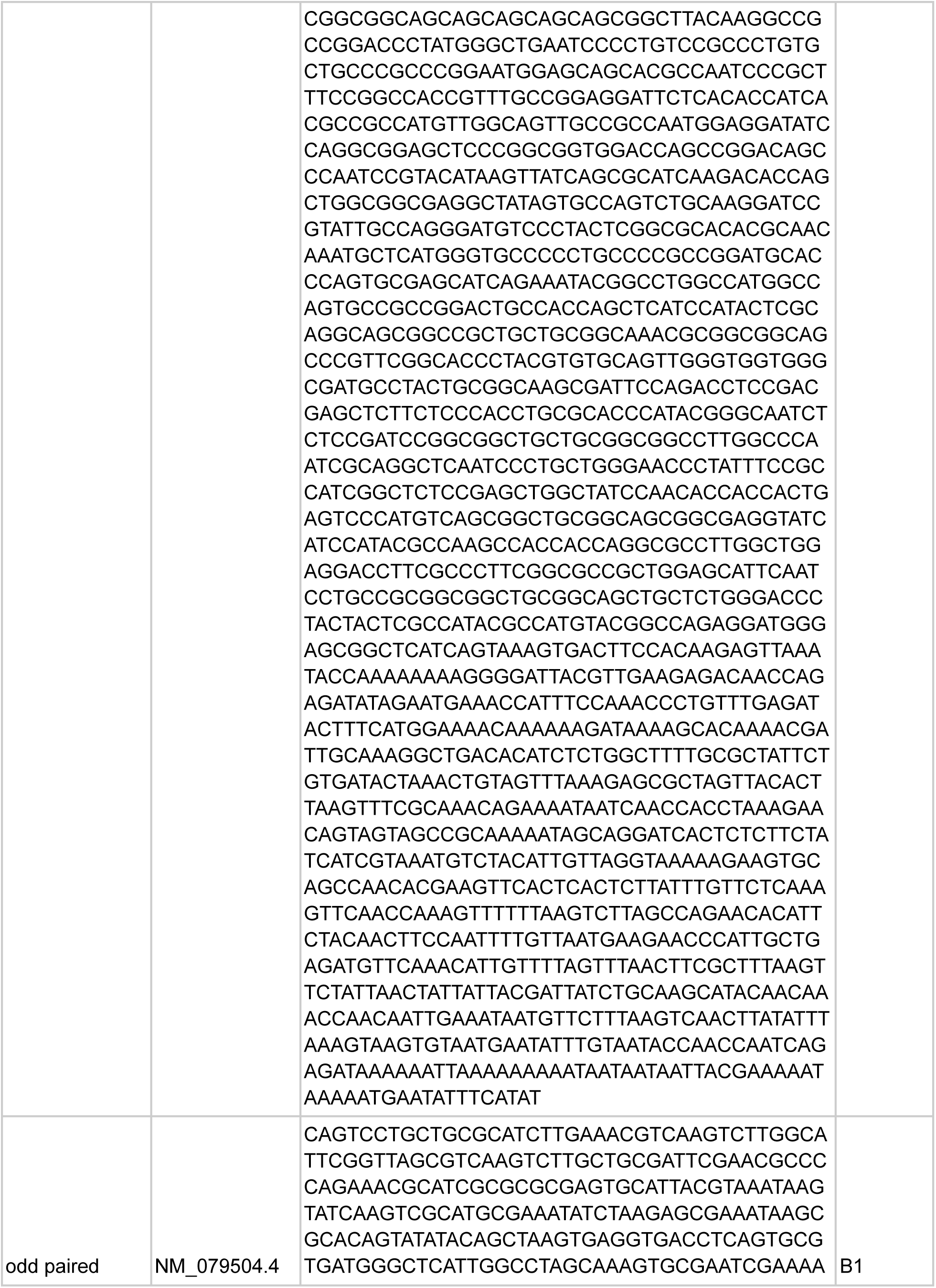
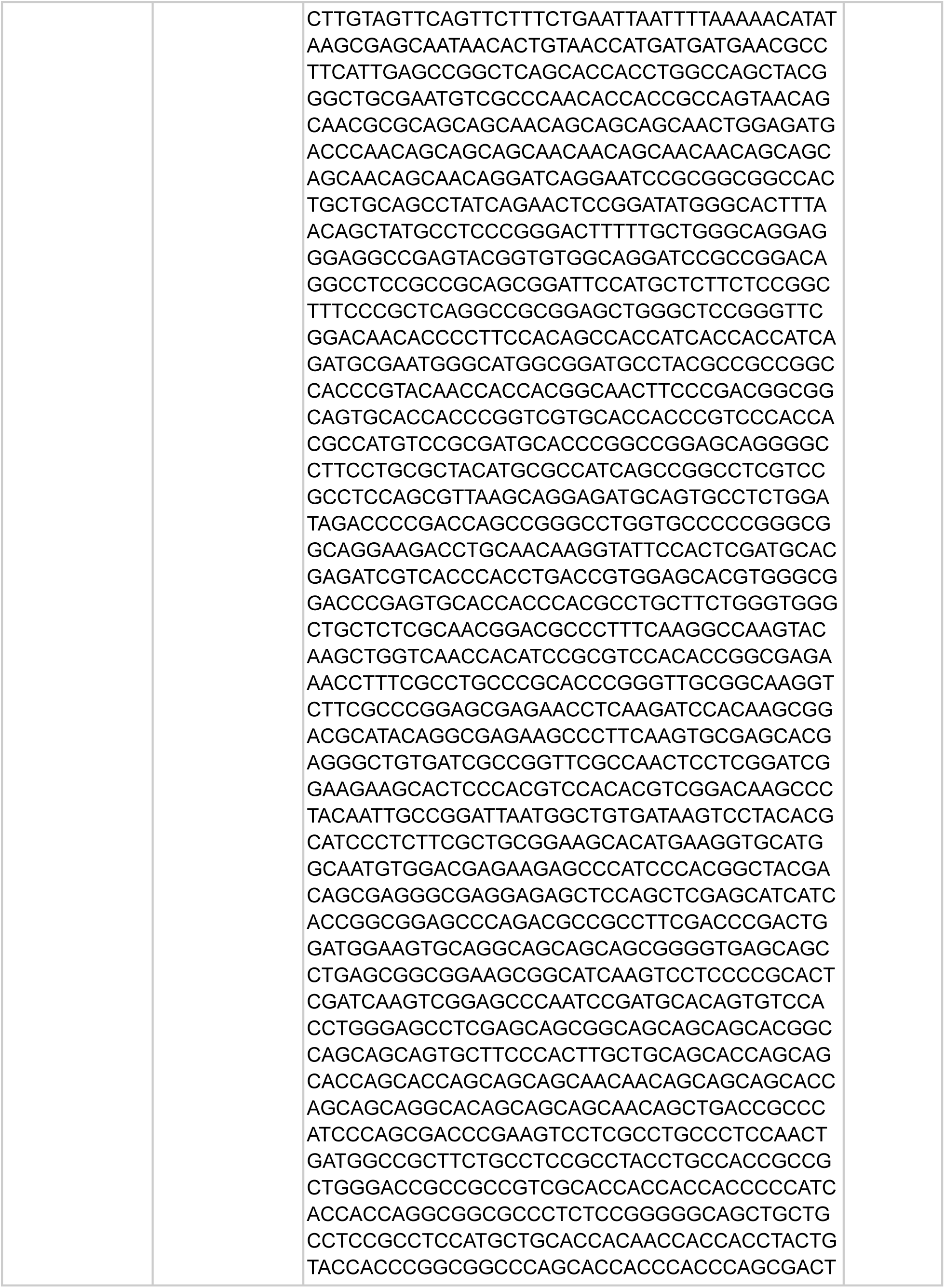
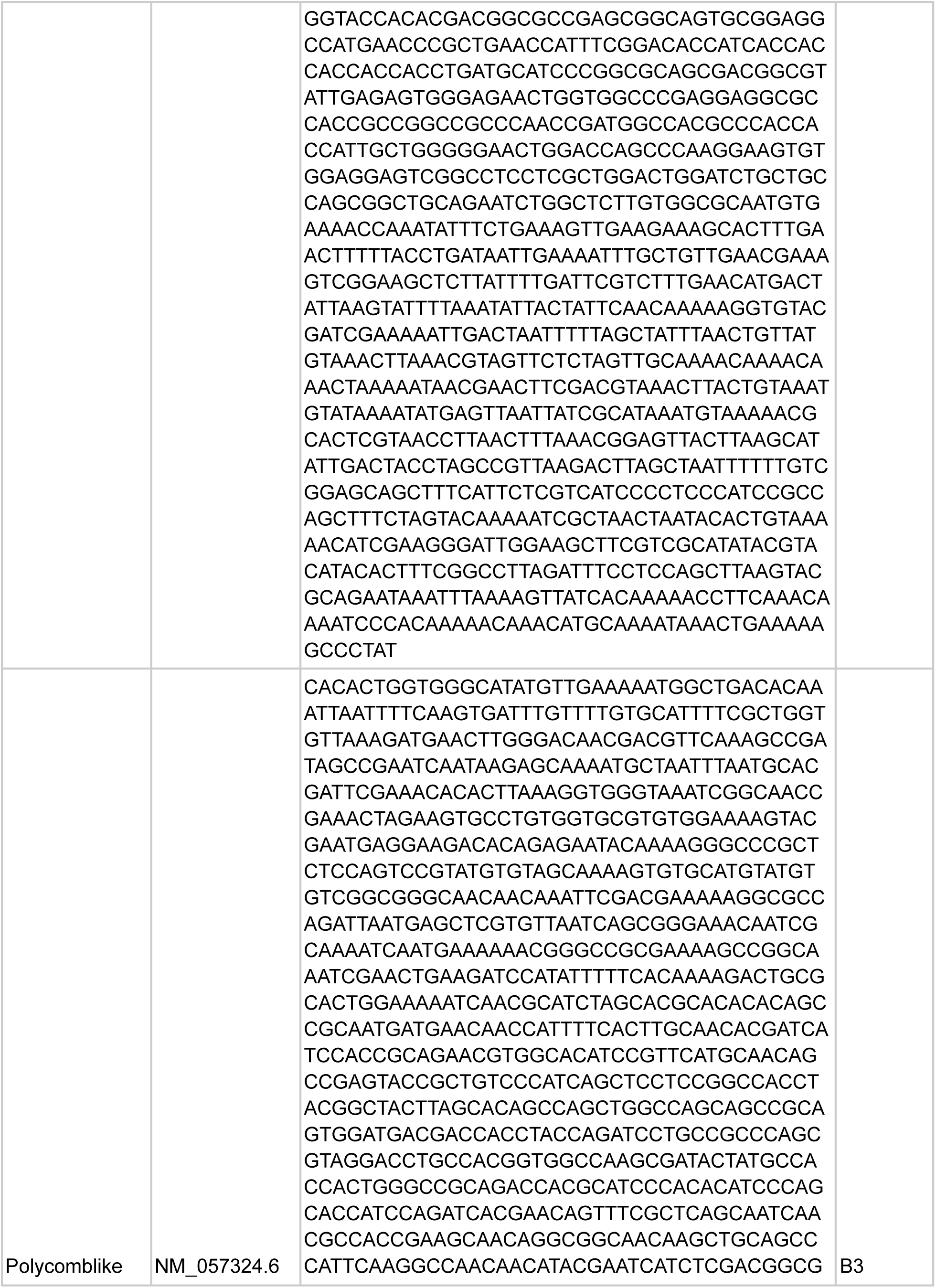
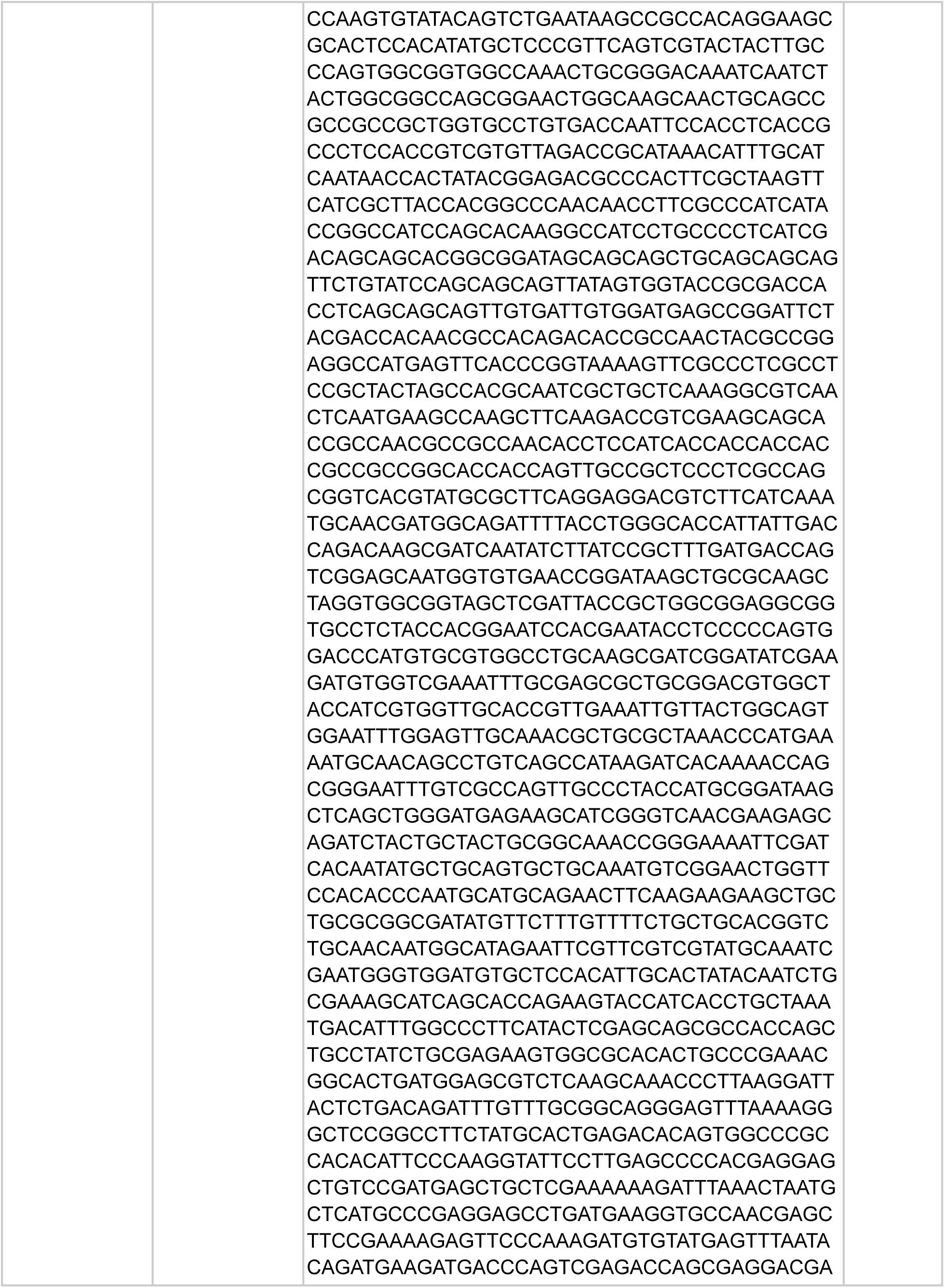
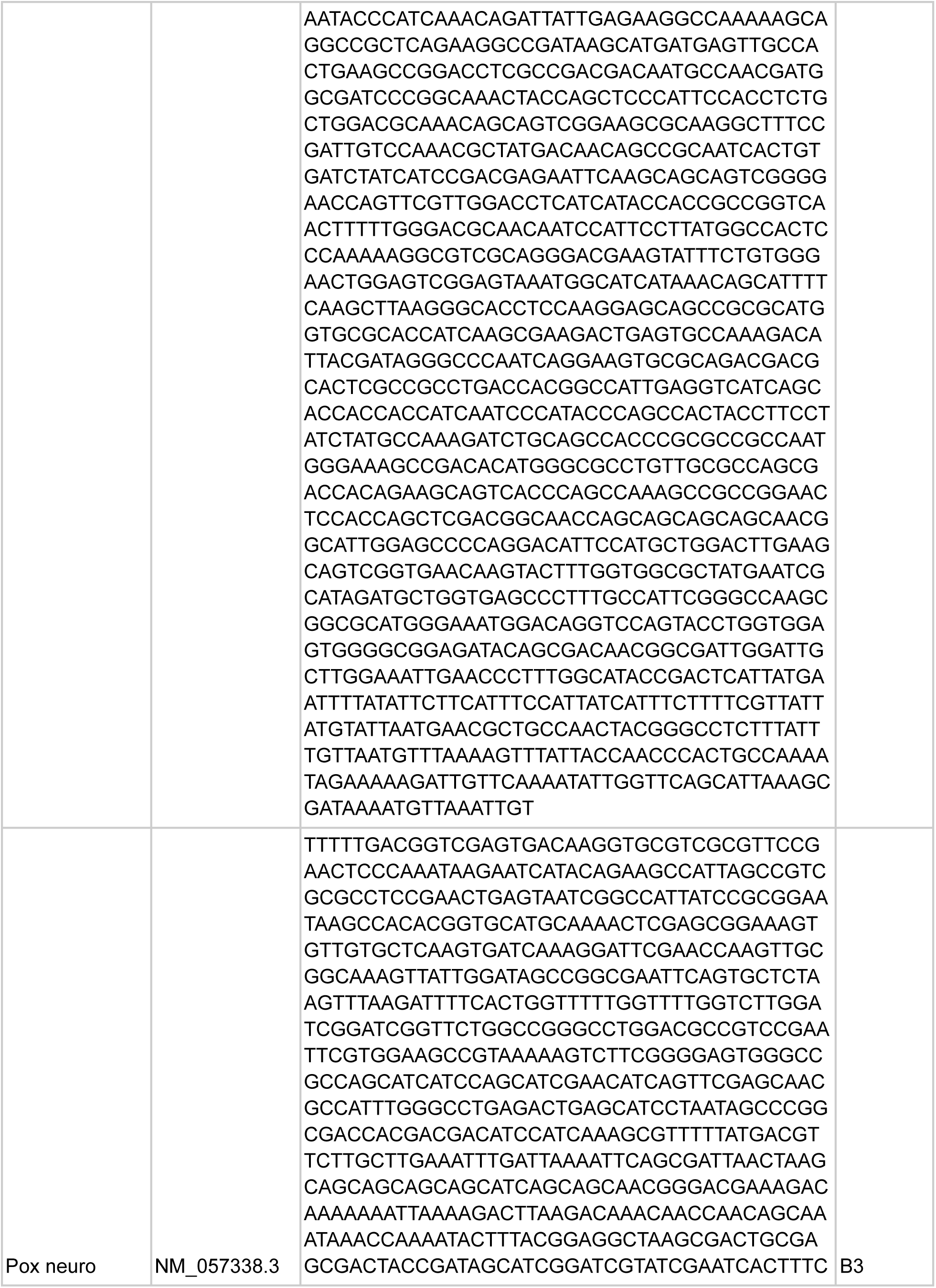
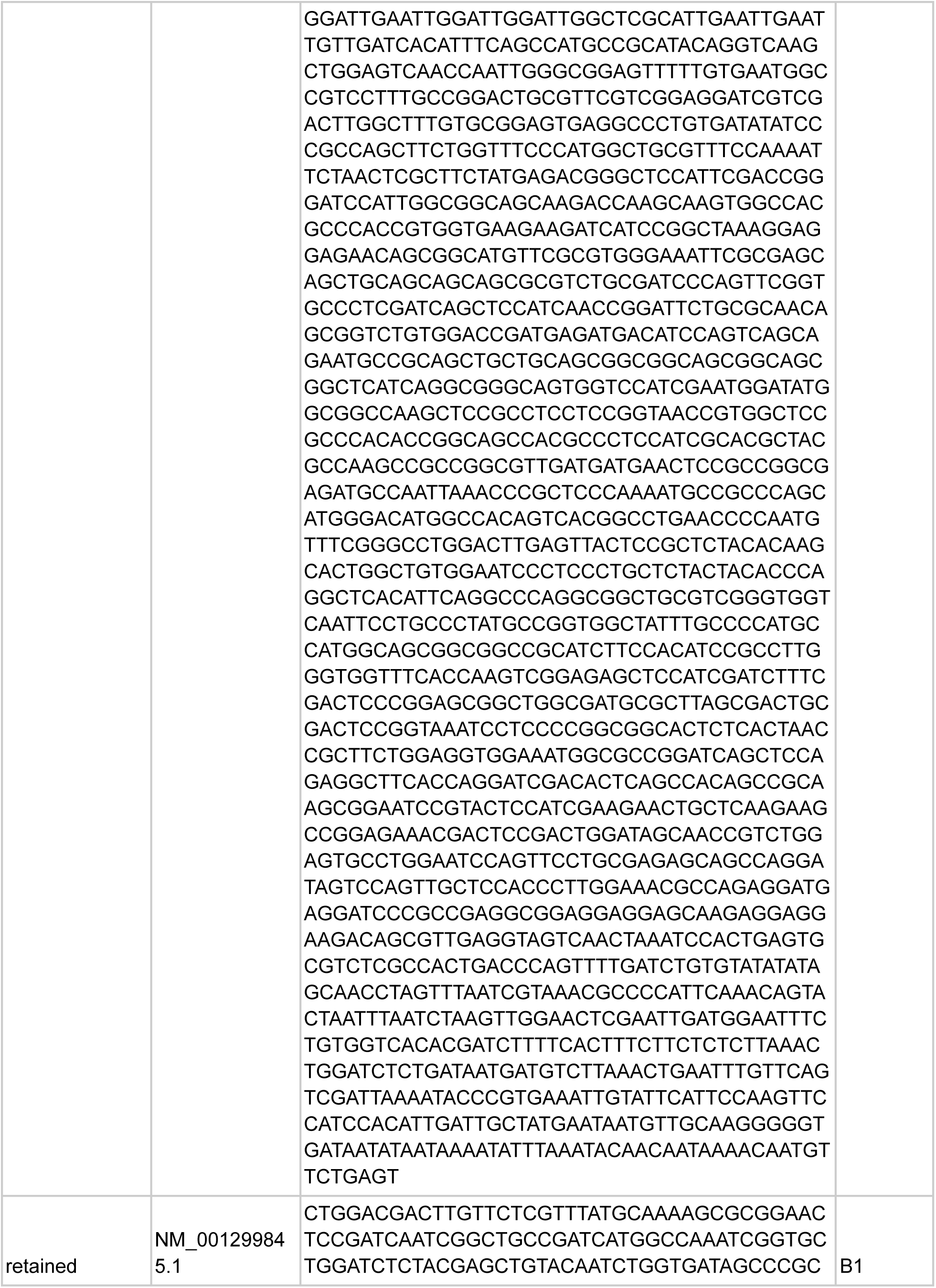
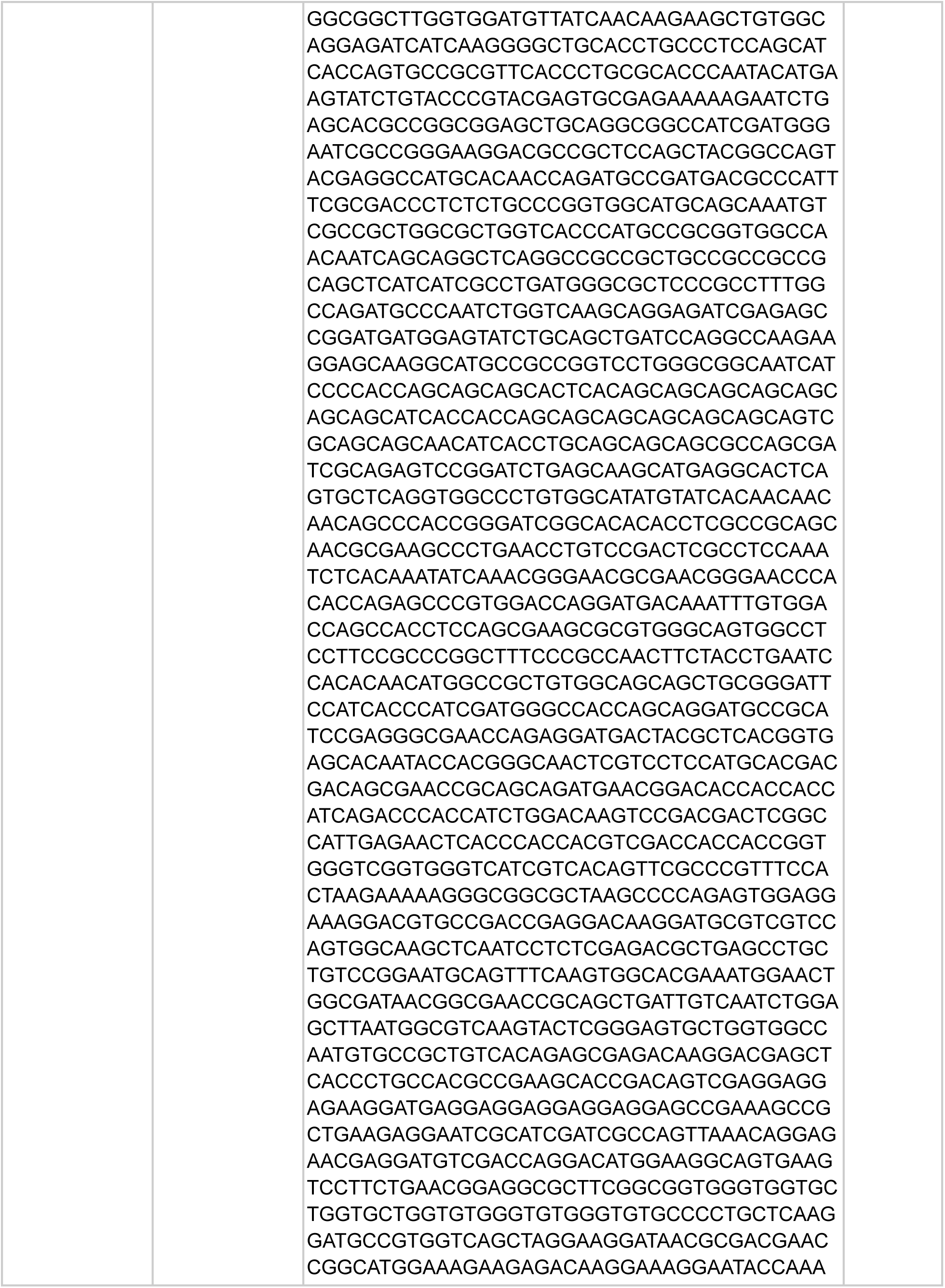
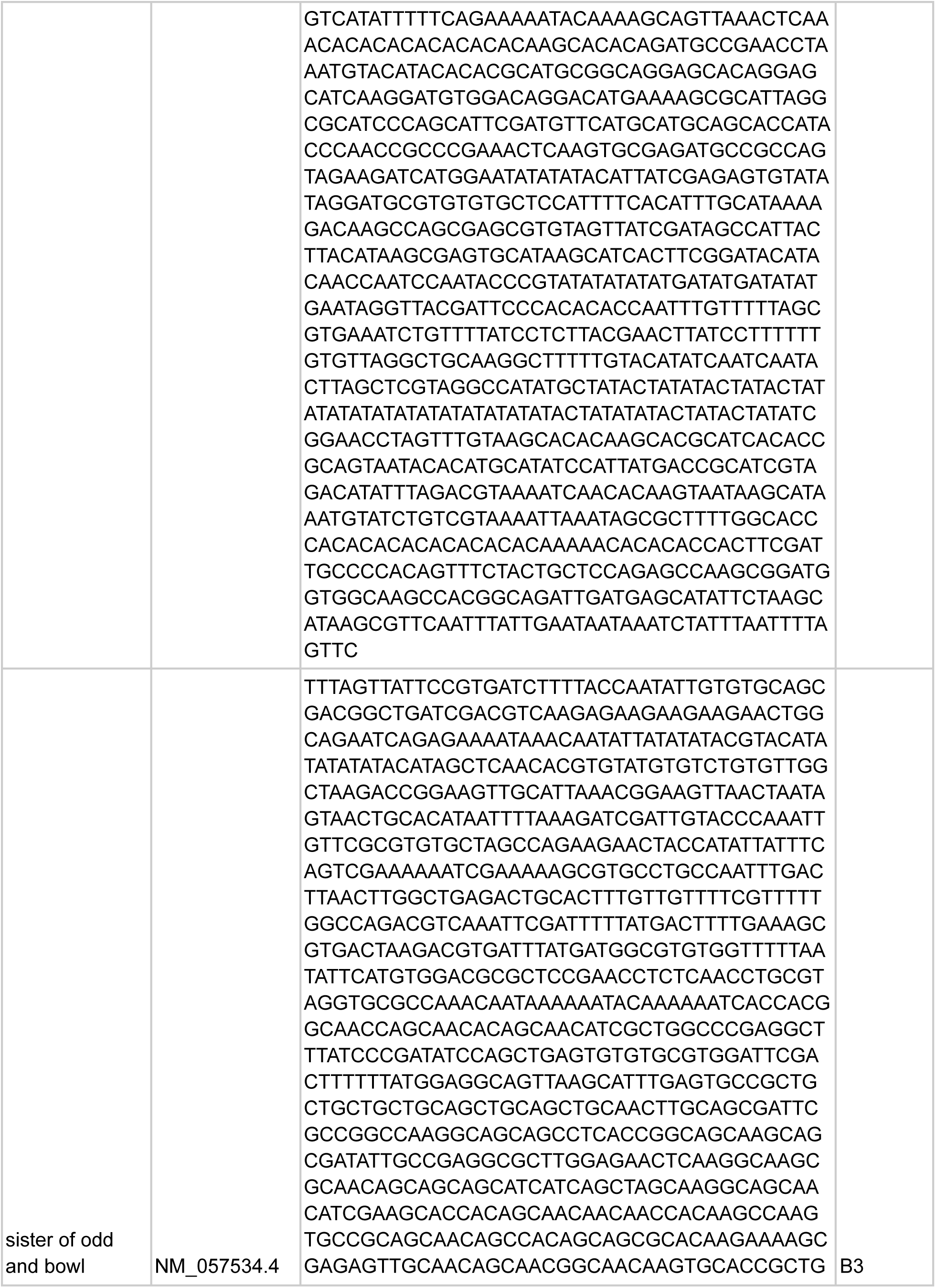
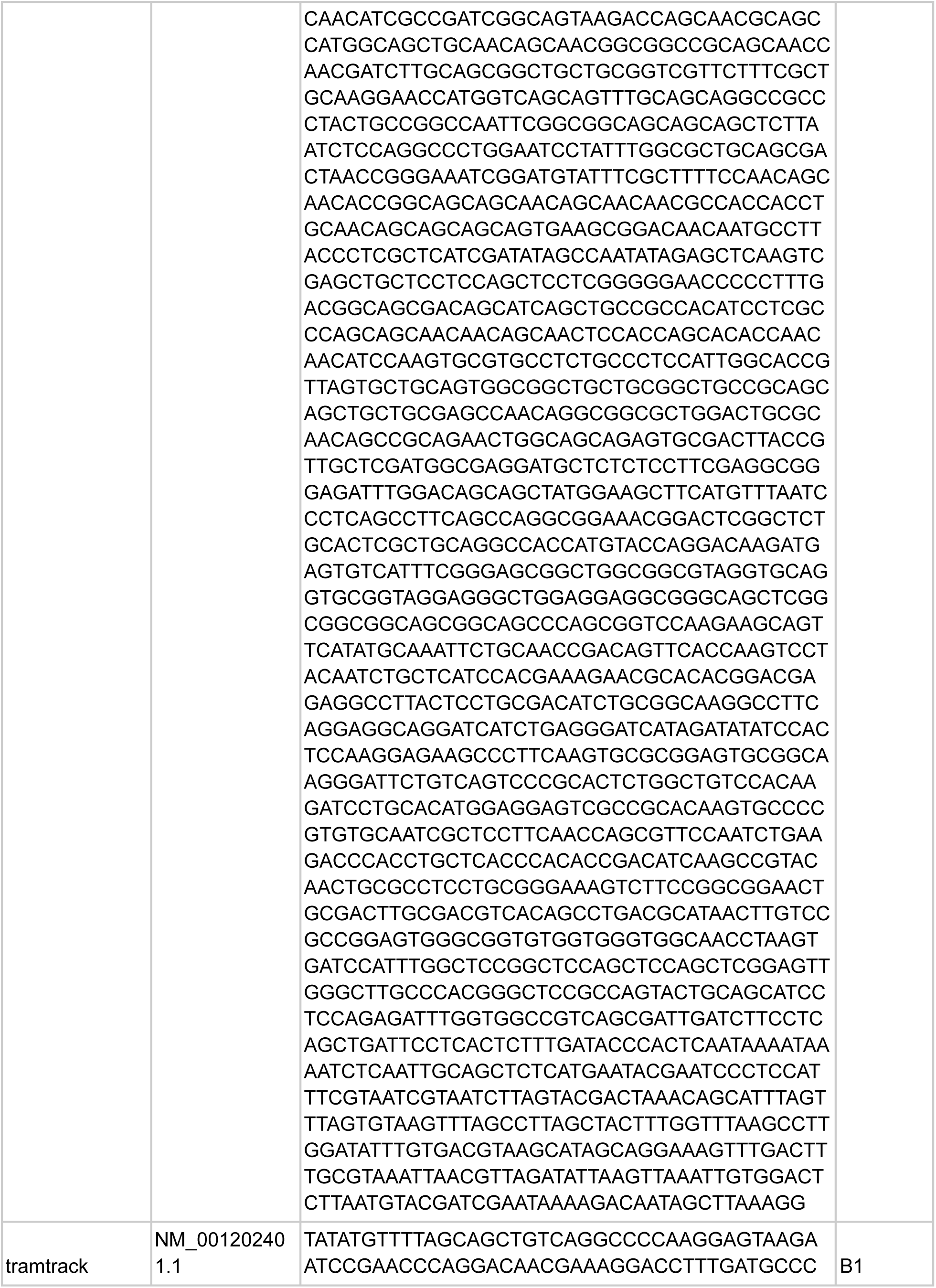
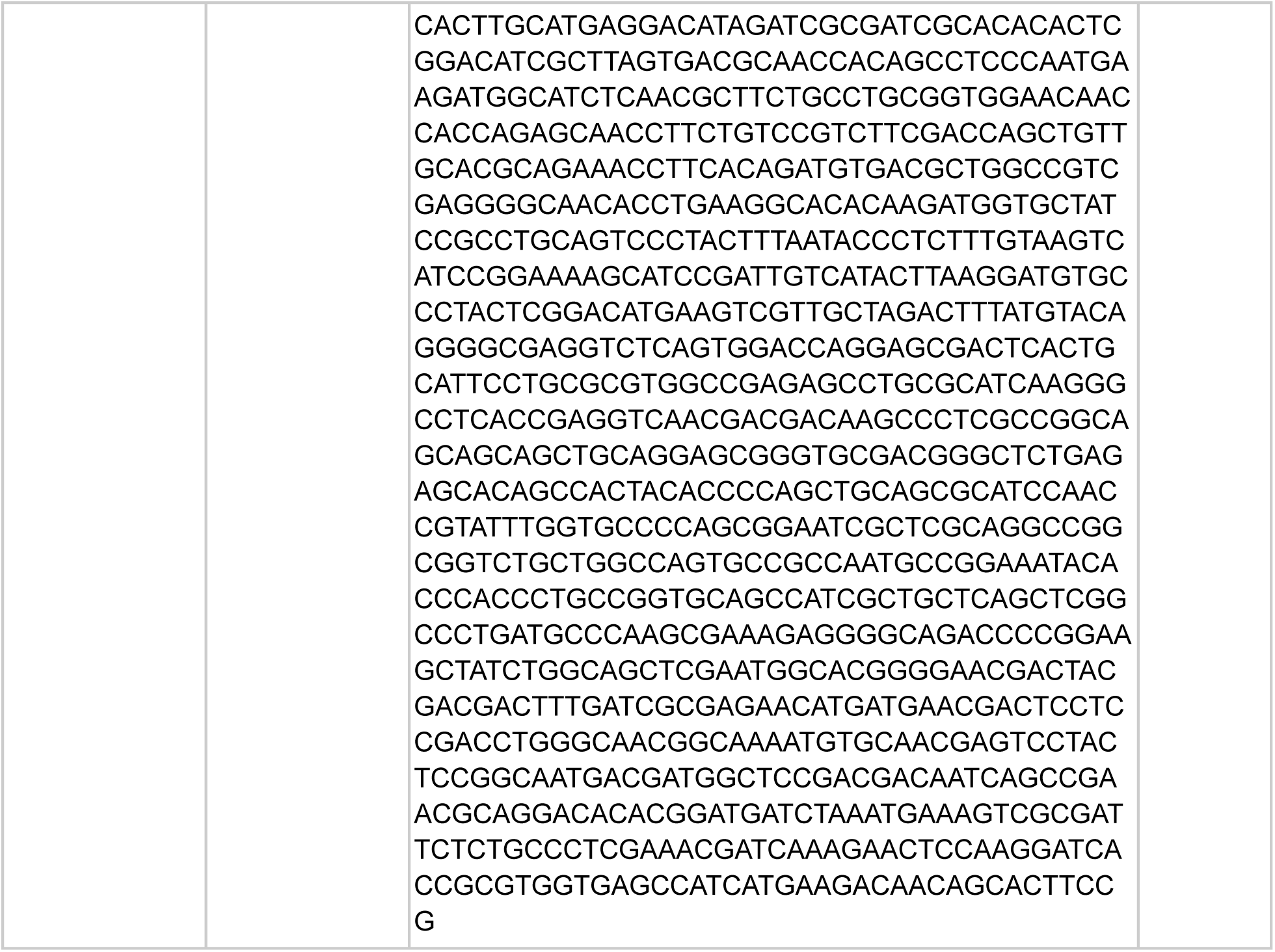
Summary of transcription factor expression signal in male genital structures. Image stacks of HCR signal were manually inspected, and expression levels were qualitatively assigned to one of three categories for each structure in the developing male genitalia: strongly expressed (2); weakly expressed (1); not expressed (0). Genes exhibiting ubiquitous expression patterns are also indicated, but these genes may exhibit varying expression levels across tissues.

